# A Stress-Adaptive Lipid Kinase Axis Defines Metabolic Vulnerabilities in Neuroendocrine Prostate Cancer

**DOI:** 10.64898/2026.06.15.732391

**Authors:** Yang Zheng, Caleb Cheng, Yizhi Cao, Gabriel Cruz, Yuping Zhang, Radha Paturu, Somnath Mahapatra, Jing Hu, Rahul Mannan, Hüseyin Karabürk, Rupam Bhattacharyya, Yitong Yin, Yi Zhao, Wenyan Liu, Xuhong Cao, Hui Xue, Chungen Li, Zhen Wang, Stephanie J. Miner, Ulka Vaishampayan, Vaibhav Sahai, Lois S. Weisman, Ke Ding, Costas A. Lyssiotis, Yuzhuo Wang, Yuanyuan Qiao, Arul M. Chinnaiyan

## Abstract

Neuroendocrine prostate cancer (NEPC) persists in a profoundly hypoxic microenvironment, yet the mechanisms enabling tumor adaptation to this metabolically challenging niche remain undefined. Here, we identify the lipid kinase PIKfyve as overexpressed in NEPC, functioning as a central node in a stress-lipid kinase axis that drives adaptation to persistent endoplasmic reticulum (ER) stress. Mechanistically, NEPC requires PIKfyve-mediated lysosomal degradation and lipid recycling to maintain metabolic homeostasis under hypoxia. PIKfyve inhibition disrupts lysosomal function, leading to ER stress accumulation and activation of a compensatory, sterol regulatory element-binding protein (SREBP)-dependent de novo lipogenesis program essential for NEPC survival. This stress-lipid axis creates a synthetic vulnerability between PIKfyve and fatty acid synthase (FASN), where dual inhibition synergistically amplifies ER stress, triggers the terminal unfolded protein response, and induces tumor cell death. These findings reveal a metabolic adaptation in NEPC and provide preclinical evidence that co-targeting PIKfyve and FASN can overcome hypoxia-associated stress adaptation.

## Introduction

Neuroendocrine prostate cancer (NEPC) is a highly aggressive form of prostate cancer marked by the loss of androgen receptor (AR) signaling and the acquisition of neuroendocrine features such as expression of synaptophysin (SYP) and neuron-specific enolase (NSE)^1^. While NEPC can arise *de novo*, it more frequently emerges following resistance to potent AR pathway inhibitors including enzalutamide and abiraterone^2^. Genetically, NEPC has been characterized by frequent loss of tumor suppressors *RB1*, *TP53*, and *PTEN*, and transcriptional upregulation of neuroendocrine lineage factors such as *FOXA2*, *SOX2*, *ASCL1*, and *ONECUT2*^3^. Despite advances in the molecular basis of the disease, therapeutic progress for NEPC has lagged. Platinum-based chemotherapy remains the mainstay of treatment, but responses are typically transient with heavy toxicity^4^. The limited efficacy of current targeted therapies underscores an urgent need to identify and exploit molecular vulnerabilities that are required for NEPC cell survival and treatment resistance.

Hypoxia is a hallmark of solid tumors and is known to disrupt protein folding within the endoplasmic reticulum (ER), leading to ER stress and activation of the unfolded protein response (UPR)^5^. The UPR consists of three canonical branches: PERK-ATF4, IRE1-XBP1, and ATF6, that orchestrate adaptive responses or apoptosis depending on the duration and severity of stress^6,7^. ER stress and UPR activation have been implicated in multiple tumor types, including breast cancer, pancreatic ductal adenocarcinoma (PDAC), and multiple myeloma^8–10^. NEPC tumors exhibit high levels of hypoxia, which contribute to lineage plasticity in part through transcription factors such as *ONECUT2*^11,12^. However, the presence and functional relevance of ER stress and UPR activation in NEPC remain undefined.

Autophagy and lysosomal degradation are critical cellular processes that enable the clearance of misfolded proteins and damaged organelles under stress conditions such as hypoxia and ER stress, while also contributing to lipid homeostasis^13–16^. In prostate cancer, autophagy plays context-dependent roles: it may suppress tumor development in localized disease by maintaining proteostasis, but in advanced stages such as castration-resistant prostate cancer (CRPC), it frequently becomes cytoprotective, promoting tumor cell survival under therapeutic and metabolic stress^17^. As lysosomal function is essential for autophagic flux, targeting lysosome-associated pathways has emerged as a strategy to disrupt this adaptive survival mechanism^18,19^. Although agents like chloroquine and hydroxychloroquine have been repurposed to block lysosomal acidification and autophagy, their clinical utility has been constrained by suboptimal pharmacokinetics^20^. These limitations highlight the need for more potent and selective approaches to target the autophagy-lysosome axis in cancer.

PIKfyve is a lipid kinase that synthesizes phosphatidylinositol 3,5-bisphosphate (PI(3,5)P_2_), a key phosphoinositide that regulates endo-lysosomal membrane dynamics. It plays an essential role in late endosome maturation, lysosomal membrane homeostasis, and autophagosome-lysosome fusion^21^. Pharmacologic or genetic disruption of PIKfyve impairs lysosomal trafficking and blocks autophagic flux^22^. Our previous work identified PIKfyve as the molecular target of ESK981, a potent small-molecule inhibitor, and demonstrated the therapeutic potential of PIKfyve inhibition in AR-positive (AR+) prostate cancer^22^. However, NEPC is a biologically and clinically distinct entity with unique genetic and metabolic features, and the role of PIKfyve in NEPC remains unexplored.

Here, we uncover a distinct ER stress phenotype in NEPC characterized by predominant activation of the PERK-ATF4 axis and heightened autophagic activity. We identify PIKfyve as a critical stress-integrating lipid kinase that supports NEPC survival by maintaining lysosomal function. Inhibition of PIKfyve shifts tumor cell reliance from lysosome-mediated autophagy to sterol regulatory element-binding protein (SREBP)-driven lipogenesis, a compensatory pathway that mitigates ER stress. Dual inhibition of PIKfyve and lipogenesis induces a synthetic vulnerability in NEPC by concurrently disabling two parallel stress-adaptive mechanisms. These findings position PIKfyve as a central node of metabolic and stress signaling in NEPC and reveal a targetable co-dependency that may overcome therapeutic resistance in this lethal disease subtype.

## Results

### PIKfyve is overexpressed and serves as a therapeutic target in NEPC

To elucidate the role of PIKfyve in NEPC, we analyzed the expression of both the protein and RNA in human tissue. Using a validated PIKfyve immunohistochemistry (IHC) antibody (**Figure S1A**) and *PIKFYVE* RNA-ISH probe^23^, we evaluated the expression of PIKfyve in anatomically distinct metastatic lesions from a rapid autopsy case of NEPC, termed WA76, including paraesophageal lymph node (LN), liver, central periaortic LN, common iliac LN, lower aortic LN, pelvic LNs (left and right sides), humerus bone marrow (BM), and femur BM. PIKfyve IHC and *PIKFYVE* RNA-ISH results indicated elevated *in situ* PIKfyve expression in NEPC tumor regions compared to adjacent benign tissues (**Figures 1A and S1B**). In addition, these findings were validated in a series of primary NEPC needle biopsies containing paired NEPC tumor and adjacent benign regions. Consistently, PIKfyve expression was increased in NEPC tumor regions across all cases (**Figures S1C and S1D**). Together, these findings provide the first *in situ* evidence that PIKfyve is upregulated in both primary and metastatic NEPC tumors, highlighting its clinical relevance and potential as a therapeutic target.

**Figure 1.**
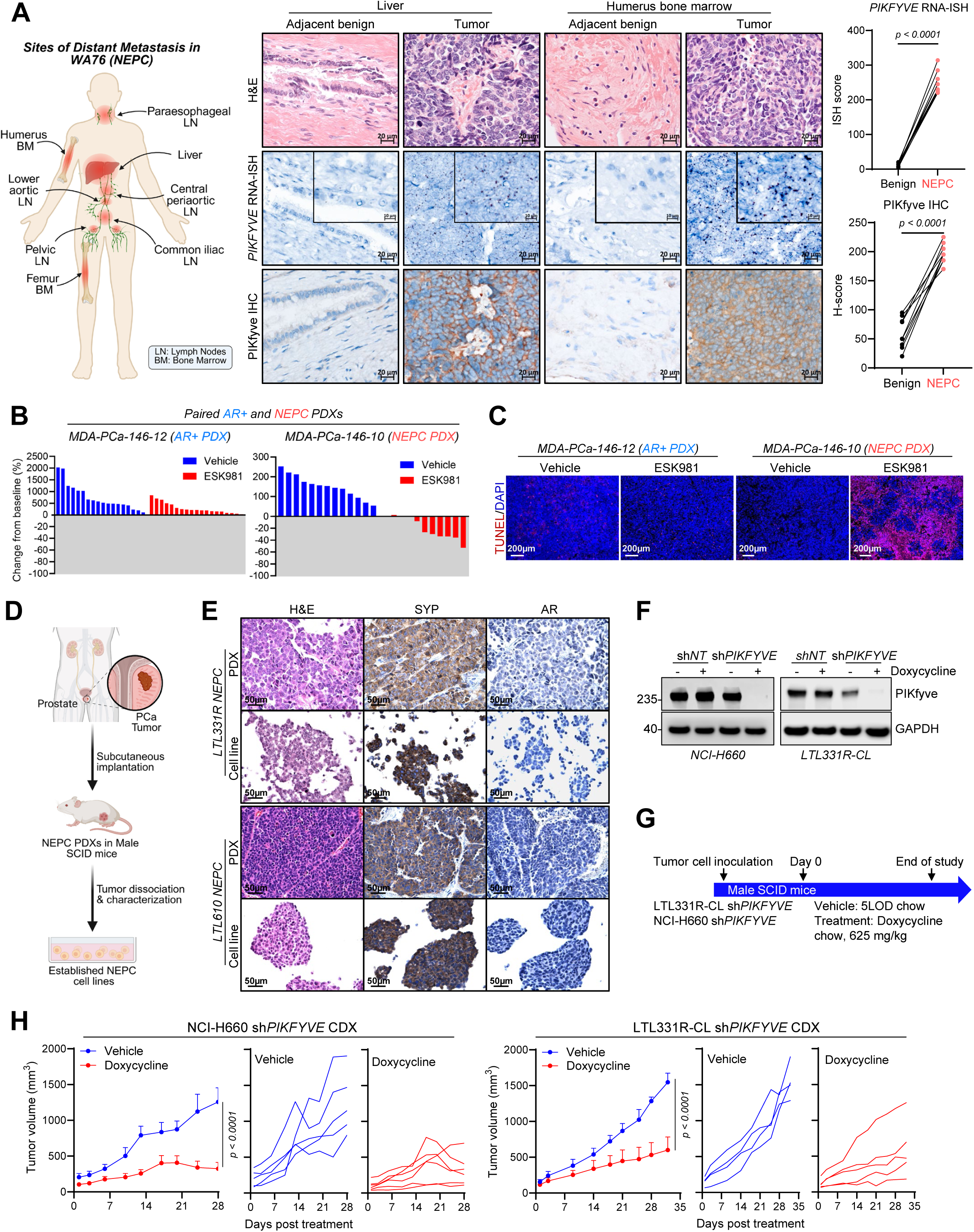
PIKfyve is overexpressed and serves as a therapeutic target in NEPC. A. Overview and analysis of metastatic PIKfyve expression in a rapid autopsy NEPC case (WA76). Left: schematic representation of nine anatomically distinct metastatic sites sampled from WA76, including paraesophageal lymph node (LN), liver, central periaortic LN, common iliac LN, right humerus bone marrow (BM), lower aortic LN, pelvic LNs (left and right), and right femur BM. Middle: representative histological and molecular images from two metastatic sites (liver and humerus bone marrow), showing hematoxylin and eosin (H&E) staining, RNA *in situ* hybridization (RNA-ISH), and immunohistochemistry (IHC) for PIKfyve in matched tumor and adjacent benign tissues. Right: quantification of *PIKFYVE* RNA-ISH signal (top) and PIKfyve IHC H-score (bottom) across all nine metastatic sites. Paired tumor and benign regions were scored separately. *P* values calculated by a paired student’s *t*-test. B. Waterfall plots showing individual tumor volume changes from baseline in vehicle or ESK981 treated group in a paired AR+ prostate cancer and NEPC patient derived xenograft (PDX) series, MDA-PCa-146-12 (AR+ PDX, vehicle *n* = 18, ESK981 *n* = 19) and MDA-PCa-146-10 (NEPC PDX, vehicle *n* = 14, ESK981 *n* = 10). Data adapted from Qiao et al., *Nature Cancer*, 2021^22^. C. Representative images of TUNEL staining showing *in situ* cell death post five days of either vehicle or ESK981 treatment in MDA-PCa-146-10 and MDA-PCa-146-12 tumors. D. Workflow for establishment of NEPC cell lines. PCa, prostate cancer. E. Representative images of H&E and IHC (AR and SYP) in LTL331R and LTL610 NEPC PDX and *ex vivo* cell lines (LTL331R-CL and LTL610-CL). F. Western blot of PIKfyve in the indicated NEPC cell lines with doxycycline-inducible sh*NT* and sh*PIKFYVE* with or without 1 µg/mL doxycycline treatment for 48 hours. G. Study design for NEPC sh*PIKFYVE* cell line-derived xenograft (CDX) model *in vivo*. H. Average and individual tumor growth curves of NEPC sh*PIKFYVE* CDXs treated with vehicle (NCI-H660 *n* = 5, LTL331R-CL *n* = 4) or doxycycline (NCI-H660 *n* = 6, LTL331R-CL *n* = 5) *in vivo*. Data presented as mean ± SEM*. P* values calculated using two-way ANOVA.

To investigate the differential dependency of PIKfyve in prostate cancer, we reanalyzed tumor responses to the PIKfyve inhibitor, ESK981, in a panel of prostate cancer models^22^. The waterfall plots indicated that NEPC (MDA-PCa-146-10 patient-derived xenograft (PDX)) and AR-negative (AR-) prostate cancer (DU145 cell line-derived xenograft (CDX)) exhibited tumor regression following PIKfyve inhibition by ESK981 treatment, whereas AR+ prostate cancer models (MDA-PCa-146-12 PDX and CRPC VCaP CDX) showed predominantly tumor stabilization (**Figures 1B and S2A**). Importantly, *in situ* cell death detected by TUNEL staining demonstrated that PIKfyve inhibition induced a robust cytotoxic effect in NEPC and AR-prostate tumors following five days of treatment, compared with a cytostatic effect in AR+ prostate tumors (**Figures 1C and S2B**). These findings indicate that PIKfyve is indispensable for NEPC tumor survival *in vivo*, and AR-independent prostate cancer has a higher reliance on PIKfyve than AR+ prostate cancer.

Recognizing the paucity of NEPC models *in vitro*, we established two novel NEPC cell lines, LTL331R-CL and LTL610-CL, derived from the corresponding patient-derived xenograft (PDX) models LTL331R and LTL610, respectively (**Figure 1D**). Histological characterization confirmed that these cell lines retained morphologies and neuroendocrine features resembling their parental tumors (**Figure 1E**). Both LTL331R-CL and LTL610-CL exhibited clustered growth in suspension and showed comparable doubling times to the established NEPC cell line NCI-H660 *in vitro* (**Figures S2C and S2D**). Short tandem repeat (STR) profiling confirmed the genetic identity of each cell line relative to its parental tumor, with stable allele patterns maintained across early to late passages (**Table S1**). Next, we generated stable cell lines of LTL331R-CL and NCI-H660 with doxycycline-inducible knockdown of *PIKFYVE* or non-targeting control (sh*PIKFYVE* or sh*NT*) (**Figure 1F**). Doxycycline-induced *PIKFYVE* knockdown impaired the tumor growth of xenografts derived from LTL331R-CL sh*PIKFYVE* and NCI-H660 sh*PIKFYVE* cells (**Figures 1G and 1H**), without affecting body weight (**Figure S2E**). Immunoblotting confirmed PIKfyve protein reduction in tumors after doxycycline treatment, albeit with residual protein detectable (**Figure S2F**). These findings demonstrate PIKfyve is an essential mediator of NEPC tumors and provide novel preclinical models to further investigate PIKfyve biology and therapeutic targeting.

### Pharmacological inhibition of PIKfyve exerts cytotoxicity in NEPC *in vivo*

PIKfyve catalyzes the phosphorylation of PI3P to generate PI(3,5)P_2_ and PI5P in various mammalian cell types, including mouse fibroblasts^21^, HEK293 cells, CHO cells^24^, and human PDAC cells^23^ (**Figure 2A**). To evaluate whether PIKfyve retains this canonical lipid kinase function in NEPC, we performed radioactive inositol labeling in NCI-H660 cells to trace phosphoinositide conversion following treatment with PIKfyve inhibitors ESK981 and apilimod^25^. Both inhibitors substantially decreased PI(3,5)P_2_ and PI5P levels while increasing upstream PI3P levels, without affecting PI4P and PI(4,5)P_2_ (**Figure 2B**), confirming that PIKfyve enzymatic activity was effectively inhibited by both ESK981 and apilimod in NEPC cells. Target engagement was further validated by cellular thermal shift assays (CETSA), which showed increased thermal stability of PIKfyve protein upon ESK981 (ΔTm = 0.5 °C) or apilimod (ΔTm = 1.2 °C) treatment in LTL331R-CL cells and additional prostate cancer models (LNCaP and DU145) (**Figures 2C and S3A**). These results confirmed the direct binding of ESK981 and apilimod to PIKfyve. Functionally, pharmacological inhibition of PIKfyve induced prominent cytoplasmic vacuolization in NEPC (NCI-H660, LTL331R-CL) and AR-prostate cancer cells (DU145, PC3) (**Figures 2D and S3B**), in line with prior reports^22,26^. PIKfyve inhibition disrupted autophagic flux, evidenced by the accumulation of lipidated LC3A/B and p62 proteins in NCI-H660, LTL331R-CL, and LTL610-CL cells (**Figure 2E**), increased autophagosome abundance (**Figure S3C**), and by impaired autophagic flux in GFP-LC3-RFP-LC3ΔG reporter DU145 and PC3 cells (**Figure S3D**). These results ascertain that both ESK981 and apilimod are specific PIKfyve inhibitors. Given the known pharmacokinetic limitations of apilimod^27^, we prioritized ESK981, a clinically applicable PIKfyve inhibitor, for subsequent *in vivo* evaluation.

**Figure 2.**
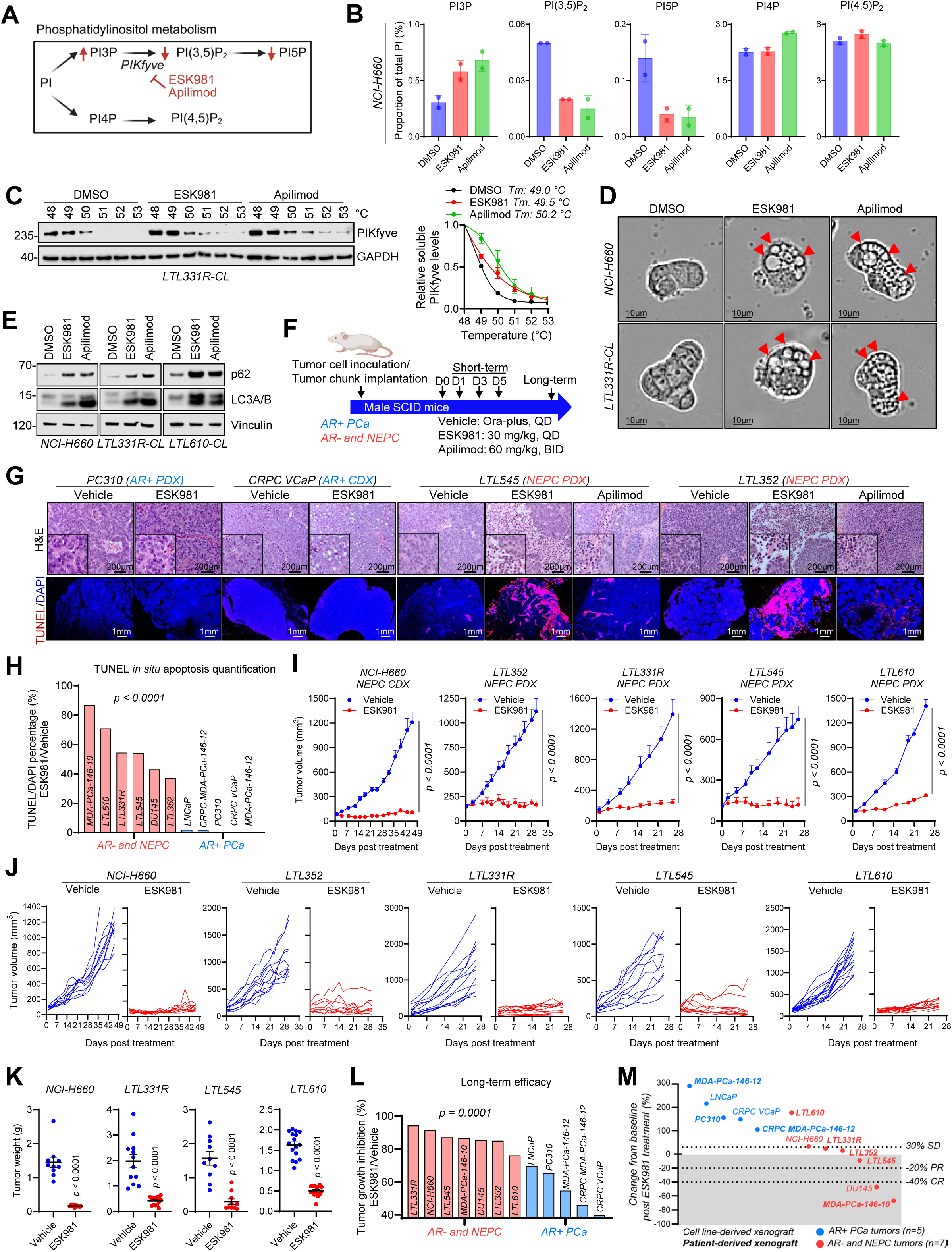
Pharmacological inhibition of PIKfyve exerts cytotoxicity in NEPC *in vivo*. A. Diagram of phosphatidylinositol metabolism showing the role of PIKfyve in the synthesis of PI(3,5)P_2_ and PI5P from PI3P. B. Quantification of lipid labeling analysis showing PI3P, PI(3,5)P_2_, PI5P, PI4P, and PI(4,5)P_2_ levels after DMSO, ESK981 (1 µM), or apilimod (1 µM) treatment for 24 hours in NCI-H660 cells. Data presented as mean ± SD. Experiments were independently repeated twice. C. Western blot of cellular thermal shift assay (CETSA) showing PIKfyve protein levels under increasing temperatures after treatment with DMSO, ESK981 (1 µM), or apilimod (1 µM) for 3 hours in LTL331R-CL cells (left); Quantification of relative soluble PIKfyve levels under the indicated treatments across temperatures. Data were normalized to GAPDH expression and to the 48 °C condition in each treatment. Tm was defined as the temperature at which 50% of soluble PIKfyve remained following heating. Data presented as mean ± SEM (right). D. Representative images showing vacuolization morphology induced by DMSO, ESK981 (1 µM), or apilimod (1 µM), for 24 hours in NCI-H660 and LTL331R-CL cells. Red arrows indicate vacuoles. E. Western blot of LC3A/B and p62 post DMSO, ESK981 (1 µM), or apilimod (1 µM) treatment for 24 hours in the indicated NEPC cell lines. F. Study design evaluating ESK981 monotherapy in various prostate cancer models *in vivo*. Treatments were given five days per week for vehicle and ESK981. Pharmacodynamic assessment includes one-day (D1), three-day (D3), and five-day (D5) treatment for vehicle, ESK981, and apilimod. QD, once per day. BID, 2 times per day. PCa, prostate cancer. G. Representative images of H&E and TUNEL *in situ* cell death staining in a panel of AR+ prostate and NEPC tumors post five days of vehicle, ESK981, or apilimod treatment. H. Quantification of TUNEL positive signals of the indicated AR+ or AR- and NEPC tumors post five days of vehicle, ESK981, or apilimod treatment. *P* value calculated using repeated-measures two-way ANOVA. I. Tumor growth curves of the indicated NEPC models treated with vehicle or ESK981 monotherapy for the indicated period. NCI-H660/LTL352/LTL545: vehicle *n* = 10, ESK981 *n* = 10; LTL331R: vehicle *n* = 12, ESK981 *n* = 14; LTL610: vehicle *n* = 16, ESK981 *n* = 16. Data presented as mean ± SEM*. P* value calculated using a two-tailed unpaired *t*-test with Welch’s correction. J. Individual tumor growth curves of NEPC tumors receiving either vehicle or ESK981 monotherapy for the indicated period. K. Tumor weight comparisons between vehicle and ESK981 treated groups in the indicated NEPC models. Data presented as mean ± SEM*. P* values calculated using a two-tailed unpaired *t*-test with Welch’s correction. L. Quantification of percent tumor growth inhibition for ESK981 monotherapy in AR+ or AR- and NEPC tumors. *P* value calculated using a two-tailed unpaired *t*-test. M. Tumor volume changes from baseline comparing ESK981 treatment to vehicle in multiple prostate cancer tumor models according to the RECIST criteria, where complete response (CR), average response ≤ −40 %; partial response (PR), > −40 % to ≤ −20 %; and stable disease (SD), > −20 % to < +30 %.

A series of *in vivo* experiments were designed to comprehensively examine the anti-tumor efficacy of PIKfyve inhibitors across a panel of prostate cancer PDX and CDX models (**Figure 2F**). Pharmacodynamic evaluation was performed on tumors receiving short-term treatment with vehicle, ESK981, or apilimod. Immunoblot analysis revealed a time-dependent accumulation of lipidated LC3A/B after 1, 3, and 5 days of ESK981 treatment (**Figure S3E**), consistent with disruption of PIKfyve-regulated lysosomal function *in vivo*. Extensive tumor necrosis and apoptosis were only observed in NEPC tumors compared to AR+ prostate cancer evidenced by H&E and TUNEL staining (**Figures 2G and 2H**). Apilimod exhibited weaker effects compared to ESK981, consistent with its *in vivo* instability (**Figure 2G**). The increased apoptotic response was corroborated by elevated c-PARP levels in NEPC tumors (**Figure S3F**). Additionally, time-course analyses showed a progressive enhancement of cell death with ESK981, which outperformed apilimod (**Figures S3G-I**). Importantly, acute cytotoxicity triggered by ESK981 translated into durable anti-tumor effects in five NEPC PDX and CDX models. ESK981 treatment led to marked suppression of tumor growth across models (**Figures 2I and 2J**), significant reduction in tumor weights (**Figures 2K and S3J**), and body weight profiles within limits (**Figure S3K**). In summary, tumor growth inhibition by ESK981 was over 1.5-fold greater in NEPC compared to AR+ prostate cancer models (**Figure 2L**), and waterfall plot analysis demonstrated greater responses in AR- and NEPC tumors, with some NEPC models achieving complete responses by RECIST criteria (**Figure 2M**). Consistent with prior reports^22,28,29^, PIKfyve inhibition modestly enhanced IFNγ-induced *CXCL10* expression and increased MHC class I levels in NEPC cells (**Figure S3L and S3M**), suggesting a potential link between PIKfyve inhibition and tumor cell-intrinsic immune signaling. Collectively, these findings show that PIKfyve is an essential regulator of NEPC tumor survival, and the availability of a phase II PIKfyve inhibitor, ESK981, provides immediate translational potential to target this vulnerability in NEPC patients^30,31^.

### NEPC tumors exhibit elevated baseline ER stress and autophagy

To investigate the high dependency of NEPC on PIKfyve, we performed RNA sequencing (RNA-seq) analysis on tumors derived from five NEPC and four AR+ prostate cancer xenograft models (**Figure 3A**). Pathway enrichment analyses confirmed robust enrichment of NEPC-specific signatures^32^ in NEPC tumors, while AR signaling signatures^32^ predominated in AR+ prostate tumors (**Figure 3B**). Gene set enrichment analysis (GSEA) validated the distinction, revealing upregulation of NEPC signatures and downregulation of AR pathways in NEPC tumors (**Figures S4A and S4B**). Consistent with prior reports^11,12^, hypoxia-related pathways were elevated in NEPC compared to AR+ tumors (**Figure S4C**). This was corroborated by increased HIF-1α expression observed with HIF-1α IHC, enhanced Hypoxyprobe signal observed by pimonidazole (PIMO) IHC staining (**Figure S4D**), and upregulation of *ONECUT2* (**Figure S4E**), a master transcription factor that cooperates with hypoxia signaling^12^.

**Figure 3.**
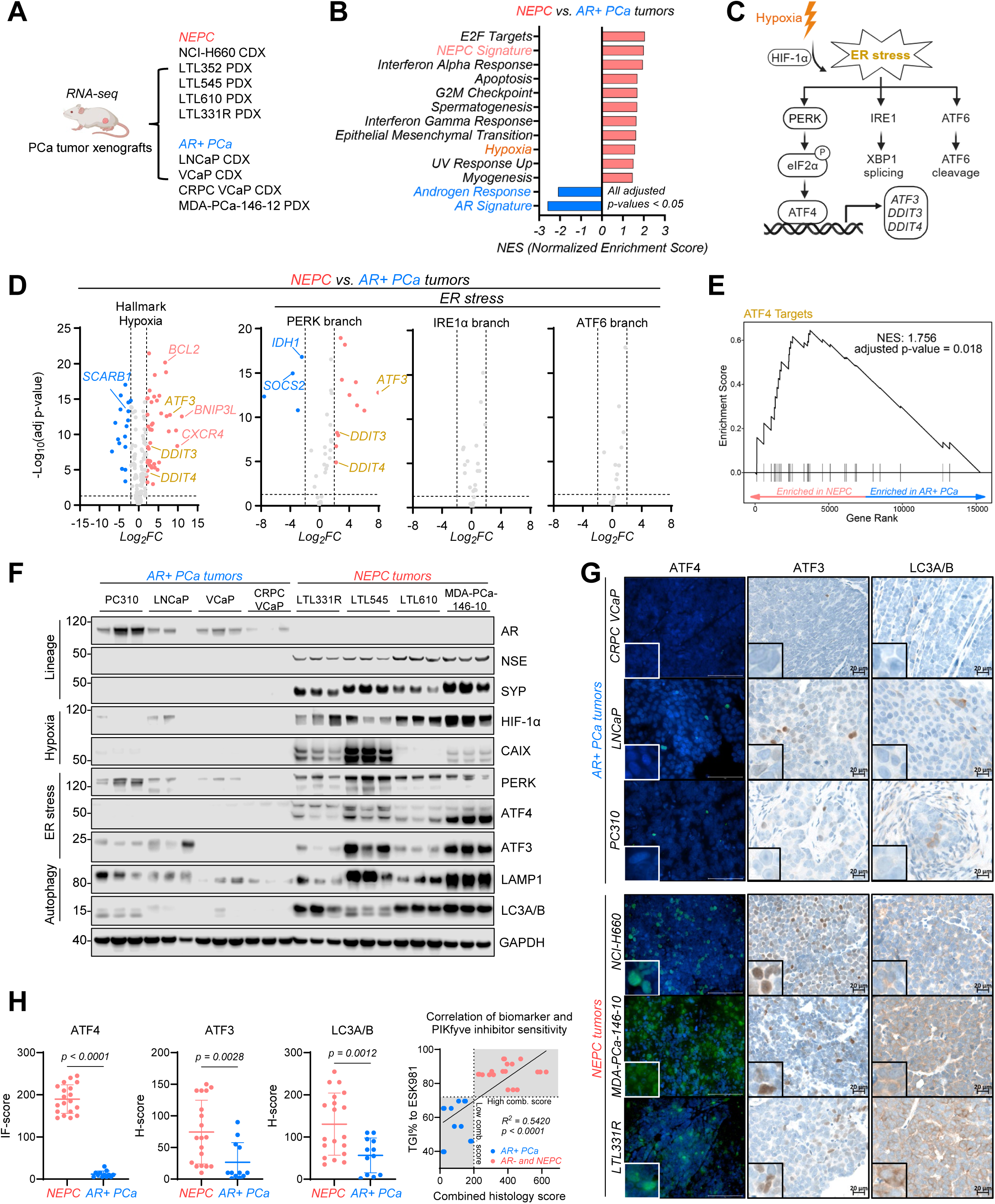
NEPC tumors exhibit elevated baseline ER stress and autophagy. A. Study design for RNA-seq sample collection of multiple AR+ prostate and NEPC tumors. Sample sizes for each tumor type: LNCaP (*n* = 3), VCaP (*n* = 5), CRPC VCaP (*n* = 4), MDA-PCa-146-12 (*n* = 2), NCI-H660 (*n* = 4), LTL352 (*n* = 4), LTL545 (*n* = 2), LTL610 (*n* = 8), and LTL331R (*n* = 4). B. Hallmark pathways significantly altered in NEPC tumors compared with AR+ prostate tumors. All pathways shown had adjusted *p*-values < 0.05. C. Diagram showing the activation of ER stress by hypoxia and three branches of ER stress. D. Differentially expressed genes (DEGs) involved in the hypoxia pathway and three branches of ER stress pathway in NEPC versus AR+ prostate tumors. DEGs were defined using thresholds of log_2_FC greater than 2 or less than -2 and adjusted *p-*value below 0.05. E. GSEA of ATF4 targets in NEPC tumors compared with AR+ prostate tumors. F. Western blot showing key proteins in lineage markers, ER stress, autophagy, and hypoxia pathways in NEPC (*n* = 3 for each tumor type) and AR+ prostate (*n* = 3 for each tumor type) tumors. G. Representative immunofluorescence (IF) images of ATF4 and IHC images of ATF3 and LC3A/B in NEPC and AR+ prostate tumors. H. IF-scores of ATF4 and H-scores of ATF3 and LC3A/B in NEPC (*n* = 19) and AR+ prostate (*n* = 12) tumors. Data presented as mean ± SD*. P* values calculated using a two-tailed unpaired *t*-test with Welch’s correction (left). Correlation analysis between combined histology score of individual tumors and tumor growth inhibition (TGI %) following ESK981 treatment (right). The combined histology score was calculated by summing ATF4 IF-scores and ATF3 and LC3A/B H-scores with equal weighting for each marker. Each data point represents an individual tumor across models. Tumors were stratified into high-score and low-score groups using a combined histology score threshold of 200 and an ESK981 TGI % threshold of 72%. Pink indicates high combined histology score/greater ESK981 sensitivity, and blue indicates low combined histology score/lower ESK981 sensitivity.

Hypoxia is known to induce ER stress^6^ by promoting the accumulation of misfolded proteins (**Figure 3C**). We, therefore, examined the activation of the UPR pathways in NEPC tumors. Transcriptomic analysis revealed predominant upregulation of the PERK-ATF4 branch of the UPR, including increased expression of downstream effectors such as *ATF3*, *DDIT3*, and *DDIT4*, whereas no changes were observed in markers of the IRE1-XBP1 and ATF6 branches of the UPR (**Figures 3D, S4F, and S4G**). GSEA confirmed enrichment of ATF4 downstream targets^33^ in NEPC tumors (**Figures 3E**), and the increased *ATF3* mRNA expression was validated by qPCR (**Figure S4H**). Notably, ATF4 protein levels were markedly increased in NEPC tumors (**Figure 3F**), despite no significant change in *ATF4* mRNA expression (**Figure S4H**). This suggests translational regulation of ATF4 occurs in NEPC tumors, consistent with previous findings^34^.

Autophagy is one of the key adaptive mechanisms to resolve ER stress^35^. Protein markers of autophagy, including LC3A/B and LAMP1, were enriched in NEPC tumors along with hypoxia-(HIF-1α, CAIX) and ER stress-associated proteins (ATF4, ATF3), while levels of ATF6 and XBP1s were not consistently enriched in one tumor type compared to the other (**Figures 3F, S4I, and S4J**). These findings were further supported by increased nuclear staining of ATF4 by immunofluorescence (IF) and IHC analysis showing increased ATF3 and LC3A/B expression in NEPC relative to AR+ prostate tumors (**Figures 3G and 3H**), along with upregulation of *LC3B* mRNA expression (**Figure S4K**). The combined histology score of ATF4, ATF3, and LC3A/B positively correlated with the *in vivo* tumor growth inhibition percentage to PIKfyve inhibitor ESK981 (R² = 0.5420, *p* < 0.0001; **Figure 3H**), suggesting that tumors with higher baseline stress pathway activation may be more dependent on PIKfyve-mediated adaptive mechanisms and exhibit enhanced sensitivity to PIKfyve inhibition. These findings support the potential use of baseline stress and autophagy markers to inform biomarker-driven prioritization of tumors for ESK981-based treatment strategies. Together, these data demonstrate that NEPC tumors exhibit a heightened stress-adaptive state characterized by hypoxia-associated ER stress, predominant activation of the PERK-ATF4 pathway, and concomitant upregulation of autophagy.

### ER stress increases NEPC dependency on PIKfyve

To assess the role of PIKfyve in the context of ER stress, we modeled *in vitro* ER stress conditions pharmacologically using thapsigargin or physiologically through serum starvation. Thapsigargin, an ER calcium pump inhibitor^36^, robustly increased ATF4 and ATF3 protein levels, along with upregulation of *ATF3* and *DDIT3* mRNA expression in LTL331R-CL and NCI-H660 cells (**Figures S5A and S5B**), confirming effective ER stress induction *in vitro*. To probe the role of PIKfyve in NEPC, we employed a potent PROTAC degrader of PIKfyve, PIK5-33d (**Figure S5C**), developed based on our previous work^37^. Quantitative proteomics analysis confirmed PIKfyve was the only depleted protein by PIK5-33d in LTL331R-CL cells after four hours of treatment, confirming the specificity of PIK5-33d to PIKfyve (**Figure 4A**). Immunoblotting demonstrated dose- and time-dependent PIKfyve degradation with accumulation of lipidated LC3A/B after PIK5-33d treatment in LTL331R-CL cells, consistent with disrupted autophagic flux, which was similarly observed in NCI-H660 cells (**Figures S5D and S5E**). Drug repurposing analysis using the Profiling Relative Inhibition Simultaneously in Mixtures (PRISM) screening platform identified APY0201, apilimod, and YM-201636, all established PIKfyve inhibitors, as top correlates of PIK5-33d sensitivity, further supporting PIK5-33d as a potent and specific tool compound for targeting PIKfyve (**Figure 4B**).

**Figure 4.**
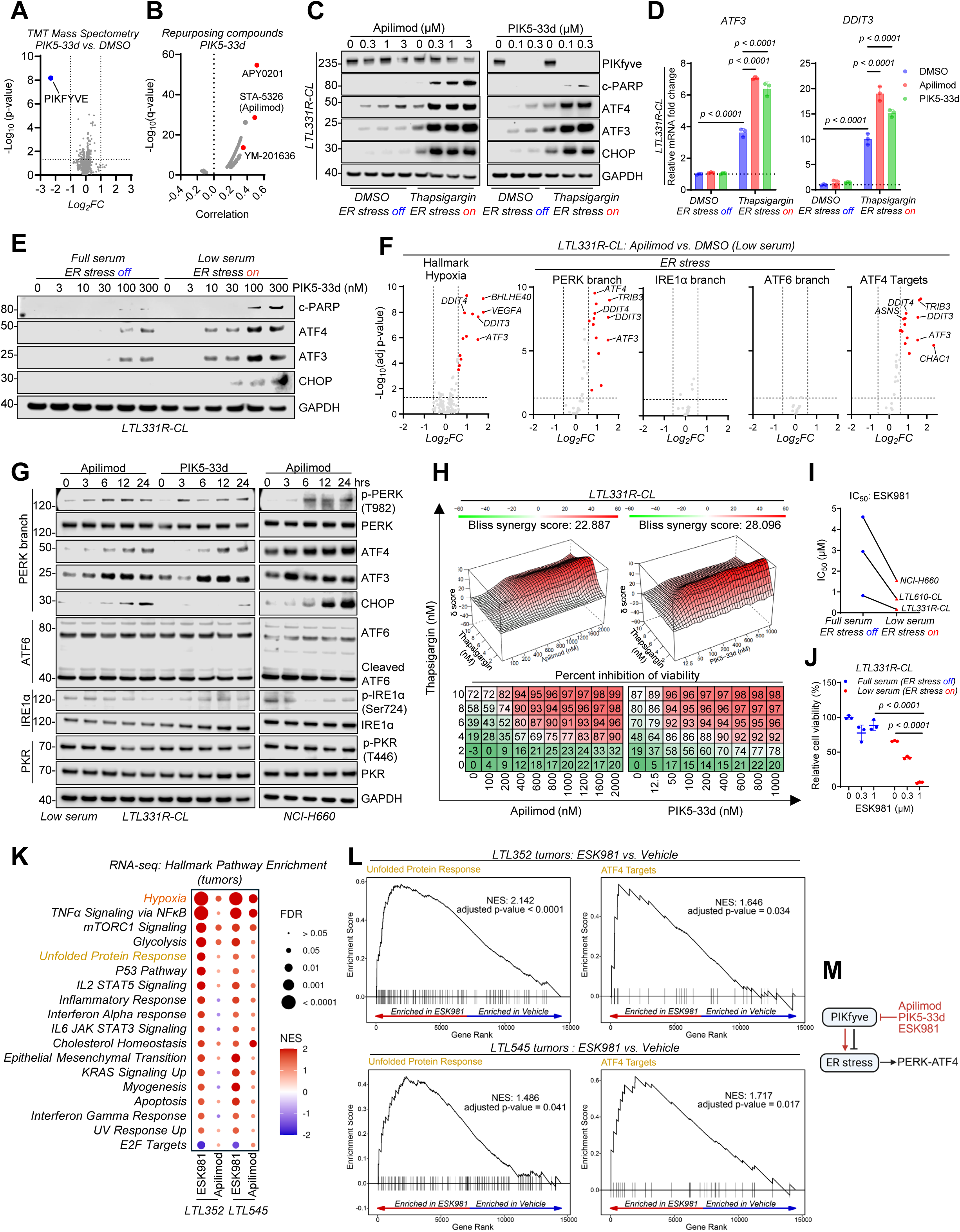
ER stress sensitizes NEPC cells to PIKfyve inhibition. A. TMT mass spectrometry proteomics analysis in LTL331R-CL cells treated with either DMSO or PIK5-33d (0.1 µM) for 4 hours. Differentially expressed proteins (DEPs) were defined using thresholds of log_2_FC greater than 1 or less than -1 and adjusted *p-*value below 0.05. B. Correlation analysis of repurposing compounds and PIK5-33d sensitivity determined by high-throughput PRISM screen. Red highlighted compounds are known PIKfyve inhibitors. C. Western blot of PIKfyve, c-PARP, ATF4, ATF3, and CHOP in LTL331R-CL treated with various concentrations of apilimod or PIK5-33d with or without thapsigargin (10 nM) treatment for 24 hours. D. qPCR of *ATF3* and *DDIT3* mRNA expression in LTL331R-CL cells treated with DMSO, apilimod (1 µM), or PIK5-33d (0.1 µM), with or without thapsigargin (10 nM) treatment for 6 hours. Data presented as mean ± SD*. P* values calculated using two-way ANOVA. E. Western blot of c-PARP, ATF4, ATF3, and CHOP in LTL331R-CL cells treated with increasing concentrations of PIK5-33d under full serum or low serum conditions for 24 hours. F. DEGs associated with hypoxia and the three branches of the ER stress pathway (PERK, ATF6, and IRE1α), as well as ATF4 target genes, in LTL331R-CL cells treated with apilimod (1 µM) versus DMSO under low-serum conditions for 6 hours. DEGs were defined using thresholds of log_2_FC greater than 0.58 or less than -0.58 and adjusted *p-*value below 0.05. G. Western blot analysis of key proteins involved in the three branches of ER stress and PKR of the integrated stress response pathway following treatment with apilimod (1 µM) or PIK5-33d (0.3 µM) for the indicated time points in LTL331R-CL and NCI-H660 cells. H. Synergy matrix and percent inhibition of viability between thapsigargin and apilimod or PIK5-33d treatment for 7 days in LTL331R-CL cells. Synergy scores were calculated using the Bliss independence model, with scores above 10 indicating synergy. I. IC_50_ values of ESK981 in NEPC cells under full serum or low serum conditions for 7 days. J. Relative cell viability of LTL331R-CL cells treated with increasing concentrations of ESK981 under full serum or low serum conditions for 7 days. Data presented as mean ± SD*. P* values calculated using two-way ANOVA. K. Hallmark pathway enrichment analysis post ESK981 or apilimod treatment for 5 days, compared to vehicle, in LTL352 and LTL545 NEPC PDXs *in vivo*. L. GSEA of unfolded protein response and ATF4 targets in LTL352 and LTL545 NEPC tumors after treatment of ESK981 and vehicle for 5 days. M. Model schematic illustrating that PIKfyve restrains ER stress and that PIKfyve inhibition (apilimod, PIK5-33d, ESK981) amplifies ER stress.

Importantly, combining PIK5-33d or apilimod with thapsigargin markedly amplified ATF4-ATF3-CHOP signaling at both mRNA and protein levels, leading to increased c-PARP expression in LTL331R-CL and NCI-H660 cells (**Figures 4C, 4D, and S5F**). Similarly, serum starvation as a physiological ER stressor led to ATF4 and ATF3 upregulation (**Figure S5G**) and sensitized NEPC cells to PIKfyve inhibition, as indicated by enhanced c-PARP induction and activation of the ATF4-ATF3-CHOP axis in LTL331R-CL and NCI-H660 cells (**Figures 4E and S5H-J**). Given that CHOP (encoded by *DDIT3)* functions as a central pro-apoptotic effector during terminal UPR activation^15^, these findings suggest that NEPC cells rely on PIKfyve to restrain CHOP-mediated apoptosis under ER stress. In melanoma models, PIKfyve inhibition has been shown to induce IL24 expression through CHOP activation^38^. Consistent with this, treatment of A375 melanoma cells with various PIKfyve inhibitors (WX8, ESK981, apilimod, and PIK5-33d) robustly induced *IL24* mRNA expression (**Figure S5K**). However, in LTL331R-CL NEPC cells, *IL24* was only marginally induced or unchanged under the same treatments (**Figure S5L**), suggesting a divergent apoptotic program in NEPC compared to melanoma, whereby CHOP activation promotes apoptosis independently of IL24. These collective results demonstrate that ER stress amplifies NEPC cell dependency on PIKfyve to suppress CHOP-mediated apoptosis. Disruption of PIKfyve function under ER stress leads to terminal UPR and apoptotic cell death, underscoring a context-dependent vulnerability.

Proteasome inhibition can induce ER stress^39^. To determine whether ER stress induced by PIKfyve inhibition results from lysosomal dysfunction or proteasome impairment, we compared PIKfyve inhibitors (ESK981, apilimod, PIK5-33d) with lysosomal inhibitors (bafilomycin A1, chloroquine). PIKfyve inhibition phenocopied lysosomal blockade by inducing ATF4 and LC3 lipidation (**Figure S5M**) but did not impair proteasome activity or increase ubiquitinated proteins, in contrast to proteasome inhibitor bortezomib (**Figures S5N and S5O**). These results indicate that PIKfyve inhibition-induced ER stress is driven by lysosomal dysfunction rather than proteasome inhibition.

To assess which ER stress pathways are engaged by PIKfyve inhibition, we analyzed RNA-seq data comparing apilimod, ESK981, and PIK5-33d with DMSO in LTL331R-CL cells. PIKfyve inhibition predominantly enriched the PERK pathway, particularly ATF4 target genes, without clear activation of the IRE1α or ATF6 branches (**Figures 4F and S6A; Tables S2-S5**). Time-course qPCR confirmed rapid PERK activation, with only modest trends toward IRE1α signaling at late time points (**Figure S6B**). In contrast, tunicamycin, which blocks N-linked glycosylation in the ER, and thapsigargin, which disrupts ER calcium homeostasis, induced all three UPR branches, albeit with distinct kinetics (**Figure S6C**). Consistently, immunoblotting showed increased phospho-PERK following PIKfyve inhibition, whereas ATF6 cleavage, phospho-IRE1α, phospho-PKR, and *XBP-1* mRNA splicing remained largely unchanged (**Figures 4G and S6D**). By comparison, tunicamycin increased PERK phosphorylation, ATF4/CHOP, cleaved ATF6, and XBP-1s levels (**Figures S6E and S6F**). These data indicate that PIKfyve inhibition predominantly activates the PERK-ATF4 branch of ER stress in NEPC cells.

PERK can sense lipid bilayer perturbation independently of proteotoxic stress^40,41^. We therefore asked whether PIKfyve inhibition altered membrane lipid properties in NEPC cells using the c-Laurdan generalized polarization (GP) assay. Apilimod reduced whole-cell GP in LTL331R-CL cells, similar to the cholesterol-depletion control MβCD, indicating decreased membrane lipid packing and increased membrane fluidity (**Figure S7A**). Although this assay does not resolve ER-specific membrane changes, these data suggest that PIKfyve inhibition induces broader membrane lipid bilayer perturbation that may contribute to PERK activation.

Because PIKfyve-complex disruption has been reported to acutely modulate lysosome-ER-mitochondria crosstalk and mitochondrial respiration^42^, and mitochondrial stress can activate the HRI-eIF2α pathway^43^, we next tested whether mitochondrial dysfunction contributes to integrated stress response (ISR) activation in NEPC cells. Under the same low-serum conditions in which PIKfyve inhibition induced ER stress markers, apilimod or PIK5-33d did not alter mitochondrial membrane potential, induce mitochondrial superoxide, or increase whole-cell reactive oxygen species (ROS), despite robust responses to positive controls in each assay (**Figures S7B-S7D**). These findings are consistent with the recent work showing that PIKfyve inhibition does not reduce basal oxygen consumption in pancreatic cancer cells^23^. Moreover, *EIF2AK1*/*HRI* knockdown had little effect on PIKfyve inhibition-mediated eIF2α phosphorylation or PARP cleavage (**Figure S7E**). Together, these data argue against detectable mitochondrial stress or HRI as primary mediators of ISR activation under these conditions and support a PERK-dominant mechanism in NEPC cells.

To further validate the functional consequence of PIKfyve inhibition under *in vitro* ER stress conditions, we evaluated the viability of NEPC cells under induced ER stress conditions in combination with PIKfyve inhibition. Co-treatment with PIKfyve inhibitors and thapsigargin or serum starvation synergistically decreased viability in NCI-H660, LTL331R-CL, and LTL610-CL cells (**Figures 4H-J and S8A-D**), indicating heightened dependency for cell proliferation on PIKfyve under ER stress. To test whether PIKfyve inhibition also modulates upstream ER stress signaling under hypoxia *in vivo*, we performed RNA-seq analysis of NEPC PDX tumors (LTL352 and LTL545) treated with either ESK981 or apilimod. Hallmark pathway enrichment analysis revealed upregulation of UPR and hypoxia pathways in both apilimod and ESK981 treated groups, with ESK981 showing more pronounced effects (**Figures 4K, S8E, and S8F**). GSEA further confirmed robust activation of UPR and ATF4 targets gene sets in ESK981-treated NEPC tumors (**Figure 4L**). These findings establish that elevated ER stress is a key driver of PIKfyve dependency in NEPC, and that PIKfyve disruption leads to UPR activation and apoptotic cell death when lysosomal resolution mechanisms are compromised (**Figure 4M**).

### PIKfyve restrains SREBP-dependent lipid biosynthesis

In addition to autophagy, cancer cells engage compensatory lipogenic approaches to restore membrane integrity and resolve ER stress^44,45^. PIKfyve has been implicated in lipid metabolism in PDAC^23^; however, its role in regulating lipid homeostasis in NEPC, particularly under ER stress, remains undefined. Given the intricate relationship between lysosomal function, ER stress, and lipid biosynthesis^46^, we hypothesized that PIKfyve may act as a critical regulator of lipogenic adaptation in NEPC. To test this hypothesis, we performed RNA-seq and proteomics profiling in LTL331R-CL cells treated with PIKfyve inhibitors (apilimod, PIK5-33d). Hallmark pathway enrichment analysis revealed that cholesterol and fatty acid metabolism were upregulated by both apilimod and PIK5-33d together with UPR and hypoxia (**Figure 5A**). GSEA on cholesterol homeostasis and fatty acid metabolism confirmed that these gene signatures were enriched after PIK5-33d, apilimod, or ESK981 treatment (**Figures 5B, S9A and S9B**). Analysis of differentially expressed genes showed that the top upregulated genes were involved in cholesterol homeostasis (*HMGCR*, *SQLE*, *FDFT1*, *HMGCS1*, *MVK*) and fatty acid metabolism (*SCD*, *ACSS2*, *ACLY*, *FASN*, *ACACA*) (**Figures 5C, 5D, and S9C**). These transcriptional changes were validated by qPCR in both LTL331R-CL and NCI-H660 cells (**Figure S9D**). The upregulation of fatty acid and cholesterol metabolism was also observed at the protein level, as evidenced by proteomics from whole cell lysates (**Figures 5E and S9E**) and confirmed by western blot with time- and dose-dependent induction of SCD1, FASN, ACSS2, and FDFT1 by apilimod and PIK5-33d in LTL331R-CL and NCI-H660 cells (**Figures 5F, S9F, and S9G**).

**Figure 5.**
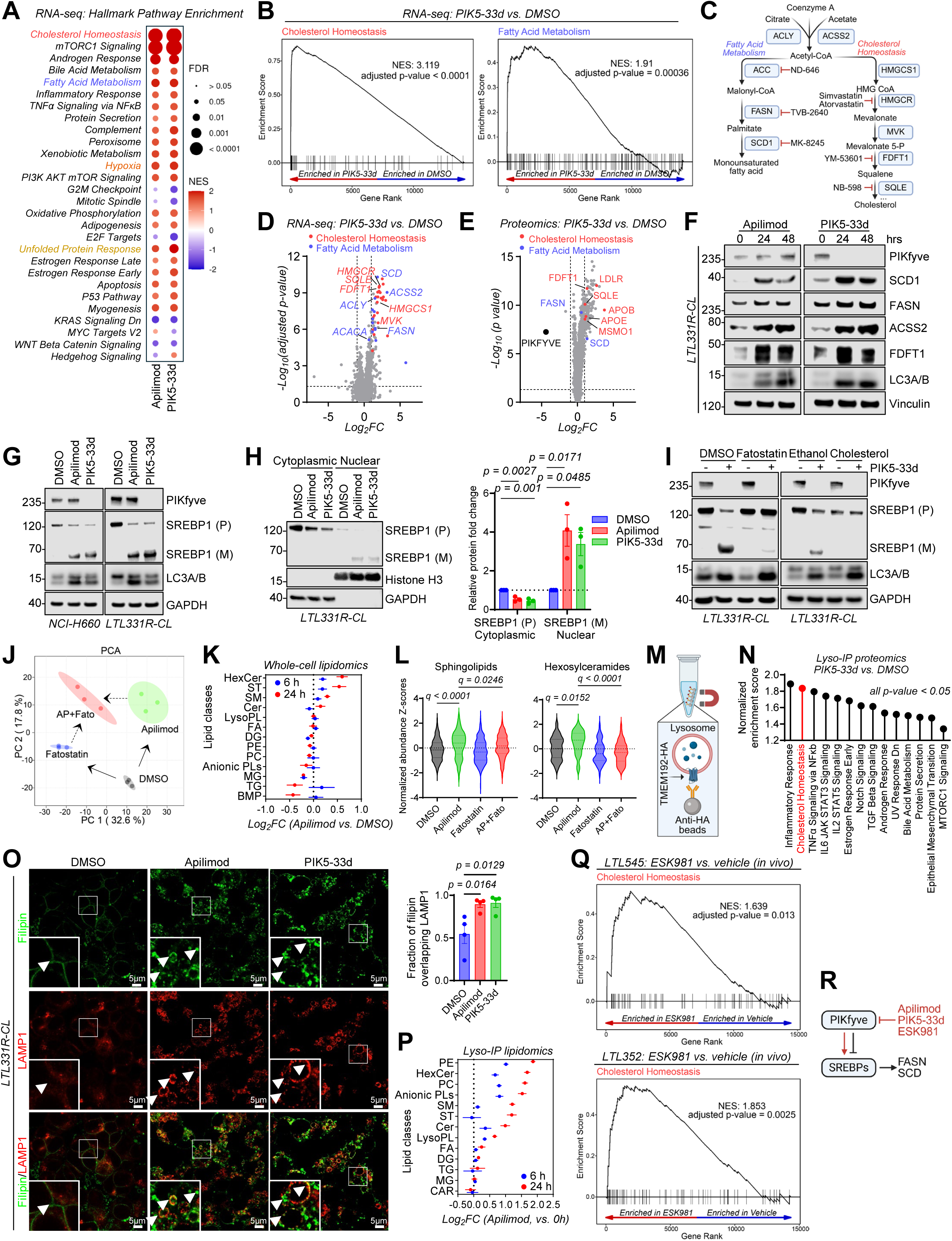
PIKfyve inhibition activates SREBP-dependent lipid biosynthesis. A. RNA-seq analysis showing significant hallmark pathways altered in LTL331R-CL cells post 6 hours of apilimod (1 µM) or PIK5-33d (0.3 µM) treatment compared with DMSO. B. GSEA plots showing enrichment of cholesterol homeostasis and fatty acid metabolism pathways in LTL331R-CL cells post treatment of PIK5-33d compared with DMSO. C. Pathway illustrating the major enzymes and products involved in fatty acid biosynthesis and cholesterol biosynthesis. D. Volcano plot of DEGs post 6 hours of PIK5-33d treatment, compared with DMSO, in LTL331R-CL cells. Genes upregulated in the cholesterol biosynthesis pathway are in red and fatty acid biosynthesis in cyan. DEGs were defined using thresholds of log_2_FC greater than 1 or less than -1 and adjusted *p-*value below 0.05. E. Volcano plot of DEPs post 24 hours of PIK5-33d treatment, compared with DMSO, in LTL331R-CL cells. Proteins increased in the cholesterol biosynthesis pathway are in red and fatty acid biosynthesis in cyan. DEPs were defined using thresholds of log_2_FC greater than 1 or less than -1 and adjusted *p-*value below 0.05. F. Western blot of the indicated targets involved in fatty acid and cholesterol biosynthesis in LTL331R-CL and NCI-H660 NEPC cells after apilimod (1 µM) or PIK5-33d (0.3 µM) treatment at the indicated time points. G. Western blot of PIKfyve, SREBP1 and LC3A/B in LTL331R-CL and NCI-H660 NEPC cells after DMSO, apilimod (1 µM), or PIK5-33d (0.3 µM) treatment for 6 hours. P: precursor; M: mature. H. Western blot analysis of cytoplasmic SREBP1 (P) and nuclear SREBP1 (M) in LTL331R-CL cells treated with DMSO, apilimod (1 µM), or PIK5-33d (0.3 µM) for 6 hours. GAPDH and histone H3 were used as loading controls for cytoplasmic and nuclear fractions, respectively (left). Quantification of relative protein levels of cytoplasmic SREBP1 (P) and nuclear SREBP1 (M) (right). Data presented as mean ± SEM. *P* values calculated using one-way ANOVA. I. Western blot of PIKfyve, SREBP1, and LC3A/B in LTL331R-CL cells after PIK5-33d (0.3 µM) treatment with or without fatostatin (20 µM) or cholesterol supplementation for 6 hours. J. Principal component analysis (PCA) of whole-cell lipidomic profiles from NCI-H660 cells treated with DMSO, apilimod (1 µM), fatostatin (20 µM), and the combination for 24 hours. K. Forest plot of lipid class changes of whole-cell lipidomic profiles from NCI-H660 cells post apilimod treatment compared with DMSO at indicated time points. L. Violin plots showing relative abundance (Z score-normalized) of sphingolipids and hexosylceramides of whole-cell lipidomic profiles from NCI-H660 cells across the indicated treatment conditions at 24 hours. *P* values calculated using one-way ANOVA. M. Schematic of the lysosome immunoprecipitation (lyso-IP) workflow using NEPC TMEM192-HA cells and anti-HA magnetic beads. N. Hallmark pathway enrichment analysis in TMEM192-HA tagged LTL331R-CL cells following 24 hours of PIK5-33d (0.1 µM) treatment compared to DMSO by lyso-IP based proteomics. O. Representative confocal images of filipin and LAMP1 co-staining post 24 hours of DMSO, apilimod (1 µM), or PIK5-33d (0.1 µM) treatment in LTL331R-CL cells. Filipin was shown in green pseudocolor, LAMP1 in red, and co-localization in yellow. Arrows indicated co-localization between filipin and LAMP1 (left). Quantification of filipin signal colocalizing with LAMP1 across treatment groups (right). *n* = 4 random fields. Experiments were repeated three times. Data presented as mean ± SEM*. P* values calculated using one-way ANOVA. P. Forest plot of lipid class changes of lysosomal lipidomic profiles from TMEM192-HA tagged NCI-H660 cells post apilimod treatment for 6 and 24 hours compared with 0 hour. Q. GSEA of cholesterol homeostasis in LTL352 and LTL545 NEPC tumors after treatment with ESK981 and vehicle for 5 days. R. Model schematic illustrating that PIKfyve restrains SREBP-mediated lipogenesis and that PIKfyve inhibition activates SREBPs.

SREBPs are master regulators of cholesterol and fatty acid biosynthesis, and when sterol levels are low, SREBPs initiate proteolytic cleavage to generate mature SREBPs as transcription factors for fatty acid and cholesterol biogenesis^47–49^. We sought to examine whether PIKfyve inhibition-induced cholesterol and fatty acid biosynthesis were SREBP-dependent. Treatment with PIK5-33d or apilimod led to marked accumulation of mature SREBP1 and SREBP2 in a time-dependent manner (detectable as early as 2 hours post treatment) in LTL331R-CL, NCI-H660, and LTL610-CL cells (**Figures 5G, S10A, and S10B**). Quantitative analysis of subcellular fractionation revealed a significant increase in nuclear SREBP1 (M), accompanied by a corresponding reduction in the cytoplasmic precursor form SREBP1 (P), indicating enhanced SREBP processing and nuclear translocation rather than changes in total protein abundance (**Figures 5H, S10C, and S10D**). Fatostatin binds to SREBP’s escort protein SCAP, preventing SREBPs from trafficking to the Golgi for activation^50^. Treatment with fatostatin suppressed SREBP activation and downstream target gene and protein expression induced by PIKfyve inhibition (**Figure 5I, S10E, and S10F**), supporting that lipid biosynthesis driven by PIKfyve inhibition is SREBP-dependent. Since SREBP activation is classically driven by reduced cytosolic cholesterol levels, we found that exogenous cholesterol supplementation attenuated PIKfyve inhibition-mediated SREBP activation (**Figure 5I**).

Untargeted whole-cell lipidomics was next performed to characterize global lipidomic changes following PIKfyve inhibition and to identify lipid classes synthesized *de novo* in an SREBP-dependent manner using fatostatin. Principal component analysis (PCA) revealed distinct lipidomic profiles between apilimod and fatostatin treatments, with the combination treatment producing a separable lipid state (**Figure 5J**). Rather than inducing broad lipid accumulation, apilimod treatment resulted in time-dependent increases in specific lipid classes, including hexosylceramides (HexCer), sterol lipids (ST), sphingomyelins (SM), and ceramides (Cer), whereas phosphatidylethanolamine (PE) and phosphatidylcholine (PC) levels remained largely unchanged (**Figure 5K**). Notably, co-treatment with fatostatin selectively reduced apilimod-induced increases in HexCer and ST. Given that HexCer, SM, and Cer are sphingolipids, we further observed that the apilimod-induced accumulation of sphingolipid species was specifically attenuated by fatostatin (**Figures 5L and S10G**). Consistently, analysis of the top lipid species enriched by apilimod and reversed upon fatostatin treatment demonstrated predominant representation of sphingolipid classes (**Figure S10H**), supporting that PIKfyve inhibition-induced synthesis of sphingolipids is SREBP-dependent.

To determine the underlying mechanism, we leveraged lysosomal immunoprecipitation (lyso-IP) using TMEM192-HA expressing NEPC cells (**Figure 5M**). Proteomics profiling of the lysosomal fraction revealed a globally enrichment-skewed lysosomal proteomic profile, with an enrichment of the cholesterol homeostasis pathway and an accumulation of lipid metabolism-associated enzymes, including FASN, SQLE, and FDFT1, following PIK5-33d or apilimod treatment, along with autophagy markers LC3 and p62 (**Figures 5N, S11A, and S11B**), suggesting impaired lysosomal turnover of lipid-related proteins. Intracellular cholesterol levels and localization were then assessed following PIKfyve inhibition. Both apilimod and PIK5-33d increased total cholesterol in NEPC cells, an effect largely suppressed by the HMGCR inhibitor simvastatin (**Figure S11C**). Intracellular cholesterol distribution was assessed by filipin staining to visualize unesterified cholesterol in combination with LAMP1 staining to label lysosomes in LTL331R-CL and NCI-H660 cells. PIKfyve inhibition by apilimod or PIK5-33d led to an increase in filipin fluorescence that colocalized with LAMP1-positive vacuoles, displaying both punctate and ring-like patterns suggestive of lysosomal cholesterol accumulation, whereas U18666A, an NPC1/2 inhibitor that blocks cholesterol efflux from lysosomes, produced punctate colocalization of filipin and LAMP1 without vacuolization in LTL331R-CL and NCI-H660 cells (**Figures 5O, S11D, and S11E**). These findings demonstrate that PIKfyve is required for lysosomal cholesterol efflux, and PIKfyve inhibition results in cholesterol accumulation within lysosomes and depletion of cytosolic cholesterol. Untargeted lipidomics of lysosome-enriched fractions further demonstrated a pronounced, time-dependent accumulation of lipid species following apilimod treatment (**Figures 5P and S11F**). Multiple lipid classes, including PE, PC, HexCer, and ST, were broadly enriched within lysosomes (**Figures 5P and S11G**), indicating generalized lipid retention associated with lysosomal dysfunction. Together, these findings suggest that PIKfyve inhibition disrupts lysosomal lipid handling and alters the intracellular availability of lipid species, giving rise to a selective lipid biosynthetic response rather than global lipid overproduction.

Importantly, pathway enrichment analysis of NEPC tumors treated *in vivo* with ESK981 or apilimod revealed consistent activation of cholesterol homeostasis pathways (**Figure 5Q**), indicating that the lipogenic adaptation observed *in vitro* is preserved *in vivo*. Moreover, SREBP activation was also observed in AR+ (LNCaP and VCaP) and AR-negative (DU145 and PC3) prostate cancer cells, suggesting that SREBP activation represents a conserved response to PIKfyve inhibition (**Figures S11H-J**).

Taken together, these data uncover a previously unrecognized function of PIKfyve in restraining SREBP activation in NEPC. By maintaining lysosomal cholesterol efflux and ER membrane composition, PIKfyve suppresses lipogenesis. Its inhibition leads to sterol depletion and SREBP-mediated lipid biosynthesis (**Figure 5R**).

### Dual inhibition of PIKfyve and lipogenesis induces a synthetic vulnerability

We next evaluated whether lipogenesis functions as a compensatory adaptation to PIKfyve inhibition, and whether targeting both pathways would produce a synthetic vulnerability in NEPC. To pharmacologically inhibit lipid biosynthesis, we used the FASN inhibitor TVB-2640 and the ACC inhibitor ND-646, both of which showed effective target engagement (**Figures S12A and S12B**). Co-treatment with either TVB-2640 or ND-646 with PIKfyve inhibitors (apilimod, PIK5-33d, or ESK981) produced strong synergistic anti-proliferative effects in LTL331R-CL cells (**Figures 6A, S12C, and S12D**). Consistent with the effects observed with SREBP inhibition, co-treatment with TVB-2640 markedly altered the lipidomic response to apilimod (**Figure 6B**), selectively suppressing apilimod-induced enrichment of HexCer (**Figures 6C, S12E, and S12F**). In addition, TVB-2640 further depleted TGs and fatty acids (FA), in line with effective FASN inhibition (**Figures 6C and S12G**). Together, these data indicate that PIKfyve inhibition induces a compensatory, SREBP-dependent lipogenic program that can be therapeutically exploited to induce a synthetic vulnerability in NEPC cells.

**Figure 6.**
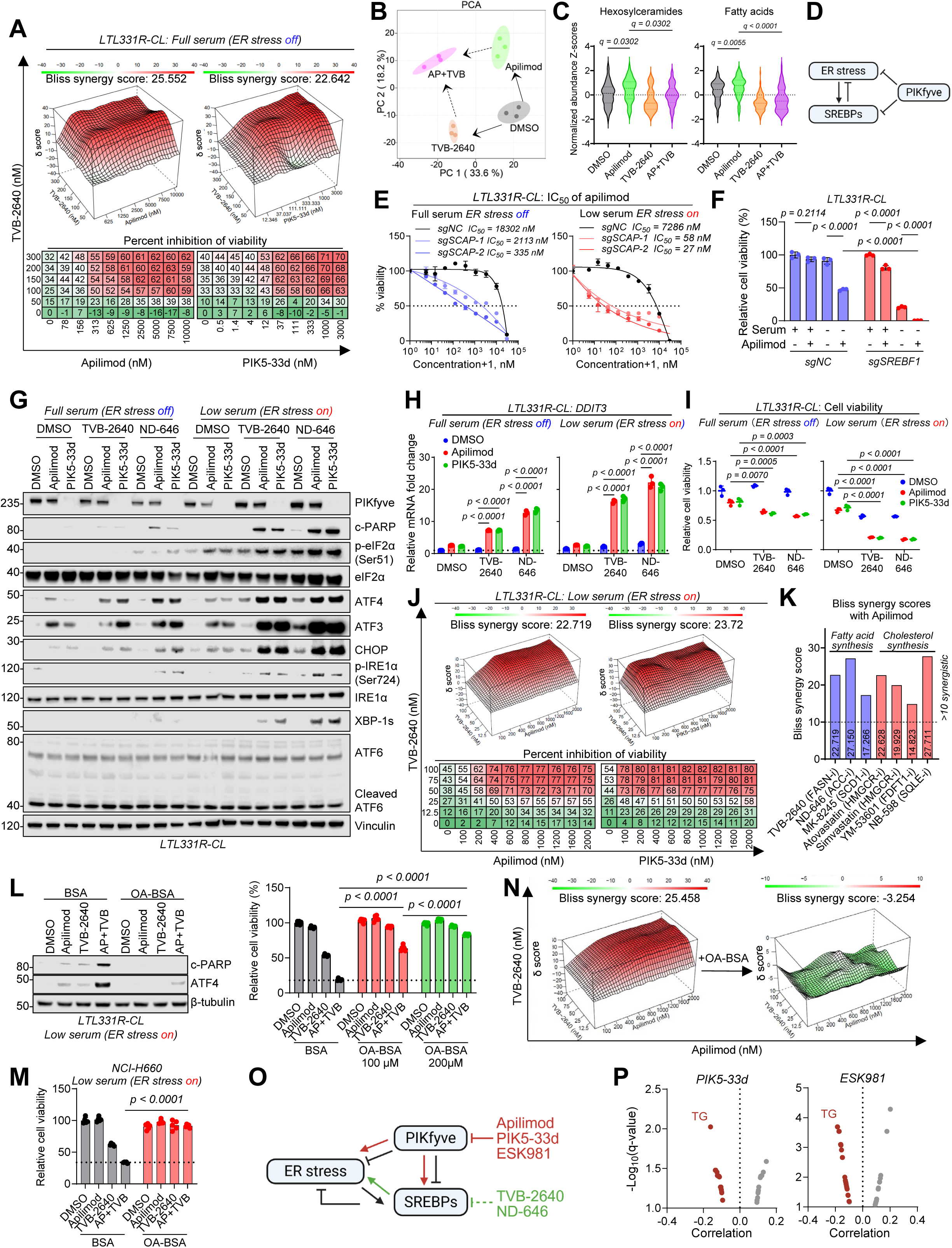
Dual inhibition of PIKfyve and lipogenesis induces synthetic vulnerability in NEPC. A. Synergy matrix and percent inhibition of viability of TVB-2640 and PIKfyve inhibitors (Apilimod or PIK5-33d) under full serum conditions for 7 days in LTL331R-CL cells. B. PCA of whole-cell lipidomic profiles from NCI-H660 cells treated with DMSO, apilimod (1 µM), TVB-2640 (1 µM), and the combination for 24 hours. C. Violin plots showing relative abundance (Z score-normalized) of hexosylceramides and fatty acids of whole-cell lipidomic profiles from NCI-H660 cells across the indicated treatment conditions at 24 hours. *Q* values calculated using Welch’s ANOVA, followed by pairwise multiple comparisons with Benjamini-Hochberg FDR correction. D. Model schematic depicting reciprocal regulation between ER stress, SREBPs, and PIKfyve: PIKfyve restrains ER stress and SREBPs activation, whereas ER stress promotes SREBPs activation, and SREBPs activation feeds back to attenuate ER stress. E. Dose-response analysis of apilimod sensitivity in LTL331R-CL cells expressing CRISPRi *sgNC*, *sgSCAP-1*, or *sgSCAP-2* under full serum and low serum conditions for 7 days. Data presented as mean ± SD. F. Relative cell viability of LTL331R-CL cells expressing CRISPRi *sgNC* or *sgSREBF1* under full serum or low serum conditions, with or without apilimod treatment (1 µM). Data presented as mean ± SD. *P* values calculated using two-way ANOVA. G. Western blot of PIKfyve, c-PARP and representative proteins from the three ER stress branches following DMSO, apilimod (1 µM), and PIK5-33d (0.3 µM) treatment, with or without TVB-2640 (1 µM) or ND-646 (1 µM), under full serum or low serum conditions in LTL331R-CL cells for 24 hours. H. qPCR of *DDIT3* mRNA expression after DMSO, apilimod (1 µM), and PIK5-33d (0.3 µM) treatment, with or without TVB-2640 (1 µM) or ND-646 (1 µM), under full serum or low serum conditions in LTL331R-CL cells for 6 hours. Data presented as mean ± SD*. P* values calculated using two-way ANOVA. I. Relative cell viability of LTL331R-CL cells treated with DMSO, apilimod (1 µM), and PIK5-33d (0.3 µM), with or without TVB-2640 (50 nM) or ND-646 (5 nM) under full serum and low serum conditions for 7 days. Data presented as mean ± SD*. P* values calculated using two-way ANOVA. J. Synergy matrix and percent inhibition of viability of TVB-2640 and PIKfyve inhibitors (apilimod or PIK5-33d) under low serum conditions for 7 days in LTL331R-CL cells. K. Summary of Bliss synergy scores between apilimod and multiple lipogenesis inhibitors under low serum conditions for 7 days in LTL331R-CL cells. L. Western blot analysis of c-PARP and ATF4 following treatment with DMSO, apilimod (1 µM), TVB-2640 (1 µM), or the combination treatment, supplemented with BSA or oleic acid-BSA (OA-BSA) (200 µM) under low serum conditions in LTL331R-CL cells for 24 hours (left). Relative cell viability of LTL331R-CL cells treated with DMSO, apilimod (1 µM), TVB-2640 (1 µM), or the combination in the presence of BSA or OA-BSA at the indicated concentrations for 5 days (right). Data presented as mean ± SD. *P* values calculated using two-way ANOVA. M. Relative cell viability of NCI-H660 cells treated with DMSO, apilimod (1 µM), TVB-2640 (1 µM), or the combination in the presence of BSA or OA-BSA (200 µM) under low serum conditions for 7 days. Data presented as mean ± SD. *P* values calculated using two-way ANOVA. N. Synergy matrix of TVB-2640 and apilimod, supplemented with BSA or OA-BSA (200 µM), under low serum conditions for 7 days in NCI-H660 cells. O. Model illustrating bidirectional crosstalk between PIKfyve, ER stress, and SREBP signaling: PIKfyve inhibition amplifies ER stress and promotes SREBP activation, while lipogenesis inhibition disrupts SREBP-mediated adaptive feedback. P. Correlation analysis of metabolites and PIK5-33d or ESK981 sensitivity determined by high-throughput PRISM screen. Triglycerides (TG) are shown in red.

ER stress is a known activator of SREBP-dependent lipogenesis^51^, while inhibition of lipid biosynthesis can reciprocally exacerbate ER stress^52^. Given that PIKfyve restrains both ER stress and lipogenesis, we asked whether elevated ER stress would further potentiate PIKfyve inhibition-induced lipogenic responses in NEPC (**Figure 6D**). Serum starvation, a physiological ER stress inducer, independently induced expression of lipogenesis-related pathways and genes such as *FASN* and *SCD* in LTL331R-CL cells (**Figure S13A**). Under these stress conditions, PIKfyve inhibition further amplified gene signatures associated with cholesterol and fatty acid biosynthesis, hypoxia, and the UPR (**Figure S13B**). qPCR validated synergistic upregulation of lipogenic genes (**Figure S13C**), and immunoblots confirmed enhanced activation of SREBP1, SCD1, and ATF4 (**Figure S13D**). To determine whether ATF4 contributes to PIKfyve inhibition-induced lipogenesis under ER stress, we generated CRISPRi-mediated *ATF4* knockdown NEPC cells (**Figure S13E**). *ATF4* depletion reduced NEPC cell proliferation (**Figure S13F**) and attenuated PIKfyve inhibition-induced activation of SREBP1 and downstream targets (*FASN* and *SCD1)*, without affecting the SREBP2 pathway (**Figures S13G-J**). These results support a model in which ER stress potentiates SREBP-mediated lipid biosynthesis in the context of PIKfyve inhibition.

CRISPRi-mediated *SCAP* knockdown sensitized NEPC cells to apilimod, causing a marked reduction in IC_50_ values under both basal and serum starvation-induced ER stress conditions (**Figures 6E and S13K**). Under full-serum conditions, CRISPRi-mediated *SCAP* knockdown reduced the apilimod IC_50_ by approximately tenfold, whereas under ER stress the IC_50_ was reduced by over 100-fold. These findings highlight a critical role for SCAP-dependent SREBP activation in mitigating PIKfyve inhibition-induced stress, especially under conditions of elevated basal ER stress. Similarly, depletion of *SREBF1* abrogated apilimod-induced *FASN* and *SCD1* expression, concomitantly enhanced ATF4-mediated ER stress, and resulted in near-complete loss of cell viability under ER stress (**Figures 6F and S13L**). Notably, *ATF4* knockdown phenocopied *SREBF1* knockdown for enhanced apilimod-induced apoptosis, as evidenced by elevated c-PARP and exacerbated loss of cell viability (**Figures S13M-O**), indicating that ATF4-dependent SREBP1 activation promotes cellular tolerance of PIKfyve inhibition under ER stress conditions. In summary, PIKfyve, ER stress, and lipogenesis form an interdependent regulatory network in NEPC (**Figure 6D**).

Consistent with previous results which showed that inhibition of lipid biosynthesis can activate autophagy^53–55^, both TVB-2640 and ND-646 increased autophagic flux, especially under serum starvation, as measured by GFP-LC3-RFP-LC3ΔG DU145 and PC3 reporter cell lines (**Figure S14A**), supporting a reciprocal relationship between lipid metabolism and autophagy under ER stress. We then combined PIKfyve inhibitors (apilimod or PIK5-33d) with lipogenesis inhibitors (TVB-2640 or ND-646) in LTL331R-CL cells. These combinations robustly activated the PERK-eIF2α-ATF4-CHOP pathway, as evidenced by elevated levels of *ATF3* and *DDIT3* transcripts and p-eIF2α, ATF4, ATF3, CHOP, and c-PARP protein levels, which was more prominent under ER stress conditions. In contrast, autophagy markers, including LC3A/B and p62, and lipogenic enzymes, including FASN and SCD1, were largely induced by PIKfyve inhibition alone and maintained following combination treatment (**Figures 6G, 6H, S14B, and S14C**). Pharmacologic inhibition of PERK, but not inhibition of HRI, ATF6α, or IRE1, attenuated ATF4 induction under combination treatment (**Figure S14D**). Consistently, *EIF2AK1/HRI* knockdown did not suppress PERK-ATF4 pathway activation (**Figure S14E**). Concomitantly, the IRE1α branch was engaged, supported by increased p-IRE1α relative to total IRE1α and induction of canonical downstream targets (*XBP-1s* and *EDEM1*) upon combination treatment (**Figures 6G and S14F**). By contrast, ATF6 cleavage was readily detectable under basal conditions but was not further enhanced by combination treatment, and expression of ATF6 target genes (*GRP94* and *ERP72*) remained unchanged (**Figures 6G and S14F**), indicating predominant engagement of the PERK and IRE1α pathways without additional involvement of the ATF6 branch^40,41,56,57^. These molecular responses translated into pronounced loss of cell viability, which was markedly enhanced under serum starvation conditions (**Figures 6I and S14G**) and phenocopied by thapsigargin-induced ER stress (**Figure S14H**).

Importantly, this synthetic vulnerability was not restricted to fatty acid synthesis blockade. Combined inhibition of PIKfyve and cholesterol biosynthesis using the HMGCR inhibitor, atorvastatin, similarly exacerbated ER stress and apoptosis under serum starvation (**Figure S15A**). Moreover, pharmacological disruption of lysosomal function with chloroquine or bafilomycin A1 further sensitized cells to fatty acid synthesis inhibition, leading to increased ER stress and apoptosis (**Figures S15B and S15C**). Together, these findings indicate that concurrent disruption of lysosomal function and lipid biosynthesis poses a convergent vulnerability in NEPC cells.

To determine the dominant mode of cell death induced by dual inhibition, we evaluated the effects of pathway-specific cell death inhibitors. PARP cleavage was completely abolished by the pan-caspase inhibitor Q-VD-OPh, while ER stress markers ATF3 and CHOP remained elevated, indicating that ER stress occurs upstream of caspase activation and that apoptosis is caspase dependent (**Figure S15D**). In contrast, markers of ferroptosis and necroptosis were not significantly altered (**Figure S15D**), and only caspase inhibition, rather than ferroptosis or necroptosis inhibitors, partially rescued the cell viability loss induced by the combination treatment (**Figure S15E**). Although *ATF4* knockdown sensitized cells to PIKfyve inhibition alone, it did not further enhance apoptosis or viability loss induced by dual inhibition (**Figures S15F and S15G**), supporting that ATF4 acts upstream to facilitate SREBP-1 dependent adaptive lipogenesis upon PIKfyve inhibition.

Synergy matrices revealed that serum starvation markedly enhanced drug cooperativity, reducing the effective concentrations required for synergy by approximately 3- to 10-fold across PIKfyve and lipogenesis inhibitors (**Figures 6J and S16A**). Similar synergy was independently validated in additional NEPC cell lines, including NCI-H660 and LTL610-CL cells (**Figures S16B and S16C**). To assess the breadth of this vulnerability, we extended our combination testing to inhibitors of other key lipogenesis related enzymes, including HMGCR (atorvastatin, simvastatin), FDFT1 (YM-53601), SQLE (NB-598), and SCD1 (MK-8245) in LTL331R-CL cells. All inhibitors exhibited synergistic effects with apilimod, indicating that PIKfyve inhibition renders NEPC broadly reliant on lipid biosynthetic pathways under ER stress (**Figures 6K, S16D, and S16E**). We next examined whether this synthetic vulnerability is further modulated by hypoxia, another physiologically relevant inducer of ER stress. Mild hypoxia was modeled using CoCl_2_ treatment, while severe hypoxia was induced using 0.1% O_2_, with normoxia (21% O_2_) as control. Hypoxic conditions stabilized HIF-1α and concomitantly induced ATF4 expression and LC3A/B lipidation (**Figure S16F**). Importantly, the Bliss synergy score between apilimod and TVB-2640 was highest in 0.1% O_2_ conditions (**Figure S16G**), supporting the notion that dual inhibition of PIKfyve and lipogenesis is more effective in microenvironments characterized by elevated ER stress.

To determine whether the lipids induced by PIKfyve inhibition functionally contribute to cell survival, we supplemented cells with oleic acid-BSA (OA-BSA) following combined apilimod and TVB-2640 treatment. OA-BSA supplementation rescued the PARP cleavage and induction of ATF4 induced by the combination treatment (**Figure 6L**). OA-BSA supplementation also rescued cell viability in a concentration-dependent manner in LTL331R-CL cells (**Figure 6L**). Similar results were observed in NCI-H660 cells, where OA-BSA completely rescued the loss of cell viability induced by the combination treatment and reverted the synergy between apilimod and TVB-2640 (**Figures 6M, 6N, and S16H**). Together, these data demonstrate that lipid availability is a critical determinant of cellular survival upon PIKfyve inhibition, with SREBP-dependent lipogenesis acting as a compensatory stress-buffering program rather than a passive metabolic byproduct, particularly in the context of elevated ER stress (**Figure 6O**).

To assess whether the stress-lipid dependency uncovered in NEPC reflects a broader metabolic principle underlying PIKfyve sensitivity, we leveraged the PRISM platform using PIK5-33d and ESK981 across almost 900 cell lines^58^. Correlation analysis revealed a striking inverse association between intracellular triglyceride (TG) abundance and PIK5-33d or ESK981 sensitivity (**Figure 6P**). TG accumulation is a well-established metabolic feature of hypoxic and ER-stressed states, including in both tumors (e.g., hepatoma cells) and non-tumor cell types (e.g., macrophages), as well as in hypoxic tissues such as the adrenal gland and liver serum^45,59–62^. While PIKfyve dependency in these contexts has not been systemically examined, the conserved association between TG abundance and PIKfyve inhibitor sensitivity observed here supports a broader model in which PIKfyve functions as a stress-adaptive metabolic node, creating a shared vulnerability in tumors characterized by elevated hypoxia and ER stress.

### Dual inhibition of PIKfyve and fatty acid synthesis confers a prolonged survival benefit

We next assessed the translational potential of this dual PIKfyve-lipogenesis targeting strategy *in vivo* using TVB-2640 and ESK981, both of which are under clinical investigation in CRPC as monotherapies (NCT05743621 and NCT05988918, respectively). Consistent with our *in vitro* findings, combination-treated LTL331R NEPC tumors showed elevated UPR and ATF4 transcriptional signatures at PD5 compared with vehicle- or ESK981-treated tumors (**Figures 7A and S17A**). Immunoblot analysis further showed c-PARP induction in ESK981-containing treatment groups, including the combination group, supporting apoptosis following PIKfyve inhibition-based treatment *in vivo* (**Figure 7B**). To capture early pharmacodynamic changes, we performed an additional PD3 study in the LTL331R NEPC PDX model. Compared with vehicle, the combination treatment showed pharmacodynamic pathway engagement, including increased eIF2α phosphorylation and HMGCR induction, while ESK981-induced autophagy markers, including p62 and LC3A/B, remained elevated in the combination group (**Figures 7C, 7D, and S17B**). Together, these *in vivo* data support the proposed mechanism of dual PIKfyve and FASN inhibition, linking sustained UPR activation with impaired stress adaptation and apoptosis in NEPC tumors.

**Figure 7.**
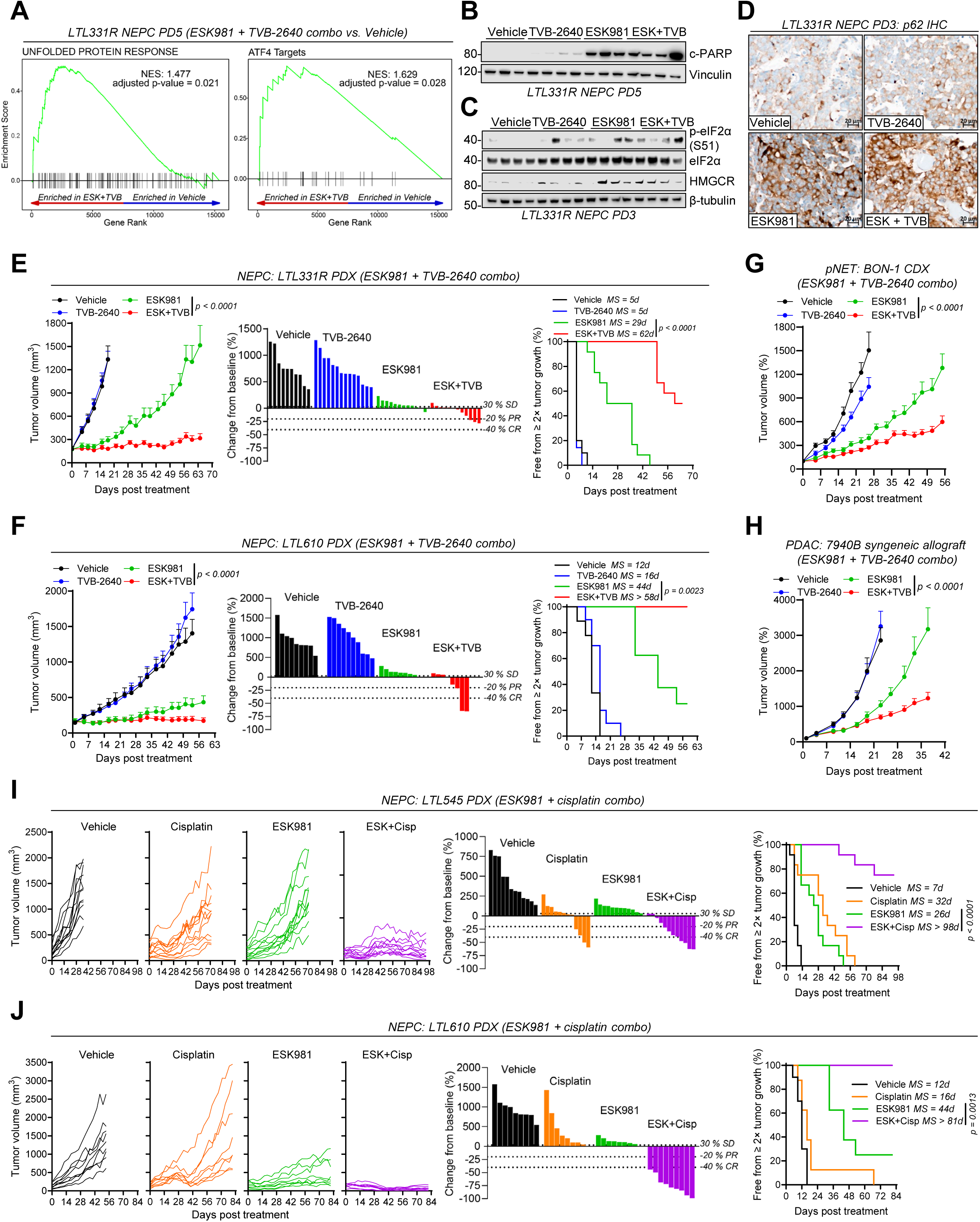
Translational *in vivo* evaluation of PIKfyve-targeted combination therapies. A. GSEA of hallmark unfolded protein response (left) and ATF4 target (right) gene set in LTL331R NEPC tumors treated with the combination of ESK981 and TVB-2640 compared with vehicle for 5 days. B. Western blot of c-PARP in LTL331R NEPC tumors treated with vehicle, TVB-2640, ESK981, or the combination of ESK981 and TVB-2640 for 5 days. C. Western blot of indicated targets in LTL331R NEPC tumors treated with vehicle, TVB-2640, ESK981, or the combination of ESK981 and TVB-2640 for 3 days. D. Representative IHC images of p62 in LTL331R NEPC tumors treated with vehicle, TVB-2640, ESK981, or the combination of ESK981 and TVB-2640 for 3 days. E-F. Tumor growth curves (left), waterfall plots of percent change from baseline (middle), and Kaplan-Meier analyses showing freedom from ≥2× tumor growth (right) in NEPC LTL331R (E) and LTL610 (F) PDX models treated with vehicle, TVB-2640, ESK981, or the combination. Sample sizes: Vehicle (*n* = 10 for LTL331R; *n* = 9 for LTL610), TVB-2640 (*n* = 14 for LTL331R; *n* = 10 for LTL610), ESK981 (*n* = 12 for LTL331R; *n* = 8 for LTL610), and ESK981 + TVB-2640 (*n* = 12 for LTL331R; *n* = 8 for LTL610). MS, median survival. G-H. Tumor growth curves in pNET BON-1 CDX (G) and PDAC 7940B syngeneic allograft (H) models treated with vehicle, TVB-2640, ESK981, or the combination. Sample sizes: Vehicle (*n* = 12 for 7940B; *n* = 16 for BON-1), TVB-2640 (*n* = 16 for 7940B; *n* = 17 for BON-1), ESK981 (*n* = 16 for 7940B and BON-1), and ESK981 + TVB-2640 (*n* = 16 for 7940B; *n* = 18 for BON-1). I-J. Individual tumor growth curves (left), waterfall plots of percent change from baseline (middle), and Kaplan-Meier analyses showing freedom from ≥2× tumor growth (right) in NEPC LTL545 (I) and LTL610 (J) PDX models treated with vehicle, cisplatin, ESK981, or the combination. Sample sizes: Vehicle (*n* = 12 for LTL545; *n* = 10 for LTL610), cisplatin (*n* = 12 for LTL545; *n* = 8 for LTL610), ESK981 (*n* = 12 for LTL545; *n* = 8 for LTL610), and ESK981 + cisplatin (*n* = 12 for LTL545; *n* = 9 for LTL610). The Vehicle and ESK981 groups in Fig. 7F and Fig. 7J were shared from the same LTL610 animal study; different combination arms are displayed in the two panels to highlight distinct comparisons. Data presented as mean ± SEM where applicable. *P* values calculated using two-way ANOVA for tumor growth curves and log-rank (Mantel-Cox) tests for Kaplan-Meier analyses.

In both LTL331R and LTL610 NEPC PDX models, TVB-2640 monotherapy had no appreciable anti-tumor effect, whereas combination treatment with ESK981 and TVB-2640 significantly delayed tumor progression, reduced tumor burden, and prolonged tumor tripling/doubling-free survival (**Figures 7E, 7F, S17D, and S17E**). Importantly, body weights remained stable over the treatment period (**Figure S17D and S17E**), indicating that the combination regimen was well tolerated.

To evaluate whether this therapeutic strategy extends beyond NEPC, we first performed *in vitro* combination testing in metabolically stressed tumor models. Across multiple PDAC and gastric cancer (GC) cell lines, TVB-2640 and PIKfyve co-inhibition consistently produced strong synergistic suppression of cell growth (**Figures S17F-I**). We then performed a series of *in vivo* studies evaluating this co-inhibition strategy targeting PIKfyve and FASN. A strong synergistic anti-tumor effect was observed in the pNET BON-1 CDX model (**Figures 7G and S18A**) and the murine PDAC 7940B model in an immunocompetent host (**Figures 7H and S18B**). No significant impact on body weight was observed in either model. Collectively, these data support dual inhibition of PIKfyve and FASN as a therapeutic strategy in tumors where PIKfyve-dependent stress signaling drives reliance on adaptive lipogenic processes.

### PIKfyve inhibition synergizes with standard of care therapy in NEPC

Cisplatin-based chemotherapies are a first-line treatment for NEPC, but their clinical benefit is typically limited by short-lived responses^4^. We therefore examined whether PIKfyve inhibition could potentiate cisplatin-based therapy in NEPC. In both LTL545 and LTL610 NEPC PDX models, cisplatin monotherapy produced only partial tumor growth inhibition. In contrast, combined ESK981 and cisplatin treatment markedly prolonged progression-free survival (**Figures 7I, 7J, S18C, and S18D**). Consistent with these growth kinetics, all tumors in the LTL545 PDX model achieved at least stable disease following combination treatment, with four out of twelve tumors achieving complete responses based on RECIST criteria (**Figure 7I**). Notably, all nine tumors in the LTL610 PDX model treated with ESK981 and cisplatin achieved complete responses (**Figure 7J**). Across all treatment groups, body weight changes remained within a tolerable range (**Figures S18C and S18D**), indicating that the combination regimen was well tolerated. Together, these findings demonstrate that PIKfyve inhibition markedly enhances the efficacy and durability of cisplatin-based chemotherapy in NEPC, underscoring the therapeutic potential of PIKfyve inhibition-based combination strategies.

Next, we systematically evaluated the acute and chronic toxicity profiles of combined PIKfyve and FASN inhibition as well as combined PIKfyve and cisplatin treatment in outbred CD-1 male mice at PD5 and PD30 timepoints. All treatments were well tolerated, with no significant body weight loss throughout the treatment course (**Figure S19A**). Complete blood count (CBC), liver, and kidney function were analyzed at PD5 and PD30, with no signs of detrimental effects from either combination treatment (**Figures S19B-D**). Histopathological evaluation of visceral organs (heart, lung, liver, spleen, kidney, small intestine), male reproductive organs (prostate, testes), and hematopoietic tissue (bone marrow) revealed no treatment-related adverse effects (**Figure S19E**). Collectively, these data confirm that the ESK981 and TVB-2640 combination and the ESK981 and cisplatin combination are well tolerated with a favorable safety profile in immune competent CD-1 male mice.

## Discussion

In this study, we identify that the lipid kinase PIKfyve is overexpressed in NEPC and is a central mediator of ER stress adaptation, supporting tumor cell survival by orchestrating lysosome-dependent autophagy and SREBP-mediated lipogenesis. We demonstrate that NEPC tumors are characterized by predominant activation of the PERK-ATF4 branch of the ER stress response and by elevated autophagic activity. PIKfyve inhibition disrupts autophagic flux and lysosomal cholesterol trafficking, triggering compensatory activation of *de novo* lipogenic programs through SREBP activation. This reciprocal metabolic reprogramming creates a synthetic vulnerability, whereby dual inhibition of PIKfyve and lipogenesis drives accumulation of unresolved ER stress and apoptotic cell death (**Figure 8**).

**Figure 8.**
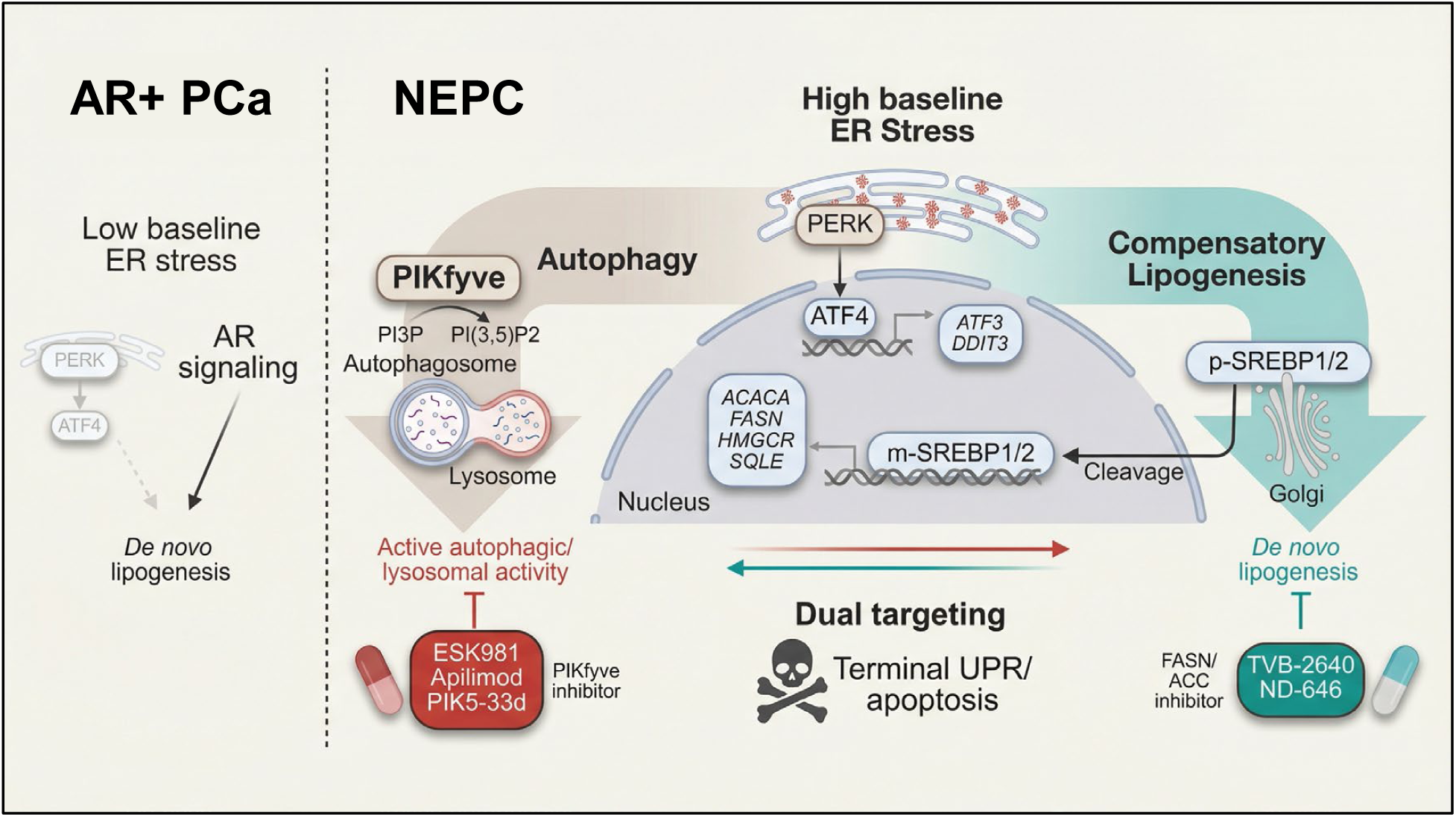
Stress-lipid kinase axis in NEPC and its therapeutic disruption. NEPC exhibits elevated baseline ER stress with predominant activation of the PERK-ATF4 pathway, whereas AR+ prostate cancer is characterized by AR-driven *de novo* lipogenesis and relatively low basal ER stress. Under this stress-adapted state, the lipid kinase PIKfyve supports lysosome-dependent autophagic flux through generation of PI(3,5)P_2,_ enabling cellular adaptation and survival. Inhibition of PIKfyve (via apilimod, PIK5-33d, or ESK981) impairs autophagy and redirects the stress response toward SREBP1/2-dependent compensatory lipogenesis, forming an ER stress-lipogenesis axis. Co-inhibition of PIKfyve and fatty acid synthesis (via TVB-2640 or ND-646) collapses both adaptive arms, amplifying ER stress and triggering the terminal unfolded protein response (UPR) and apoptosis.

The role of PIKfyve in stress adaptation has been previously linked to osmotic and nitrogen stress^63,64^, and a recent study in melanoma reported PIKfyve inhibition induces a lineage-specific induction of ER stress-mediated apoptosis via IL24^38^. Notably, Cheng et al.^23^ recently demonstrated in PDAC that PIKfyve inhibition induces a compensatory SREBP-driven lipogenic program, supporting the broader concept that PIKfyve regulates lipid metabolic adaptation beyond its canonical role in endolysosomal biology. Our findings are consistent with this emerging concept but define a distinct, tumor-intrinsic stress adaptation in NEPC. Whereas adaptive lipogenesis in PDAC is shaped by KRAS-MAPK signaling and exploited through MEK co-targeting, NEPC is characterized by PERK-ATF4-dominant stress adaptation and is sensitized by direct inhibition of the lipogenic machinery through FASN blockade. Thus, our study extends PIKfyve-dependent lipid adaptation to a KRAS-independent neuroendocrine malignancy and links this axis to ER stress buffering, autophagy, and apoptotic vulnerability in NEPC. Notably, the involvement of PERK-ATF4 signaling mirrors observations in PTEN-deficient, MYC-overexpressing prostate cancer models, where this pathway promotes cytoprotective autophagy and tumor progression under stress^65,66^.

Among the three UPR branches, our data support predominant PERK-ATF4 activation, with additional IRE1α engagement upon co-targeting PIKfyve and lipogenesis, whereas ATF6 was not further activated under the tested conditions. PERK and IRE1α can respond to membrane lipid perturbation in addition to proteotoxic stress^40^, whereas ATF6 is activated primarily through ER proteostasis stress involving BiP dissociation, ER-to-Golgi trafficking, and site-1/site-2 proteolytic cleavage^7^. Although ATF6 and SREBPs both undergo Golgi-dependent proteolytic processing, SREBPs are regulated by SCAP/INSIG-dependent sterol and lipid sensing^47^. Thus, shared processing machinery does not imply coordinated ATF6 and SREBP activation, consistent with robust SREBP activation without substantial ATF6 engagement in our system.

Our data further delineate a metabolic divergence between NEPC and AR+ prostate cancer. AR+ prostate cancer is intrinsically reliant on *de novo* lipogenesis driven by AR-mediated activation of SREBPs^67–69^, and inhibition of FASN in these tumors induces ER stress and tumor suppression^68^. Conversely, NEPC tumors, which have lost AR signaling, rely primarily on PIKfyve-dependent autophagy for ER stress resolution. This mechanistic distinction may explain the limited efficacy of ESK981 in two Phase II trials conducted exclusively in AR+ metastatic CRPC patients^30,31^, where compensatory lipogenesis may blunt the impact of PIKfyve inhibition. Our findings thus provide a strong biological rationale for prioritizing PIKfyve-targeted therapies in NEPC. Supporting this, our preclinical models show that ESK981, unlike the earlier-generation PIKfyve inhibitor apilimod, consistently induced robust apoptosis and durable tumor regression in NEPC. A Phase II trial (NCT05988918) in NEPC patients evaluating the therapeutic potential of ESK981 is ongoing.

Moreover, we show that dual inhibition of PIKfyve and multiple lipogenic enzymes, including FASN, ACC, SCD1, HMGCR, FDFT1, and SQLE, induces a robust synthetic vulnerability in NEPC by concurrently disabling autophagic clearance and lipid biosynthesis. Despite compelling preclinical evidence, no clinical trials to date have evaluated co-targeting autophagy and lipogenesis in any cancer type. Current clinical efforts remain focused on single agents, such as hydroxychloroquine or TVB-2640, or their use in combination with conventional chemotherapeutics^70,71^. Tumors such as PDAC^72^, melanoma^73^, glioblastoma^74^, and colorectal cancer^75^ commonly exploit autophagy as a resistance mechanism, while secretory neoplasms such as multiple myeloma^10^ and pancreatic neuroendocrine tumors^76^ rely heavily on UPR signaling. These ER stress-adapted malignancies may be broadly susceptible to therapeutic disruption of the stress-lipid kinase axis through concurrent targeting of PIKfyve and lipogenesis.

Given the potent synthetic vulnerability observed with dual PIKfyve and FASN inhibition in our preclinical NEPC models, further clinical trials evaluating this combination are warranted. Although our *in vivo* studies were conducted in immunodeficient hosts, prior work shows that PIKfyve inhibition enhances antigen presentation and anti-tumor immunity^22,28,29^, suggesting that therapeutic benefit may be further amplified in immune-competent settings or in combination with immunotherapy. Moreover, our *in vivo* data demonstrate that combined PIKfyve inhibition and cisplatin synergistically suppresses tumor growth in NEPC, providing direct preclinical evidence for this combination strategy. This synergy likely reflects the ability of PIKfyve inhibition to disrupt lysosomal stress resolution, thereby sensitizing tumor cells to cisplatin-induced DNA damage and apoptosis. A sequential treatment strategy, alternating targeted inhibition of PIKfyve and FASN with platinum-based chemotherapy, may therefore prolong responses, delay resistance, and potentially achieve durable control or cures in NEPC.

In conclusion, we define PIKfyve as a context-dependent vulnerability in NEPC, functioning as a central stress mediator that links autophagy and lipogenesis to ER stress adaptation. Under constitutive or exacerbated ER stress, PIKfyve becomes essential, and its combined inhibition with compensatory lipogenesis blockade induces a synthetic vulnerability. These findings establish the stress-lipid kinase axis as a central, translationally actionable target in NEPC, with potential relevance to other ER stress-driven malignancies.

## Limitations of the study

This study primarily focuses on preclinical models of NEPC and other ER stress-adapted malignancies and thus does not capture the full spectrum of inter-patient heterogeneity in stress dependency or therapeutic response. In addition, while our *in vivo* efficacy studies demonstrate durable anti-tumor activity and favorable tolerability, the long-term consequences of sustained PIKfyve inhibition and optimal clinical sequencing with metabolic or cytotoxic therapies remain to be defined in clinical settings. Moreover, the PRISM screening-derived relationships between metabolites and drug response are associative, and experimental validation is required to establish causality. Although our data support predominant PERK-ATF4 activation following PIKfyve inhibition, the precise mechanism underlying preferential PERK engagement was not fully elucidated in this study. Future studies will be needed to define how PIKfyve-dependent lysosomal dysfunction and lipid trafficking changes are transmitted to the ER to activate PERK. Finally, although our data support a stress-sensitizing role for PIKfyve inhibition and our previous studies showed PIKfyve inhibition enhances antigen presentation and anti-tumor immunity^22,28,29^, the contribution of immune-mediated effects to therapeutic response warrants further investigation.

## Resource availability

### Lead contact

Further information and requests for resources and reagents should be directed to and will be fulfilled by the lead contact, Arul M. Chinnaiyan.

### Materials availability

Correspondence and requests for materials should be addressed to Arul M. Chinnaiyan. All biological materials can be obtained from the corresponding authors following reasonable request.

### Data Availability

- RNA sequencing data generated in this study are deposited in the Gene Expression Omnibus (GEO) database (RRID: SCR_005012) under accession number GSE295392. The mass spectrometry proteomics data have been deposited to the ProteomeXchange Consortium via the PRIDE^77^ partner repository with the dataset identifier PXD076640. The metabolomics data generated in this study have been deposited in the NIH Common Fund’s National Metabolomics Data Repository, Metabolomics Workbench^78^, under Study ID ST004830 for whole-cell lipidomics and ST004831 for Lyso-IP lipidomics. The metabolomics data are accessible through the Project DOI: http://dx.doi.org/10.21228/M82S0F.
- Processed data supporting the findings of this study are available from the corresponding author upon request.
- All other relevant data are included in the manuscript and its supplementary files. No custom code was generated or used in this study.

## Acknowledgements

We gratefully acknowledge Kyle Garcia-Rogers, Arya Kamat, Rowena Kannaiyan, Adrij Mohan, Sarah N. Yee, Jacinda Liu, Ahmet Korkaya, Lisa McMurry, Fengyun Su, Rui Wang, Amanda Miller, Christine Caldwell-Smith, Xia Jiang, Shannon VanAken, Shuqin Li, Yunhui Cheng, Jean Tien, Yi Bao, Fan Yang, and Brian Magnuson from the Michigan Center for Translational Pathology at the University of Michigan for providing technical assistance. We also thank Thekkelnaycke Rajendiran, Tanu Soni, and Tushar Thoppae for their contributions to lipidomics data processing; Emily J. Bristow for assistance with phosphoinositide lipid measurements; and the Proteomics Resource Facility (RRID: SCR_026723) for mass spectrometry-based proteomics analyses. We acknowledge the NIH Common Fund’s National Metabolomics Data Repository, Metabolomics Workbench, supported by NIH grants U2C-DK119886 and OT2-OD030544, for hosting the metabolomics data associated with this study. This work was supported by the following mechanisms: National Cancer Institute (NCI) Outstanding Investigator Award R35-CA231996 (A.M.C.), NCI Early Detection Research Network U2C-CA271854 (A.M.C.), the Michigan Prostate SPORE P50-CA186786 (A.M.C.), Department of Defense Idea Development Award HT9425-23-1-0084 (Y.Q.), Neuroendocrine Research Foundation Investigator Award (Y.Q.), and National Institute of Neurological Diseases and Stroke (NINDS) R01 1NS129198 (L.S.W.). C.A.L. was supported by the NCI (R37-CA237421, R01-CA248160, R01-CA244931). C.C. was supported by an NCI F30 fellowship (F30CA288093) and NIH T32 training grants (CMB: 5T32-GM145470, MSTP: T32GM00786). A.M.C. is a Howard Hughes Medical Institute Investigator, A. Alfred Taubman Scholar, and American Cancer Society Professor.

## Author contributions

Y.Zheng, Y.Q., and A.M.C. designed and conceived the study; Y.Zheng and Y.Q. performed *in vitro* and *in vivo* experiments with assistance from C.C., Y.C., Y.Y., Y.Zhao, and W.L.; G.C., and Y.Zhang conducted RNA-seq data processing and statistical analysis; R.B. participated in statistical analysis of proteomics experiments; S.M. and Y.Zheng carried out the immunofluorescent staining; R.P., S.M., J.H., and R.M. obtained clinical tissue sections, performed immunohistochemistry and RNA-ISH staining, and histopathological interpretation. H.K. and L.S.W. performed phosphorylated phosphoinositide lipid labeling and quantification; C.L., Z.W., and K.D. synthesized PIKfyve degrader PIK5-33d; X.C. supported sequencing data processing; U.V., V.S., and C.A.L. assisted with manuscript organization; H.X. and Y.W. provided NEPC patient-derived xenografts; Y.Q. and A.M.C. provided resources and funding. Y.Zheng, Y.Q., S.J.M., and A.M.C. wrote the manuscript and organized the final figures. All authors discussed the results and participated in manuscript review.

## STAR METHODS

### EXPERIMENTAL MODEL AND STUDY PARTICIPANT DETAILS

#### Patient data

Five NEPC patient needle biopsies containing both tumor and adjacent benign prostate tissues were obtained from the Department of Pathology at the University of Michigan. In addition, the NEPC case WA76 was obtained through the Michigan Legacy Tissue Program at the University of Michigan. These samples were used for IHC and RNA-ISH staining to assess the expression of targets of interest in benign versus tumor regions. The use of archived human specimens was approved by the University of Michigan Institutional Review Board and did not require informed consent in accordance with institutional guidelines.

#### Cell lines and cell culture

LNCaP (RRID: CVCL_0395), VCaP (RRID: CVCL_2235), PC3 (RRID: CVCL_0035), DU145 (RRID: CVCL_0105), NCI-H660 (RRID: CVCL_1576), A375 (RRID: CVCL_0132), PANC-1 (RRID: CVCL_0480), HPAC (RRID: CVCL_3517), SW1990 (RRID: CVCL_1723), HPAF-II (RRID: CVCL_0313), MIA PaCa-2 (RRID: CVCL_0428), and AGS (RRID: CVCL_0139) cell lines were purchased from the American Type Culture Collection. BON-1 cells (RRID: CVCL_3985) were purchased from Creative Biolabs. The murine PDAC cell line 7940B was provided by Greggory Beatty (University of Pennsylvania). LTL331R-CL and LTL610-CL were derived from NEPC patient derived xenografts (PDXs) LTL331R and LTL610, respectively. NEPC cell lines, including NCI-H660, LTL331R-CL, and LTL610-CL, were maintained in RPMI-1640 (Invitrogen, A1049101), supplemented with 1X insulin-transferrin-selenium (Gibco, 41400045), 10 nM hydrocortisone (Sigma-Aldrich, H0888), 10 nM beta-estradiol (Sigma-Aldrich, E2758), extra 2 mM Glutamax (Gibco, 35050061), and 5% fetal bovine serum (FBS, Cytiva, SH30071.03). Serum starvation for NEPC cells was performed by reducing the FBS concentration from 5% to 0.1%. LNCaP and PC3 cells are cultured in RPMI-1640 (Invitrogen, A1049101), VCaP and A375 in DMEM (Gibco, 10566016), and DU145 in EMEM (Fisher, 30-2003), all supplemented with 10% fetal bovine serum (FBS). All other cell lines were maintained under standard culture conditions recommended by the vendor. All cells were grown in 5% CO_2_ incubators at 37°C, regularly checked for mycoplasma contamination, and authenticated by genotyping.

#### Mouse studies

All animal experiments were approved by the University of Michigan Institutional Animal Care and Use Committee. Six- to eight-week-old male CB17 severe combined immunodeficient (SCID) mice were used for all prostate cancer models and for the BON-1 pNET model. C57BL/6 female mice were used for the 7940B PDAC syngeneic model. CD-1 male mice were used for systemic toxicity assessments. At the indicated time points or study endpoints, tumors were harvested, weighed, imaged, and either flash-frozen or fixed in formalin for downstream protein, DNA, and RNA analyses or FFPE embedding. For **cell line-derived xenograft (CDX) models**, tumor cells were suspended in serum-free medium with 50% Matrigel (BD Biosciences) and injected subcutaneously into both flanks. Tumors were established in SCID mice using VCaP (3 × 10^6^/site), LNCaP and DU145 (1 × 10^6^/site), NCI-H660 (5 × 10^6^/site), and BON-1 (2 × 10^6^/site). For the PDAC syngeneic model, 7940B cells (0.5 × 10^6^/site) were implanted into C57BL/6 mice. Tumor volumes were measured at least twice weekly using digital calipers and calculated as (π/6) × (length × width²). Once the average tumor volume reached 100-200 mm³ (or approximately 50 mm³ for the 7940B model), mice were randomized into treatment groups and received vehicle, single-agent compounds, or combination therapies as indicated. In CRPC VCaP models, mice were surgically castrated upon tumors reaching approximately 200 mm³, and treatment was initiated following tumor regrowth to pre-castration size. Kaplan-Meier analyses for NEPC efficacy studies were standardized using freedom from ≥ 2× baseline tumor volume as the event criterion to enable consistent comparison across NEPC treatment groups and models. For non-NEPC models, event thresholds were defined based on model-specific tumor growth kinetics: freedom from ≥ 4× baseline tumor volume was used for the BON-1 pNET model, and freedom from ≥ 10× baseline tumor volume was used for the 7940B PDAC model. Thresholds were defined prior to analysis and are indicated in respective figure legends.

For **human patient-derived xenograft (PDX) models**, MDA-PCa-146-12 (AR+ CRPC) and MDA-PCa-146-10 (NEPC) were acquired from the University of Texas MD Anderson Cancer Center^79^; the PC310 PDX model was obtained from Erasmus Medical Center (Rotterdam, Netherlands); and NEPC PDX models LTL331R, LTL352, LTL545, and LTL610 were obtained from the Living Tumor Laboratory^80^. PDXs were maintained in intact male SCID mice and passaged by implanting 2-5 mm³ tumor chunk into the dorsal flanks subcutaneously. When tumors reached 100-200 mm³, mice were randomized into treatment groups similarly to CDX models.

For the **systemic toxicity assessment model**, male CD-1 mice were treated with single-agent or combination regimens of ESK981 + TVB-2640 or ESK981 + cisplatin. Acute and chronic toxicity was evaluated at post-dose day 5 (PD5) and day 30 (PD30), respectively, with monitoring of body weight, hematologic parameters, serum chemistry, and histopathologic assessment of major organs.

### METHOD DETAILS

#### Compounds

ESK981 was provided by Esanik Therapeutics. Apilimod, Torin-1, bafilomycin A1, simvastatin, atorvastatin, WX8, MK-8245, ND-646, and Q-VD-OPh were purchased from Selleck Chemicals. PIK5-33d was provided by K. Ding. U18666A, YM-53601, and NB-598 were obtained from MedChemExpress. TVB-2640 was purchased from Nantong Hi-future Biotechnology. Doxycycline hydrochloride was obtained from Fisher Scientific. Thapsigargin was purchased from Cell Signaling Technology. All compounds used in this study were prepared based on instructions from manufacturers, and concentrations are stated in corresponding Methods or figure legends. A list of compounds used in the study can be found in the STAR Methods.

#### Cellular thermal shift assay (CETSA)

The ability of compounds to interact with and stabilize the target protein in intact cells was analyzed as described previously^81^. Briefly, LTL331R-CL cells were seeded overnight and then treated with either DMSO, ESK981 (1 μM), or apilimod (1 μM) for 3 hours at 37°C and 5% CO_2_. Subsequently, single cell suspensions were obtained by trypsinization. One million single cells per sample were suspended in 50 μL PBS. The cell suspensions were incubated in a PCR thermal cycler at various temperatures for two cycles, each consisting of 3 minutes of heating followed by 3 minutes of cooling at room temperature (RT). Cells were lysed using three cycles of freeze-thawing with liquid nitrogen and a 25°C water bath. The lysed cells were then centrifuged and mixed with sample loading dye. A 15 μL aliquot of the sample was subjected to analysis by immunoblotting. CETSA was similarly performed in LNCaP and DU145 cells, with the treatment of either DMSO, ESK981 (0.3 μM), or apilimod (0.3 μM) for 2 hours.

#### Western blot

Cell lysates were prepared in Pierce RIPA buffer (ThermoFisher, 89901) supplemented with halt protease and phosphatase inhibitor cocktail (ThermoFisher 78441). Protein concentrations were determined using the Pierce 660nm Protein Assay Reagent (ThermoFisher 22660). 15 μg of whole cell lysates were loaded onto NuPAGE 4-12% Bis-Tris Midi gels (Invitrogen, WG1403BX10) and transferred to 0.45 μm PVDF membrane (Immobilon, IPVH20200) using a Trans-Blot Turbo transfer system (Bio-Rad, 1704150). The membranes were blocked with 5% non-fat dry milk in Tris-buffered saline with 0.1% Tween 20 (TBS-T), followed by incubation with primary antibodies for 1 hour at RT and overnight at 4°C. Membranes were washed in TBS-T and incubated with corresponding HRP-conjugated secondary antibodies for 1 hour. Protein signals were visualized using SuperSignal West Femto Maximum Sensitivity Substrate (ThermoFisher, 34095) and captured with the Odyssey Fc imaging system (LI-COR Biosciences). A list of antibodies used in the study can be found in the STAR Methods.

#### Nuclear fractionation of SREBP1

Nuclear and cytoplasmic fractions were prepared using an optimized NE-PER (Thermo Fisher Scientific, 78833) based protocol for NEPC cells. Briefly, NCI-H660 and LTL331R-CL cells (10 × 10⁶ cells per condition) were treated with DMSO, apilimod (1 µM), or PIK5-33d (0.3 µM) for the indicated time points. Cells were harvested by centrifugation (500 × g, 5 min, 4 °C), washed once with ice-cold PBS, and lysed in Cytoplasmic Extraction Reagent I (CER I) freshly supplemented with protease/phosphatase inhibitors and 0.5 mM DTT. After incubation on ice for 10 min, CER II was added, followed by brief incubation and centrifugation (16,000 × g, 5 min, 4 °C) to collect the cytoplasmic fraction. The nuclear pellet was washed once with PBS containing inhibitors and MgCl₂, then resuspended in Nuclear Extraction Reagent (NER) supplemented with protease/phosphatase inhibitors, 0.5 mM DTT, MgCl₂, and benzonase (250 U/mL). Samples were incubated on ice for 30 min with gentle mixing, followed by centrifugation (16,000 × g, 10 min, 4 °C) to obtain the nuclear fraction. Fraction purity was assessed by immunoblotting using histone H3 as a nuclear marker and GAPDH as cytoplasmic markers.

#### CytoID autophagosome accumulation assay

Autophagosome accumulation was assessed using the CytoID® Autophagy Detection Kit (Enzo Life Sciences, ENZ-KIT175) according to the manufacturer’s instructions. Briefly, cells were plated in poly-D-lysine-coated 96-well plates and treated with the indicated compounds for 24 hours. Cells were then incubated with CytoID Green detection reagent and Hoechst 33342 in staining buffer for 30 minutes at 37 °C, protected from light. After washing, fluorescence was measured immediately using a microplate reader, with CytoID signal detected in the FITC channel and Hoechst signal used for nuclear normalization. CytoID fluorescence intensity was normalized to Hoechst signal to account for differences in cell number.

#### TUNEL *in situ* cell death assay

Apoptosis was quantified in formalin-fixed paraffin-embedded (FFPE) tissue sections using the Terminal deoxynucleotidyl transferase dUTP Nick End Labeling (TUNEL) assay (*In Situ* Cell Death Detection Kit, TMR red, Roche, 12156792910). Tissue sections were first deparaffinized using sequential xylene and graded ethanol washes, followed by digestion with 20 µg/mL proteinase K for 30 mins (QIAGEN, 19133). Sections were incubated with the TUNEL reaction mixture at 37°C for 1 hour. Nuclei were counterstained with 5 µg/mL DAPI for 10 mins (Sigma-Aldrich, D9542). Slides were mounted and imaged using a Zeiss Axio Imager M1 microscope using the following channels: DAPI: 405 nm channel; TUNEL: 568 nm channel. For quantitative analysis, images were scanned using the EVOS M7000 imaging system (ThermoFisher). Fiji/ImageJ (v2.3.0) was used for automated image thresholding and quantification. TUNEL-positive area was normalized to the DAPI-positive area. Images were coded prior to quantification to mask treatment identity.

#### *In vivo* compound formulation and administration

ESK981 was formulated at 7.5 mg/mL in Ora-Plus (Perrigo, 0-30574-030316), vortexed, and sonicated until a homogeneous suspension was achieved. Apilimod and TVB-2640 were similarly prepared at 15 mg/mL and 6.25 mg/mL, respectively. Cisplatin was dissolved in normal saline. All compounds except cisplatin were administered by oral gavage at 4 μL/g body weight to reach final dosages: ESK981 at 30 mg/kg (exception: 20 mg/kg for the BON-1 model), apilimod at 60 mg/kg, and TVB-2640 at 25 mg/kg, once daily, five days per week. Gavage was performed according to the prescribed dosing schedule. Cisplatin was administered intraperitoneally once per week at 2 mg/kg for the LTL545 model, 0.25 mg/kg for the LTL610 model (with one higher dose of 2 mg/kg administered as indicated), and 2 mg/kg for CD-1 mice. Due to the distinct physical appearance of ESK981 (yellow) and vehicle (white), investigators were not blinded to treatment allocation during drug administration. However, tumor measurements and downstream quantitative analyses were performed in a blinded manner to minimize bias during outcome assessment.

#### Establishment of *ex vivo* NEPC cell lines

LTL331R-CL and LTL610-CL NEPC cell lines were derived from LTL331R and LTL610 NEPC PDX tumors, respectively. Fresh tumor tissues were harvested and kept in cold, sterile PBS. After washing twice with PBS, tumors were minced on ice and digested in a solution containing collagenase II at 1 mg/mL (Fisher Scientific, 17101-015) and DNase I at 1 mg/mL (Sigma-Aldrich, 11284932001) at 37°C with 125 rpm shaking for 15 minutes. The resulting cell suspension was filtered sequentially through 100 µm and 40 µm strainers and centrifuged. Red blood cell lysis buffer was applied for 2 minutes at room temperature to eliminate erythrocytes. Cells were washed, and viable cells were enriched by Ficoll gradient centrifugation. For immunocytochemistry, cells were embedded in HistoGel and processed for histological (H&E) and immunohistochemical staining (AR and NE markers). Doubling time was determined using CellTiter-Glo assays. Genotyping was performed at p0, intermediate (LTL610-CL p6 and p9; LTL331R-CL p8 and p12), and late (LTL610-CL p18; LTL331R-CL p19) passages to confirm lineage and authenticity of the cell lines. Detailed genotyping results are included in **Table S1**.

#### Short hairpin RNA (shRNA)

To generate stable, doxycycline-inducible sh*PIKFYVE* knockdown cell lines, LTL331R-CL and NCI-H660 cells were infected with lentivirus derived from SMARTvector lentiviral shRNA constructs targeting *PIKFYVE* (Dharmacon, V3SH11252-225467980) or a non-targeting (NT) (Dharmacon, VSC11651) control sequence. Following lentiviral infection, transduced cells were selected with 1 µg/ml puromycin. After one week of puromycin selection, 1 µg/ml doxycycline was added to induce shRNA expression. For *in vivo* studies, LTL331R-CL sh*PIKFYVE* and NCI-H660 sh*PIKFYVE* cells (5 × 10⁶ cells/site) were subcutaneously injected into male SCID mice. After tumor volume reached 100-200mm^3^, tumor-bearing mice were randomized to two groups, where the vehicle group received regular chow and the doxycycline group was fed with doxycycline-containing chow (doxycycline hyclate, 625 mg/kg diet; Envigo, TD.09651) to induce gene knockdown. Tumor volumes were measured at least twice weekly using digital calipers and calculated as (π/6) × (length × width²).

#### Immunohistochemistry (IHC)

IHC staining was performed on 5 µm FFPE tissue sections using the Ventana Discovery ULTRA automated system (Roche Diagnostics). Sections were first baked at 60°C, deparaffinized with Discovery Wash solution (Roche, 950-510) at 75 °C, and subjected to antigen retrieval using Discovery CC1 buffer (Roche, 06414575001) at 95°C. Endogenous peroxidase activity was blocked with Discovery Inhibitor CM (Roche, 760-4840) for 12 minutes at 37°C. Slides were then incubated at 37°C with primary antibodies specific to PIKfyve (Rabbit polyclonal; Proteintech, 13361-1-AP, RRID: AB_10638310), AR (Rabbit monoclonal; Roche, 760-4605, RRID: AB_2921271), SYP (Rabbit monoclonal; Roche, 790-4407, RRID: AB_2336016), Ki-67 (Rabbit monoclonal; Roche, 790-4286, RRID: AB_2631262), ATF3 (Rabbit polyclonal; Sigma-Aldrich, HPA001562, RRID: AB_1078233), p62 (Rabbit monoclonal; Abclonal, A19700, RRID: AB_2862742), and LC3A/B (Rabbit monoclonal; Cell Signaling Technology, 12741S, RRID: AB_2617131). Detection utilized the OmniMap Anti-Rabbit HRP system (Roche, 760-4311, RRID: AB_2811043), followed by chromogenic development with Discovery ChromoMap DAB (Roche, 760-159). Counterstaining was achieved with Hematoxylin II (Roche, 790-2208) for 12 minutes and Bluing Reagent (Roche, 760-2037) for 8 minutes. Slides were then washed, manually dehydrated through ethanol/xylene gradients, and coverslipped with DPX mounting medium. For each antibody, both positive and negative contextual tissue controls were included to validate specificity and exclude non-specific signals. Scoring was based on subcellular localization: nuclear for AR, Ki-67, and ATF3, and cytoplasmic for PIKfyve, p62, LC3A/B, and SYP. For all antibodies except Ki-67 (which was scored based on the percentage of positive tumor cells), a semi-quantitative product-score was used: H-score = (1 × % weakly stained cells) + (2 × % moderately stained cells) + (3 × % strongly stained cells), with staining intensities defined as 0 (no staining), 1+ (weak), 2+ (moderate), and 3+ (strong). Internal controls and tissue controls were critical for interpreting staining quality; samples lacking detectable staining in tumor, stromal, or immune compartments (as with PIKfyve) were excluded from analysis.

#### Immunofluorescence (IF)

IF was performed for ATF4 (Proteintech, 10835-1-AP) on 5-μm FFPE tissue sections using the Ventana ULTRA automated slide staining system (Roche-Ventana Medical Systems) with the anti-rabbit OmniView HRP (760-4311, Roche-Ventana) and Cy5 Tyramide/Amplification reagent (760-238, Roche-Ventana) to develop a fluorescent signature, followed by DAPI (Prolong gold anti-fade, P36931, Thermo Fischer Scientific) for nuclear counterstaining. Slides were imaged using an EVOS 7000 system (Thermo Fisher Scientific). IF staining was independently evaluated by two study pathologists (R.P. and R.M.) at ×100 and ×200 magnification to assess biomarker expression and quantify positive signals.

#### RNA *in situ* hybridization (RNA-ISH)

RNA-ISH was conducted manually on 5 µm FFPE human tissue sections using the RNAscope® 2.5 HD Brown kit (Advanced Cell Diagnostics [ACD], 322300) with a probe specific to *PIKFYVE* (ACD, 1326631-C1), following protocols described previously^82,83^. Quality of RNA was confirmed using a positive control probe for the housekeeping gene PPIB (ACD, 313901), and assay background was assessed using a negative control probe against the bacterial gene DapB (ACD, 310043). The procedure included deparaffinization, endogenous peroxidase blocking with hydrogen peroxide, heat-induced epitope retrieval using retrieval buffer in a steamer, and protease digestion to allow probe access. Probe hybridization was carried out in a HybEZ II oven (ACD, 321710) at 40°C for 2 hours, followed by a multi-step signal amplification using Amp1-6 reagents, culminating in HRP detection and DAB chromogen development to visualize RNA puncta under bright-field microscopy. Scoring was performed by study pathologists (R.M., R.P., and J.H.) blinded to clinical data, using both surgical specimens and TMAs. Only tissue samples or TMA spots with detectable PPIB signal were included; those lacking PPIB signal were excluded. Semi-quantitative scoring was done using a validated H-score system, calculated as H-score = (A% × 0) + (B% × 1) + (C% × 2) + (D% × 3) + (E% × 4), where A-E represent the proportion of cells with increasing levels of RNA signal (from none to >15 dots or clusters). The score ranges from 0 to 400, with higher values indicating higher RNA expression.

#### Combined histology score calculation

The combined histology score was derived by summing ATF4 IF-scores with ATF3 and LC3 IHC H-scores. Each marker was assigned equal weighting, and scores were summed without additional normalization to generate a single composite value per tumor. The unit of analysis was individual tumors across all preclinical models. To assess the relationship between baseline expression and treatment response, combined scores were correlated with ESK981-induced tumor growth inhibition (TGI%). Tumors were further stratified into high and low combined score groups using a combined histology score threshold of 200 and an ESK981 TGI threshold of 72%, and response separation between groups was evaluated.

#### Histopathology evaluation

For systemic toxicity assessment, major organs (liver, spleen, kidney, lung, heart, prostate, testis, small intestine, colon, and bone marrow) from CD-1 mice were harvested at necropsy 30 days post treatment to evaluate potential treatment-related toxicity. Tissues were fixed, paraffin-embedded, sectioned at 4 μm, and stained with H&E. H&E-stained sections were independently examined by study pathologists (R.P. and R.M.), who were blinded to treatment group allocation, using bright-field microscopy to assess histopathologic changes associated with drug treatment. Histopathologic findings were recorded, tabulated, and summarized across treatment groups.

#### *In vivo* hypoxia detection by Hypoxyprobe

To assess hypoxia *in vivo*, mice were intraperitoneally injected with 60 mg/kg pimonidazole hydrochloride (Hypoxyprobe™-1 Kit, Hypoxyprobe, hp12-200kit). After 4 hours, tumors were collected, fixed in formalin, and embedded in paraffin. 5 μm FFPE sections were deparaffinized, rehydrated, and stained using the anti-pimonidazole monoclonal antibody (RRID: AB_3697387) supplied in the kit, following the manufacturer’s instructions. Hypoxic regions were visualized by DAB (3,3′-diaminobenzidine) chromogenic staining.

#### *In vitro* hypoxia and pseudohypoxia treatments

For hypoxia-associated experiments, NEPC cells were cultured under normoxic, pseudohypoxic, or hypoxic conditions as indicated. Normoxic conditions were maintained at 21% O_2_. Hypoxic conditions were achieved by incubating cells in a tri-gas hypoxia chamber equilibrated to 0.1% O_2_, 5% CO_2_, and balanced N_2_. Pseudohypoxic conditions were induced by treatment with cobalt chloride (CoCl₂) at 12.5 µM under normoxic culture conditions.

#### Cell viability and synergy assays

Single cells were plated in 96-well plates, with adherent cells seeded in flat-bottom wells (Corning, 3595) and suspension cells in U-bottom wells (Corning, 353077). Depending on the cell line, seeding densities ranged from 1,000 to 30,000 cells per well in their respective culture media. After overnight incubation, serial dilutions of the indicated compounds were added to each well. Cells were then cultured for an additional seven days for prostate cancer cell lines and five days for PDAC and GC cell lines to determine compound sensitivity and half-maximal inhibitory concentration (IC_50_). Cell viability was assessed using the CellTiter-Glo luminescent cell viability assay (Promega, G7572), and luminescence was measured with the Infinite M1000 Pro plate reader (Tecan). Drug combination experiments were performed at least twice independently, with each experiment including a full dose-dose matrix and 3-6 technical replicates per condition. Synergy was evaluated using SynergyFinder (https://synergyfinder.fimm.fi, RRID: SCR_026127) with the following parameters: outlier detection enabled, curve fitting model LL4, Bliss model for synergy calculation, and correction factor applied^84^. All drug synergy analyses in this study were quantified using the Bliss independence synergy model. Bliss synergy scores were calculated as the deviation between observed combination effects and the expected effects assuming independent drug action. A Bliss score above 10 suggests a synergistic effect between the two drugs. Each drug combination condition was performed in 3 technical replicates and repeated at least 3 times.

#### Proteasome activity assay

Proteasome chymotrypsin-like activity was measured using a fluorometric AMC-based assay (Abcam, ab107921). Cell lysates were prepared in NP-40-based buffer without protease inhibitors and incubated with an AMC-conjugated proteasome substrate in the presence or absence of a proteasome inhibitor. AMC fluorescence was recorded kinetically at Ex/Em = 350/440 nm within the linear range of the reaction. Proteasome-specific activity was calculated as inhibitor-sensitive AMC release and normalized as indicated.

#### RNA isolation and quantitative real-time PCR (qPCR)

Total RNA was extracted from cultured cells or tumor tissues using the RNeasy mini kit (Qiagen, 74104), following the manufacturer’s instructions. Complementary DNA (cDNA) was synthesized from 1 μg total RNA using the Maxima First Strand cDNA Synthesis Kit (Thermo Fisher Scientific K1642). qPCR was carried out using Fast SYBR Green Master Mix (Thermo Scientific 4385612) on a QuantStudio 6 Pro Real-Time PCR system (ThermoFisher). Target gene expression was normalized to GAPDH using the ΔΔCt method. The qPCR primer sequences are listed in **Table S6**.

#### *XBP-1* mRNA splicing assay

XBP1 mRNA splicing was assessed by RT-PCR to evaluate IRE1 pathway activation. Total RNA was isolated from treated cells and reverse-transcribed using random hexamers. PCR was performed using primers flanking the 26-nt intron excised during *XBP-1* splicing (forward: 5′-CCTTGTAGTTGAGAACCAGG-3′; reverse: 5′-GGGGCTTGGTATATATGTGG-3′). PCR products corresponding to unspliced (*XBP-1*u) and spliced (*XBP-1*s) forms were resolved on 2.5-3% agarose gels and visualized by gel imaging.

#### RNA sequencing and analysis

RNA integrity was assessed using an Agilent Bioanalyzer (RNA Nano Chip). For tumor samples, ribosomal RNA was depleted using the RiboErase selection kit (Kapa Biosystems, KK8561). For cell lines and other samples, polyadenylated transcripts were enriched using Sera-Mag Oligo(dT)-Coated Magnetic Particles (GE Healthcare Life Sciences, 38152103010150). RNA libraries were constructed using the KAPA RNA HyperPrep Kit (Roche Sequencing, KK8541) from 800 ng of total RNA. Following rRNA depletion and RNA fragmentation (200-300 bp), double-stranded cDNA was synthesized, ligated with NEB adapters, and PCR-amplified using KAPA HiFi HotStart ReadyMix and NEB dual-index barcodes. Library quality and quantity were assessed using an Agilent 2100 Bioanalyzer (DNA 1000 chip). Paired-end sequencing (2 × 151 bp) was performed on an Illumina NovaSeq 6000 platform, yielding 30-40 million reads per sample. Base calling and demultiplexing were performed using bcl2fastq v2.20 (RRID: SCR_015058). Xenograft FASTQ files were separated into host and graft using xengsort^85^. Transcript quantification was performed using kallisto, and gene-level abundances were aggregated for downstream analysis. Genes with mean TPM < 1 and non-protein coding genes in all groups were excluded. Differential gene expression analysis was conducted using the limma-voom pipeline following TMM normalization via edgeR (RRID:SCR_012802).

For pathway-level interpretation, we conducted gene set enrichment analysis (GSEA) using the fgsea R package, ranking genes by log_2_ fold change. Hallmark gene sets were obtained from MSigDB, and signature gene sets for NEPC and AR signaling were retrieved from a previously published study^32^. Additional custom gene sets, including those for ER stress^86^ and ATF4 targets^33^, were used to evaluate stress-response pathways. All gene sets used are provided in **Table S7**.

#### Filipin staining and immunofluorescence

One million NEPC cells were seeded onto poly-D-lysine/laminin-coated glass coverslips (Millipore Sigma, CLS354087) in 500 µL culture medium per well in 12-well plates and incubated overnight. Following treatments, cells were fixed with 3.2% paraformaldehyde for 15 min, quenched with 125 mM glycine for 10 min, and washed twice with PBS. Fixed cells were permeabilized with 0.1% Triton X-100 for 5 min, washed three times with PBS, and blocked in 5% BSA for 1 hour at RT. Samples were incubated with anti-LAMP1 primary antibody at 1:100 dilution (Cell Signaling Technology, 9091S, RRID: AB_2687579) overnight at 4°C, washed, and then incubated with goat anti-rabbit Alexa Fluor® 594 secondary antibody at 1:1000 dilution (Jackson ImmunoResearch, 111-585-045, RRID: AB_2338062) for 1 hour at RT. After PBS washes, unesterified cholesterol was visualized with 0.1 mg/mL filipin complex for 2 hours at RT. Coverslips were mounted in PBS and imaged using a Zeiss LSM900 confocal microscope (63× oil immersion, Airyscan mode). Excitation wavelengths were 405 nm for filipin and 561 nm for LAMP1-Alexa Fluor 594. Colocalization between filipin and LAMP1 signals was quantified using Manders’ tM2 coefficient in ZEN 3.5 software (Zeiss). tM2 was calculated above the automatically determined threshold of the filipin channel, and analysis was performed on a per-field basis. *n* = 3-4 random fields. Image quantification was performed using blinded sample identities. Experiments were repeated three times. Identical analysis parameters were applied across all images per experiment.

#### Cholesterol quantification

5 × 10^6^ NEPC, 1 × 10^6^ LNCaP, and 3 × 10^6^ VCaP cells per condition were harvested, washed with cold PBS, and resuspended in 200 µL of chloroform:isopropanol:NP-40 (7:11:0.1, v/v/v). Samples were homogenized, centrifuged at 15,000 × g for 10 min, and the organic phase was transferred to a new tube. Solvent was removed by air drying at 50 °C followed by vacuum desiccation for 30 min. Dried lipid extracts were resuspended in 100 µL of Assay Buffer by sonication or vigorous vortexing. Cellular cholesterol levels were then quantified using the Amplex® Red Cholesterol Assay Kit (Thermo Fisher Scientific, A12216) according to the manufacturer’s instructions.

#### CRISPRi sgRNA generation and establishment of stable knockdown cell lines

CRISPRi sgRNAs were designed using published sgRNA libraries^87^ and cloned into a lentiviral CRISPRi sgRNA expression vector by Golden Gate assembly. Briefly, complementary sgRNA oligonucleotides were annealed and ligated into the Esp3I-digested backbone using T4 ligase. Correct sgRNA insertion was verified by Sanger sequencing following plasmid minipreparation, and sequence-validated constructs were expanded by midiprep for lentiviral production. Lentivirus was generated in HEK293T cells using standard packaging plasmids, and viral supernatants were collected and used to transduce NEPC cell lines. Following infection, cells were selected with the appropriate antibiotic to establish stable CRISPRi cell populations. Knockdown efficiency was validated by immunoblotting and/or qPCR prior to downstream functional assays.

#### Lysosome immunoprecipitation (lyso-IP)

Lysosomes were isolated as previously described^88^. Briefly, LTL331R-CL and NCI-H660 cells were stably infected with lentivirus generated from TMEM192-3×HA construct (RRID: Addgene_102930) and selected with 1 µg/mL puromycin for 5 days. Ten million LTL331R-CL TMEM192-3×HA expressing cells were treated with either DMSO, apilimod (1 µM), or PIK5-33d (0.1 µM) for 24 hours, and then collected in 1 mL ice-cold KPBS buffer (136 mM KCl, 10 mM KH_2_PO_4_, pH 7.25), and homogenized using the Omni THq homogenizer. Lysosome containing supernatants were collected after centrifugation at 1,000 × g for 3 minutes at 4°C, and lysosomes were captured by anti-HA magnetic beads (Thermo Fisher, 88837) for 10 minutes at 4 °C. Beads were washed three times with KPBS, and lysosomes were eluted overnight in KPBS supplemented with 0.1% NP-40 and protease inhibitors at 4°C. Protein concentrations were determined using the Pierce BCA Protein Assay Kit, and equal amounts of lysosomal protein were subjected to downstream analyses.

#### Untargeted lipidomics analysis

Untargeted lipidomics was performed using liquid chromatography-mass spectrometry (LC-MS/MS). Lipids were extracted from biological samples using a modified Bligh-Dyer method. Briefly, samples were spiked with a mixture of mass spectrometry-grade internal lipid standards spanning major lipid classes and extracted at room temperature (RT) using a 2:2:2 (v/v/v) mixture of methanol, water, and dichloromethane. Following phase separation, the organic layer was collected and dried under nitrogen. Dried lipid extracts were reconstituted in 100 μL of buffer B (acetonitrile/isopropanol/water, 10:85:5, v/v/v) containing 10 mM ammonium acetate prior to LC-MS analysis.

Quality control (QC) samples were generated by pooling equal aliquots of all study samples and were injected at the beginning and end of each analytical batch and after every ten injections to monitor instrument performance, analytical stability, and reproducibility of sample preparation. Additional controls included matrix-free internal standard mixtures and characterized plasma pools, which were analyzed repeatedly throughout the run sequence.

Chromatographic separation was carried out on a Shimadzu Nexera X2 UHPLC system using a Waters Acquity HSS T3 column (1.8 μm, 50 × 2.1 mm) maintained at 55 °C. Mobile phase A consisted of acetonitrile/water (40:60, v/v) with 10 mM ammonium acetate, and mobile phase B consisted of acetonitrile/water/isopropanol (10:5:85, v/v/v) with 10 mM ammonium acetate. Lipids were eluted using a linear gradient from 40% to 98% solvent B over 10 min, held at 98% B for 7 min, followed by re-equilibration. The flow rate was 0.4 mL/min, injection volume was 5 μL, and total run time was 20 min.

Mass spectrometry data were acquired on a TripleTOF 5600 mass spectrometer (AB Sciex) equipped with a DuoSpray ion source, operated in both positive and negative ionization modes. Data-dependent acquisition (DDA) was used, consisting of one full MS survey scan followed by up to 15 MS/MS scans per cycle. Dynamic background subtraction, isotope exclusion, and dynamic exclusion were enabled. Collision energy spread was applied to facilitate lipid fragmentation across the m/z range. Automated calibration was performed at the start of each batch or upon polarity switching to maintain mass accuracy.

Raw data files were converted to mgf format and searched against LipidBlast libraries using the NIST MS PepSearch program. Search parameters were optimized based on internal standards and library comparisons. Identified lipid species were compiled into an in-house reference library and quantified using MS1 signal intensity with MultiQuant software. QC samples were used to assess analytical variability and to filter lipid species below class-specific limits of quantification or with excessive missing values. Only lipid species passing QC-based filtering criteria were retained for downstream analysis.

Following QC filtering and normalization, lipidomics data were used for multivariate and comparative analyses. Principal component analysis (PCA) was performed to assess global lipidomic differences across treatment conditions using autoscaled data. For lipid class-level comparisons, summed abundances of lipid species within each class were calculated and visualized as forest plots showing log_2_ fold changes relative to control. For selected lipid subclasses, relative abundances were Z score-normalized across samples and displayed as violin plots to compare distributional shifts between treatment groups. Differentially regulated lipid species were identified based on fold-change criteria and visualized using heatmaps or ranked plots as indicated. All analyses were performed on normalized datasets, and identical processing and visualization parameters were applied within each dataset. Whole-cell and lyso-IP based lipidomic analyses were performed in NCI-H660 TMEM192-HA cells following the indicated treatments or time points. The complete quantified lipid datasets are provided in **Tables S8** and **S9**, respectively.

#### TMT mass spectrometry-based proteomics analysis

Whole-cell lysates (75 µg in 37.5 µL RIPA buffer) and lysosomal lysates (10 µg in 45 µL KPBS buffer containing 0.1% NP-40) were submitted to the University of Michigan Proteomics Core Facility for sample preparation and mass spectrometry analysis, following established protocols^89^. Briefly, protein lysates were reduced, alkylated, and digested overnight with trypsin. The resulting peptides were labeled with appropriate TMT Isobaric Labeling Reagents (Thermo Fisher Scientific) according to the manufacturer’s instructions. Three biological replicates were analyzed per condition. TMT-labeled peptide mixtures were pooled, desalted, and fractionated into 8 or 12 fractions using high-pH reversed-phase liquid chromatography to reduce sample complexity. Each fraction was analyzed by liquid chromatography-tandem mass spectrometry (LC-MS/MS) on a Thermo Scientific Orbitrap Ascend Tribrid mass spectrometer equipped with high-field asymmetry waveform ion mobility spectrometry (FAIMS) and coupled to a Vanquish Neo UHPLC system. Data acquisition was performed in MS3 mode to ensure accurate quantification of TMT reporter ions. Raw data were processed using Proteome Discoverer v3.0 (Thermo Fisher Scientific, RRID: SCR_014477) and searched against the SwissProt human protein database (Homo sapiens, TaxID 9606, v2024-10-02, 20,351 entries). Proteins and peptides were filtered at a false discovery rate (FDR) ≤1%, and quantification was based on high-quality MS3 spectra with an average signal-to-noise ratio ≥10 and <50% isolation interference. Differential protein expression analysis was performed using normalized TMT reporter ion intensities across replicates. Whole-cell and lyso-IP based proteomic analyses were performed in LTL331R-CL cells following PIK5-33d or apilimod treatment, and the complete quantified protein datasets are provided in **Tables S10-S12**.

#### Quantitation of phosphorylated phosphoinositide lipids

Phosphorylated phosphoinositide lipids were measured by live-labeling cells with myo-[2-^3^H] inositol as previously described^90^. Specifically, NCI-H660 cells were grown to 70-80% confluence on 100 mm poly-D-lysine coated dishes (Corning, 356469). Cells were rinsed twice with 1X PBS (pH 7.4) and incubated in inositol-free medium with 10 µCi/ml ^3^H-inositol (Revivity, NET114A005MC) for 48 hours. After 48 hours, the indicated small molecules were added to a final concentration of 1 µM and incubated for 4 hours. The cells were then lysed, and all macromolecules were precipitated with perchloric acid and the myo-[2-^3^H] inositol-labeled lipids were extracted and deacylated as described^90^. The resultant glycerol-phosphoinositides were separated using ion exchange chromatography on Partisphere 5 μm SAX cartridge column, 250 x 4.6 mm (WVS hardware, MAC-MOD Analytical, 4621-1505). The raw counts for each peak are presented as a percentage of the total phosphatidylinositol, which is derived from the summation of counts across the six detectable glycero-inositol peaks (PI, PI3P, PI4P, PI5P, PI(3,5)P_2_, PI(4,5)P_2_). Background scintillation counts, determined from adjacent regions, were subtracted from all peaks. Assay reproducibility was confirmed in duplicate experiments.

#### Autophagic flux quantification

To quantify autophagic flux, DU145 and PC3 cells were stably transduced with a bicistronic pMRX-IP-GFP-LC3-RFP-LC3ΔG construct (RRID: Addgene_84572) under 1 µg/mL puromycin selection^91^. For the assay, 5,000 cells were seeded per well in black wall 96-well plates and treated with test compounds for 24-48 hours depending on the experiment. GFP and RFP fluorescence signals were measured using an Infinite M1000 Pro plate reader (Tecan) at Ex/Em wavelengths of 488/510 nm and 558/583 nm, respectively. The autophagy index was calculated as a normalized ratio of GFP to RFP signal in a sample relative to the mean GFP/RFP ratio in DMSO-treated control. A decrease in the autophagy index indicates increased autophagic flux, whereas an increased autophagy index suggests inhibited autophagic flux. Assay performance was validated using Bafilomycin A1 (autophagic flux inhibitor) and Torin-1 (autophagic flux inducer) as internal controls.

#### c-Laurdan generalized polarization assay

Membrane lipid order was assessed in live cells using c-Laurdan generalized polarization (GP) imaging. Cells were seeded on poly-D-lysine-coated coverslips, treated as indicated, and stained with 10 µM c-Laurdan in phenol-red free HBSS for 30 min at 37°C protected from light. After gentle washing with warm PBS, cells were imaged immediately in FluoroBrite™ DMEM on Zeiss LSM900 confocal microscope with a 63× oil-immersion objective in Airyscan mode. c-Laurdan was excited at 405 nm, and emission was collected simultaneously in ordered and disordered membrane channels centered at 440 nm and 490 nm, respectively. GP images were calculated in Fiji using the formula GP = (I440 − I490) / (I440 + I490), and mean GP values were quantified from various regions after background subtraction. Lower GP values indicate reduced lipid packing and increased membrane fluidity.

#### Assessment of mitochondrial function and oxidative stress

Cellular oxidative stress and mitochondrial function were assessed using plate reader-based ROS-Glo™ H_2_O_2_, TMRM, and MitoSOX assays. Cells were seeded in multi-well plates and treated as indicated. For ROS-Glo analysis, H_2_O_2_ substrate was added during the final incubation period, and luminescence was measured according to the manufacturer’s protocol. For mitochondrial membrane potential and mitochondrial superoxide measurements, cells were incubated with Image-iT™ TMRM reagent or MitoSOX Red, respectively, after treatment, washed gently as needed, and fluorescence was measured using a microplate reader. Background-subtracted luminescence or fluorescence values were normalized to the corresponding DMSO control within each cell line and condition. Positive controls were included to validate assay performance, including H_2_O_2_ for ROS-Glo, FCCP for mitochondrial depolarization, and antimycin A for mitochondrial superoxide induction.

#### PRISM drug sensitivity screening and correlation analysis

High-throughput drug sensitivity screening of PIK5-33d and ESK981 was performed in collaboration with the Broad Institute using the PRISM (Profiling Relative Inhibition Simultaneously in Mixtures) platform, as previously described^92^. Compounds were screened across a pool of DNA-barcoded human cancer cell lines using an eight-point dose-response format, with threefold dilution series and highest concentration at 10 µM. Cellular viability data were used to calculate drug response metrics as log_2_-transformed area under the curve (log_2_-AUC) values for each cell line and compound. To assess pharmacologic correlations, Pearson correlation coefficients were calculated between the log_2_-AUC profiles of PIK5-33d or ESK981 and those of other compounds in the PRISM repurposing library. Statistical significance of correlations was determined using associated *p*-values, and multiple hypothesis testing was corrected using the Benjamini-Hochberg method to obtain *q*-values. For integrative analysis with metabolomics, cell line-specific log_2_-AUC values were paired with matched metabolite abundance profiles. Univariate Pearson correlations were computed between compound sensitivity and the abundance of each metabolite across cell lines. Resulting *p*-values were adjusted for multiple comparisons using the Benjamini-Hochberg method. Correlation coefficients and *q*-values for all metabolites were retained for downstream analysis and visualization. Top metabolites correlated with PRISM drug response to PIK5-33d and ESK981 are provided in **Table S13.**

#### Statistical analysis

All statistical analyses were performed using GraphPad Prism (version 10.2, RRID: SCR_002798) and R (version 4.3.2, RRID: SCR_001905), unless otherwise noted. Specific statistical tests, significance thresholds, and data presentation formats (e.g., mean ± SD/SEM) are detailed in the corresponding figure legends and relevant sections of the Methods. No power analysis was used to predetermine sample size. Group sizes were chosen based on prior publications and experimental feasibility.

## Supplementary Figure Legends

**Figure S1. PIKfyve is overexpressed in NEPC. Related to Figure 1**.

A. Representative images of PIKfyve IHC in sh*NT* and sh*PIKFYVE* tumors of LTL331R-CL NEPC CDX to validate the specificity of PIKfyve antibody.

B. Representative histological and molecular images from remaining seven metastatic sites of WA76 case (Left pelvic LN, right pelvic LN, paraaortic LN, lower aortic LN, central periaortic, common iliac LN, and femur bone marrow), showing hematoxylin and eosin (H&E) staining, RNA *in situ* hybridization (RNA-ISH), and IHC for PIKfyve in matched tumor and adjacent benign tissues.

C. Representative histology and molecular profiling of PIKfyve expression in five primary NEPC needle biopsy specimens. Shown are adjacent benign (top) and tumor (bottom) regions from the same biopsy core, stained by H&E, *PIKFYVE* RNA-ISH, and IHC for PIKfyve, AR, SYP, and Ki-67. Insets highlight higher-magnification views of RNA and protein localization.

D. Quantification of *PIKFYVE* RNA-ISH score (left) and IHC H-score (right) across five needle biopsies between tumor regions and benign tissues. *P* values calculated using a paired two-tailed student’s *t*-test.

**Figure S2. PIKfyve serves as a therapeutic target in NEPC. Related to Figure 1**.

A. Waterfall plots showing individual tumor volume changes from baseline in vehicle or ESK981 treated tumors in DU145 (AR-CDX, vehicle *n* = 18, ESK981 *n* = 14) and CRPC VCaP (AR+ CDX, vehicle *n* = 18, ESK981 *n* = 14) tumors. Data adapted from Qiao et al., *Nature Cancer*, 2021^22^.

B. Representative images of TUNEL staining showing *in situ* cell death post five days of either vehicle or ESK981 treatment in DU145 (AR-CDX) and CRPC VCaP (AR+ CDX) tumors.

C. Bright-field images showing morphology of NEPC *ex vivo* cell lines LTL331R-CL and LTL610-CL in comparison to NCI-H660.

D. Doubling time of NCI-H660, LTL331R-CL, and LTL610-CL cells *in vitro*, determined by CellTiter-Glo. Data presented as mean ± SD.

E. Body weight of mice bearing NEPC sh*PIKFYVE* CDXs during the treatment period. For NCI-H660: *n* = 3 mice per group; for LTL331R-CL: *n* = 3 (vehicle) and *n* = 4 (doxycycline). Data presented as mean ± SEM.

F. Western blot of individual tumors showing PIKfyve protein expression in NCI-H660 sh*PIKFYVE* and LTL331R-CL sh*PIKFYVE* CDXs treated with vehicle or doxycycline at day 28 and day 11, respectively.

**Figure S3. Pharmacological inhibition of PIKfyve exerts cytotoxicity in NEPC *in vivo*. Related to Figure 2**.

A. Western blot of cellular thermal shift assay (CETSA) showing PIKfyve protein stability following treatment with either DMSO, ESK981 (300 nM), or apilimod (300 nM) for 2 hours under escalating temperatures in DU145 and LNCaP cells *in vitro*.

B. Representative images showing morphological changes of vacuoles induced by PIKfyve inhibitors, ESK981 (1 µM) and apilimod (1 µM), for 8 hours in DU145 and PC3 cells, compared with DMSO.

C. Quantification of CytoID staining showing relative autophagosome accumulation index in NCI-H660 cells following 24 hours of treatment with increasing concentrations of ESK981 or apilimod. Data were normalized to the control (0 nM). Data presented as mean ± SEM. *P* values calculated using ordinary one-way ANOVA.

D. Quantification of autophagic flux in DU145 and PC3 GFP-LC3-RFP-ΔLC3 reporter cells after treatment with DMSO, ESK981 (1 µM), apilimod (1 µM), Torin-1 (250 nM), or bafilomycin A1 (BafA1, 100 nM) for 24 hours. Data presented as mean ± SD*. P* values calculated compared with DMSO group using ordinary one-way ANOVA.

E. Western blot of LC3A/B in individual tumors after vehicle or ESK981 treatment at the indicated time points in NEPC PDXs.

F. Western blot of c-PARP post vehicle, ESK981, or apilimod treatment for five days in multiple NEPC PDXs *in vivo*.

G. TUNEL *in situ* cell death assay post vehicle or ESK981 monotherapy in LTL331R NEPC PDX at the indicated time points.

H. TUNEL *in situ* cell death assay post vehicle, ESK981, or apilimod monotherapy in MDA-PCa-146-10 NEPC PDX at the indicated time points.

I. Quantification of TUNEL signal post vehicle or ESK981 monotherapy in LTL331R (left) and post vehicle, ESK981, or apilimod monotherapy in MDA-PCa-146-10 (right) NEPC PDX at different time points. Data presented as mean ± SD*. P* values calculated compared with DMSO group using a two-tailed unpaired *t*-test with Welch’s correction.

J. Individual tumor images at endpoint of NEPC models treated with either vehicle or ESK981.

K. Mouse body weight monitoring in long-term efficacy studies in different NEPC models treated with either vehicle or ESK981. For NCI-H660 and LTL545 models: *n* = 5 mice per group; for LTL331R: *n* = 6 (vehicle) and *n* = 7 (ESK981); for LTL610: *n* = 8 mice per group. Data presented as mean ± SEM.

L. qPCR results showing *CXCL10* mRNA expression in LTL331R-CL (left) and NCI-H660 (right) cells post 24 hours of DMSO, ESK981 (1 µM), IFN-g (10 ng/mL), or the combination treatment. Data presented as mean ± SD*. P* value calculated using one-way ANOVA.

M. Western blot showing MHC-I levels in various NEPC cell lines post indicated treatments for 24 hours.

**Figure S4. NEPC tumors exhibit elevated baseline ER stress and autophagy. Related to Figure 3**.

A. GSEA of NEPC signature in NEPC tumors compared with AR+ prostate tumors.

B. GSEA of AR signature in NEPC tumors compared with AR+ prostate tumors.

C. GSEA of hallmark hypoxia pathway in NEPC tumors compared with AR+ prostate tumors.

D. Representative images of HIF-1α IHC and PIMO staining in VCaP (AR+) and LTL545 (NEPC) tumors.

E. Expression of *ONECUT2* in AR+ prostate and NEPC tumor models based on RNA-seq data. Each bar represents the mean ± SD of biological replicates for each tumor type. *P* value calculated using a two-tailed unpaired *t*-test comparing all AR+ prostate with NEPC samples.

F. DEGs of IRE1 and ATF6 branches of ER stress comparing NEPC with AR+ prostate tumors by RNA-seq. The significance cutoff was set at log_2_FC greater than 2 or less than -2 and adjusted *p-*value below 0.05.

G. DEGs involved in the ATF4 target gene set in NEPC versus AR+ prostate tumors. DEGs were defined using thresholds of log_2_FC greater than 2 or less than -2 and adjusted *p-*value below 0.05.

H. qPCR results showing *ATF3* and *ATF4* mRNA expression in NEPC (*n* = 12) and AR+ prostate (*n* = 12) tumors. Data presented as mean ± SD*. P* values calculated using a two-tailed unpaired *t*-test with Welch’s correction.

I. Western blot analysis of ATF6 and XBP-1s in NEPC (*n* = 3 for each tumor type) and AR+ prostate tumors (*n* = 3 for each tumor type).

J. qPCR analysis of *XBP-1s* mRNA expression in NEPC and AR+ prostate tumors. Data presented as mean ± SEM*. P* value calculated using a two-tailed unpaired *t*-test (top). Representative DNA agarose gel showing XBP-1 mRNA splicing patterns across the corresponding tumor samples (bottom).

K. qPCR results showing *LC3B* mRNA expression in NEPC (*n* = 12) and AR+ prostate (*n* = 12) tumors. Data presented as mean ± SD*. P* value calculated using a two-tailed unpaired *t*-test.

**Figure S5. PIKfyve inhibition amplifies ER stress. Related to Figure 4**.

A. Western blot analysis of ATF4 and ATF3 in NEPC cells treated with 10 nM thapsigargin for the indicated time points.

B. qPCR of *ATF3* and *DDIT3* mRNA expression in NEPC cells treated with DMSO and thapsigargin (10 nM) for 6 hours. Data presented as mean ± SD*. P* values calculated using a two-tailed unpaired *t*-test.

C. Chemical structure of PIKfyve PROTAC degrader PIK5-33d.

D. Western blot of PIKfyve and LC3A/B in LTL331R-CL cells after increasing concentrations of PIK5-33d treatment for the indicated time points.

E. Western blot of PIKfyve and LC3A/B in NCI-H660 cells after DMSO, apilimod (1 µM), or PIK5-33d (0.3 µM) treatment for 24 hours.

F. Western blot of PIKfyve, c-PARP, ATF4, ATF3, and CHOP in NCI-H660 cells after DMSO, apilimod (1 µM), or PIK5-33d (0.3 µM), with or without thapsigargin (10 nM) treatment for 24 hours.

G. Western blot analysis of ATF4 and ATF3 in NEPC cells under serum starvation for the indicated time points.

H. qPCR of *ATF3* and *DDIT3* mRNA expression in LTL331R-CL cells treated with DMSO, apilimod (1 µM), or PIK5-33d (0.3 µM) under full serum or low serum conditions for 6 hours. Data presented as mean ± SD*. P* values calculated using two-way ANOVA.

I. Western blot of c-PARP in LTL331R-CL cells after increasing concentrations of ESK981 or apilimod treatment under full serum or low serum conditions for 24 hours.

J. Western blot of c-PARP, ATF4, ATF3, and CHOP in NCI-H660 cells after increasing concentrations of apilimod, under full serum or low serum conditions, for 24 hours.

K. qPCR of *IL24* mRNA expression following treatment with PIKfyve inhibitors (WX8, 1 µM; ESK981, 1 µM; apilimod, 1 µM; PIK5-33d, 0.3 µM) for 24 hours in A375 melanoma cells. Data presented as mean ± SD*. P* values calculated using one-way ANOVA.

L. qPCR of *IL24* mRNA expression in LTL331R-CL cells treated with indicated compounds under full serum (left) low serum (right) conditions for 24 hours. Data presented as mean ± SD*. P* values calculated using one-way ANOVA.

M. Western blot analysis of ATF4 and LC3A/B in LTL331R-CL cells treated with DMSO, ESK981 (1 µM), apilimod (1 µM), PIK5-33d (0.3 µM), bafilomycin A1 (0.1 µM), chloroquine (100 µM) under low serum conditions for 24 hours.

N. Proteasome activity in NCI-H660 and LTL331R-CL cells following treatment with DMSO, apilimod (1 µM), or bortezomib (25 nM for NCI-H660; 10 nM for LTL331R-CL) under low serum conditions for 24 hours. Proteasome activity was measured as fluorescence intensity over time and normalized to the initial time point. Data presented as mean ± SEM*. P* values calculated using two-way ANOVA.

O. Western blot analysis of total ubiquitinated proteins, K48-linked ubiquitinated proteins, LC3A/B, and ATF4 in NCI-H660 and LTL331R-CL cells following treatment with DMSO, apilimod (1 µM), or bortezomib (25 nM for NCI-H660; 10nM for LTL331R-CL) under low serum conditions for 24 hours.

**Figure S6. Predominant activation of the PERK branch of the ER stress response following PIKfyve inhibition. Related to Figure 4**.

A. DEGs associated with hypoxia and the three branches of the ER stress pathway (PERK, IRE1á, and ATF6), as well as ATF4 target genes, in LTL331R-CL cells treated with ESK981 (1 µM, top) or PIK5-33d (0.3 µM, bottom) versus DMSO under low serum conditions for 6 hours. DEGs were defined using thresholds of log_2_FC greater than 0.58 or less than -0.58 and adjusted *p-*value below 0.05.

B. qPCR of key genes in the three branches of the ER stress pathway in LTL331R-CL cells following treatment with apilimod (1 µM) or PIK5-33d (0.3 µM) for the indicated time points under low serum conditions. Data presented as mean ± SD*. P* values calculated using Brown-Forsythe and Welch ANOVA with Dunnett’s T3 multiple-comparison test.

C. qPCR of key genes in the three branches of the ER stress pathway in LTL331R-CL cells following treatment with tunicamycin (1 µg/ml) or thapsigargin (10 nM) for the indicated time points. Data presented as mean ± SD. *P* values calculated using Brown-Forsythe and Welch ANOVA with Dunnett’s T3 multiple-comparison test.

D. Representative DNA agarose gel showing *XBP-1* mRNA splicing patterns in LTL331R-CL and NCI-H660 cells following treatment with apilimod (1 µM) or PIK5-33d (0.3 µM) for the indicated time points. Thapsigargin (10 nM, 8 hours) was used as a positive control.

E. Western blot of p-PERK and PERK in LTL331R-CL cells following treatment with tunicamycin (1 µg/mL) for the indicated time points under low serum conditions.

F. Western blot of key ER stress targets in LTL331R-CL cells following treatment with tunicamycin (1 µg/mL) for the indicated time points under low serum conditions.

**Figure S7. PIKfyve inhibition alters membrane lipid packing without inducing overt oxidative or mitochondrial stress. Related to Figure 4**.

A. Representative c-Laurdan images of LTL331R-CL cells treated with DMSO, apilimod (1 µM, 24 hours), or MβCD (5 mM, 30 mins) under low serum conditions. Blue and green emission channels were collected at 440 nm and 490 nm, respectively, and used to calculate generalized polarization (GP) using the formula: GP = (I440 − I490) / (I440 + I490). Each dot represents one cell. *n* = 17 cells per condition. Data presented as mean ± SD. *P* values calculated by Brown-Forsythe and Welch ANOVA with Dunnett’s T3 multiple-comparison test.

B. Whole-cell reactive oxygen species (ROS) production measured using the ROS-Glo H_2_O_2_ assay in LTL331R-CL and NCI-H660 cells treated with DMSO, apilimod (1 µM), or PIK5-33d (0.3 µM) under low-serum conditions for 24 hours. ROS-Glo luminescence was normalized to the corresponding control for each cell line and serum condition. H_2_O_2_ (25 µM, 2 hours) was included as a positive control for increased ROS-Glo signal. Data presented as mean ± SEM. *P* values calculated using ordinary two-way ANOVA.

C. Relative mitochondrial membrane potential measured by TMRM assay in LTL331R-CL and NCI-H660 cells treated with DMSO or apilimod (1 µM) under low serum conditions for the indicated time points. TMRM fluorescence was normalized to the DMSO control at the matched time point for each cell line. FCCP (2.5 µM, 30 mins) was included as a positive control for mitochondrial depolarization and reduced TMRM signal. Data presented as mean ± SD. *P* values calculated using unpaired t-tests with Welch’s correction.

D. Relative mitochondrial superoxide production measured by MitoSOX assay in LTL331R-CL and NCI-H660 cells treated with DMSO or apilimod (1 µM) under low serum conditions for the indicated time points. MitoSOX fluorescence was normalized to the DMSO control at the matched time point for each cell line. Antimycin A (10 µM, 30 mins) was included as a positive control for mitochondrial superoxide production and increased MitoSOX signal. Data presented as mean ± SD. *P* values calculated using unpaired t-tests with Welch’s correction.

E. Western blot of indicated targets following DMSO or apilimod (1 µM) treatment for 24 hours under low serum conditions in LTL331R-CL CRISPRi sg*NC* and sg*EIF2AK1* cells.

**Figure S8. ER stress sensitizes NEPC cells to PIKfyve inhibitors. Related to Figure 4**.

A. Synergy matrix and percentage inhibition of viability between thapsigargin and ESK981 for 7 days in LTL331R-CL cells.

B. Synergy matrix and percentage inhibition of viability between thapsigargin and apilimod treatment for 7 days in NCI-H660 cells.

C. Dose response curve of ESK981 and apilimod under full serum or low serum conditions in NEPC cell lines to determine corresponding IC_50_. Data presented as mean ± SEM.

D. Relative cell viability of NCI-H660 (top) and LTL610-CL (bottom) cells treated with increasing concentrations of ESK981 under full serum or low serum conditions for 7 days. Data presented as mean ± SD*. P* values calculated using two-way ANOVA.

E. GSEA of hallmark hypoxia pathway and ATF4 target gene set in LTL545 NEPC tumors treated with apilimod compared with vehicle for 5 days.

F. Heatmap showing expression of leading-edge genes from the hallmark hypoxia pathway and the ATF4 target gene set in LTL545 tumors treated with apilimod compared with vehicle. Gene expression values were row-scaled and displayed as normalized Z scores.

**Figure S9. PIKfyve inhibition activates lipid biosynthesis. Related to Figure 5**.

A. GSEA of cholesterol homeostasis and fatty acid metabolism hallmark pathways comparing apilimod to DMSO treatment in LTL331R-CL cells.

B. GSEA of cholesterol homeostasis and fatty acid metabolism hallmark pathways comparing ESK981 to DMSO treatment in LTL331R-CL cells.

C. Volcano plot of DEGs in LTL331R-CL cells treated with Apilimod (left) or ESK981 (right) for 6 hours compared to DMSO. Genes involved in the cholesterol biosynthesis pathway are in red and fatty acid biosynthesis in blue. DEGs were defined using thresholds of log_2_FC greater than 1 or less than -1 and adjusted *p-*value below 0.05.

D. qPCR of the indicated genes encoding key enzymes involved in fatty acid biosynthesis and cholesterol biosynthesis post treatment with DMSO, apilimod (1 µM), and PIK5-33d (0.3 µM) for 6 hours in LTL331R-CL and NCI-H660 cells. Data presented as mean ± SD*. P* values calculated using ordinary one-way ANOVA.

E. Hallmark pathway enrichment analysis of LTL331R-CL cells post 24 hours of PIK5-33d (0.1 µM) treatment compared with DMSO by whole-cell proteomics.

F. Western blot of the indicated targets involved in fatty acid and cholesterol biosynthesis in NCI-H660 cells after apilimod (1 µM) or PIK5-33d (0.3 µM) treatment at the indicated time points.

G. Western blot of the indicated proteins involved in fatty acid and cholesterol biosynthesis after increasing concentrations of apilimod or PIK5-33d treatment for 48 hours in LTL331R-CL and NCI-H660 cells.

**Figure S10. PIKfyve inhibition-mediated lipid biosynthesis is regulated through SREBPs. Related to Figure 5**.

A. Western blot analysis of SREBP2 in LTL331R-CL and NCI-H660 cells following DMSO, apilimod (1 µM), or PIK5-33d (0.3 µM) treatment for 6 hours.

B. Western blot of PIKfyve, SREBP1, SREBP2, and LC3A/B in NEPC cells post DMSO, apilimod (1 µM), or PIK5-33d (0.3 µM) treatment for 1, 2, or 4 hours in indicated cells.

C. Western blot analysis of cytoplasmic precursor SREBP1 (P) and nuclear mature SREBP1 (M) in NCI-H660 cells treated with DMSO, apilimod (1 µM), or PIK5-33d (0.3 µM) for 6 hours (left). Quantification of relative protein levels of precursor SREBP1 (cytoplasmic) and mature SREBP1 (nuclear) (right). Data presented as mean ± SEM. *P* values calculated using one-way ANOVA.

D. Western blot analysis of cytoplasmic precursor SREBP1 (P) and nuclear mature SREBP1 (M) in LTL331R-CL cells treated with DMSO, apilimod (1 µM), or PIK5-33d (0.3 µM) for 2 or 4 hours (left). Quantification of relative protein levels of precursor SREBP1 (cytoplasmic) and mature SREBP1 (nuclear) (right). Data presented as mean ± SEM. *P* values calculated using one-way ANOVA.

E. qPCR analysis of the indicated genes encoding key enzymes involved in fatty acid and cholesterol biosynthesis post treatment with DMSO, apilimod (1 µM), fatostatin (20 µM), or the combination treatment for 6 hours in LTL331R-CL (top) and NCI-H660 cells (bottom). Data presented as mean ± SD*. P* values calculated using one-way ANOVA.

F. Western blot analysis of SCD1, FDFT1, and FASN in LTL331R-CL (left) and NCI-H660 cells (right) following treatment with DMSO, apilimod (1 µM), fatostatin (20 µM), or the combination treatment for 24 hours.

G. Forest plot of lipid class changes of whole-cell lipidomic profiles from NCI-H660 cells post apilimod and fatostatin co-treatment compared with apilimod at indicated time points.

H. Heatmap of the top apilimod-enriched lipid species whose abundance were reduced upon co-treatment with fatostatin, displayed as Z score normalized values.

**Figure S11. PIKfyve inhibition-induced lysosomal accumulation of cholesterol. Related to Figure 5**.

A. Volcano plots of DEPs post 24 hours of PIK5-33d (0.1 µM) treatment compared to DMSO in TMEM192-HA tagged LTL331R-CL, as determined by lyso-IP proteomics analysis.

B. Hallmark pathway enrichment analysis and volcano plots of DEPs in TMEM192-HA tagged LTL331R-CL cells post 24 hours of apilimod (1 µM) treatment compared to DMSO by lyso-IP proteomics.

C. Quantification of total intracellular cholesterol levels in NCI-H660 (top) and LTL331R-CL (bottom) cells following treatment with DMSO, apilimod (1 µM), or PIK5-33d (0.3 µM) in the presence or absence of simvastatin (10 µM). Cholesterol levels were normalized to DMSO control and are shown as relative fold change. Data presented as mean ± SEM. *P* values calculated using one-way ANOVA.

D. Representative confocal images of filipin and LAMP1 co-staining in LTL331R-CL cells following 24 hours treatment with U18666A (5 µg/mL), a cholesterol transporter inhibitor, which was used as a positive control to show cholesterol accumulation in lysosomes. Filipin was shown in green, LAMP1 in red, and co-localization in yellow. Arrows indicated co-localization between filipin and LAMP1.

E. Representative confocal images of filipin and LAMP1 co-staining in NCI-H660 cells following 24 hours treatment with DMSO, apilimod (1 µM), PIK5-33d (0.3 µM), or U18666A (5 µg/ml) treatment. Arrows indicated co-localization between filipin and LAMP1 (left). Quantification of filipin signal colocalizing with LAMP1 across treatment groups (right). *n* = 3 random fields. Experiments were repeated three times. Data presented as mean ± SEM*. P* values calculated using one-way ANOVA.

F. PCA of lyso-IP based lipidomic profiles following apilimod (1 µM) treatment at 0, 6, and 24 hours in TMEM192-HA tagged NCI-H660 cells.

G. Heatmap of the top lipid species altered upon apilimod treatment at 0, 6, and 24 hours in TMEM192-HA tagged NCI-H660 cells.

H. Western blot analysis of PIKfyve, SREBP1, and LC3A/B in multiple prostate cancer cell lines after DMSO, apilimod (1 µM), or PIK5-33d (0.3 µM) treatment for 6 hours.

I. qPCR analysis of the indicated genes encoding key enzymes involved in cholesterol biosynthesis post treatment with DMSO or apilimod (1 µM) for 6 hours in multiple prostate cancer cell lines. Data presented as mean ± SD*. P* values calculated using two-way ANOVA.

J. Quantification of total intracellular cholesterol levels in LNCaP (left) and VCaP (right) cells following treatment with DMSO, apilimod (1 µM), or PIK5-33d (0.3 µM) in the presence or absence of simvastatin (10 µM). Cholesterol levels were normalized to DMSO control and are shown as relative fold change. Data presented as mean ± SEM. *P* values calculated using one-way ANOVA.

**Figure S12. Pharmacological inhibition of lipogenesis synergizes with PIKfyve inhibition in NEPC. Related to Figure 6**.

A. Western blot of malonyl-lysine (Mal-K) post increasing concentrations of TVB-2640 for 24 hours in NEPC cells.

B. Western blot of p-ACC and ACC post increasing concentrations of ND-646 for 8 hours in NEPC cells.

C. Synergy matrix and percent inhibition of viability of ND-646 and apilimod, PIK5-33d, or ESK981 under full serum conditions for 7 days in LTL331R-CL cells.

D. Synergy matrix and percent inhibition of viability of TVB-2640 and ESK981 under full serum conditions for 7 days in LTL331R-CL cells.

E. Forest plot of lipid class changes of whole-cell lipidomic profiles from NCI-H660 cells post apilimod and TVB-2640 co-treatment compared with apilimod at the indicated time points.

F. Heatmap of the top apilimod-enriched lipid species whose abundance were reduced upon co-treatment with TVB-2640 of whole-cell lipidomic profiles from NCI-H660 cells at 24 hours, displayed as Z score normalized values.

G. Violin plots showing relative abundance (Z score-normalized) of triglycerides of whole-cell lipidomic profiles from NCI-H660 cells across the indicated treatment conditions at 24 hours. *Q* values calculated using Welch’s ANOVA, followed by pairwise multiple comparisons with Benjamini-Hochberg FDR correction.

**Figure S13. ER stress potentiates PIKfyve inhibition-mediated lipogenesis in NEPC. Related to Figure 6**.

A. RNA-seq analysis of low serum versus full serum in LTL331R-CL cells including hallmark pathway enrichment (left), volcano plot of DEGs (middle), and GSEA of cholesterol homeostasis hallmark pathway (right). DEGs were defined using thresholds of log_2_FC greater than 1 or less than -1 and adjusted *p-*value below 0.05.

B. Hallmark pathway enrichment analysis post DMSO, apilimod (1 µM), PIK5-33d (0.3 µM), or ESK981 (1 µM) treatment in LTL331R-CL cells under low serum conditions for 6 hours.

C. qPCR of *FASN* and *SCD1* mRNA levels post DMSO, apilimod (1 µM), or PIK5-33d (0.3 µM) treatment under full serum or low serum conditions for 6 hours. Data presented as mean ± SD*. P* values calculated using two-way ANOVA.

D. Western blot of SREBP1, SCD1, and ATF4 expression post DMSO, apilimod (1 µM), or PIK5-33d (0.3 µM) treatment under full serum and low serum conditions for 24 hours in LTL331R-CL cells.

E. qPCR analysis of *ATF4* in LTL331R-CL (left) and NCI-H660 (right) cells expressing CRISPRi *sgNC*, *sgATF4-1*, and *sgATF4-2*. Data presented as mean ± SD. *P* values calculated using one-way ANOVA.

F. Relative cell viability over time in LTL331R-CL (left) and NCI-H660 cells (right) expressing CRISPRi *sgNC*, *sgATF4-1*, and *sgATF4-2*. Data presented as mean ± SEM. *P* values calculated using two-way ANOVA.

G. Western blot analysis of ATF4, SREBP1, SREBP2, and LC3A/B post DMSO or apilimod (1 µM) treatment for 6 hours in LTL331R-CL cells expressing CRISPRi *sgNC*, *sgATF4-1*, and *sgATF4-2*.

H. Western blot analysis of cytoplasmic SREBP1 (P) and nuclear SREBP1 (M) in LTL331R-CL cells expressing CRISPRi *sgNC*, *sgATF4-1*, or *sgATF4-2* following treatment with DMSO or apilimod (1 µM) for 6 hours.

I. Western blot analysis of cytoplasmic SREBP1 (P) and nuclear SREBP1 (M) in NCI-H660 cells expressing CRISPRi *sgNC* or *sgATF4-1* following treatment with DMSO or apilimod (1 µM) for 6 hours.

J. qPCR analysis of *FASN*, *SCD1*, and *FDFT1* in LTL331R-CL cells expressing CRISPRi *sgNC*, *sgATF4-1*, and *sgATF4-2* following treatment with DMSO or apilimod (1 µM) for 6 hours. Data presented as mean ± SD. *P* values calculated using two-way ANOVA.

K. qPCR analysis of *SCAP* mRNA expression in LTL331R-CL cells expressing CRISPRi *sgNC*, *sgSCAP-1*, and *sgSCAP-2*. Data presented as mean ± SD. *P* values calculated using one-way ANOVA.

L. qPCR analysis of *SREBF1*, *FASN*, *SCD1*, *ATF3*, and *DDIT3* mRNA expression in LTL331R-CL cells expressing CRISPRi *sgNC* or *sgSREBF1* following treatment with DMSO or apilimod (1 µM) for 6 hours under low serum conditions. Data presented as mean ± SD. *P* values calculated using two-way ANOVA.

M. Western blot analysis of c-PARP and SCD1 post DMSO or apilimod (1 µM) treatment for 24 hours in LTL331R-CL cells expressing CRISPRi *sgNC*, *sgATF4-1*, and *sgATF4-2*.

N. Relative cell viability in LTL331R-CL cells expressing CRISPRi *sgNC*, *sgATF4-1*, and *sgATF4-2* following treatment with DMSO or apilimod (1 µM) under full serum or low serum conditions for 7 days. Data presented as mean ± SEM. *P* values calculated using two-way ANOVA.

O. Relative cell viability in NCI-H660 cells expressing CRISPRi *sgNC*, *sgATF4-1*, and *sgATF4-2* following treatment with DMSO or apilimod (1 µM) under low serum conditions for 7 days. Data presented as mean ± SEM. *P* values calculated using two-way ANOVA.

**Figure S14. Lipogenesis blockade amplifies PIKfyve inhibition-induced ER stress and reveals synthetic vulnerability in NEPC cells. Related to Figure 6**.

A. Autophagic flux measured by DU145 and PC3 GFP-LC3-RFP-ÄLC3 reporter cells treated with DMSO, TVB-2640, and ND-646 under full serum or low serum conditions for 48 hours. Data presented as mean ± SD*. P* values calculated using ordinary one-way ANOVA compared with DMSO group.

B. Western blot of autophagy markers (LC3A/B, p62) and lipogenesis markers (FASN, SCD1) following DMSO, apilimod (1 µM), and PIK5-33d (0.3 µM) treatment, with or without TVB-2640 (1 µM) or ND-646 (1 µM), under full serum or low serum conditions in LTL331R-CL cells for 24 hours.

C. qPCR of *ATF3* mRNA expression after DMSO, apilimod (1 µM), or PIK5-33d (0.3 µM) treatment, with or without TVB-2640 (1 µM) or ND-646 (1 µM), under full serum or low serum conditions in LTL331R-CL cells for 6 hours. Data presented as mean ± SD*. P* values calculated using two-way ANOVA.

D. Western blot of ATF4 following DMSO or the combination of apilimod (1 µM) and TVB-2640 (1 µM) or ND-646 (1 µM), with or without HRI inhibitors (hemin, 2.5 µM, 5 µM), PERK inhibitor (GSK26561567, 2 µM), ATF6α inhibitor (Ceapin A7, 10 µM), IRE1 inhibitor (4µ8C, 2 µM), or ISR inhibitor (ISRIB, 1 µM), under low serum conditions in LTL331R-CL cells for 24 hours.

E. qPCR of *EIF2AK1/HRI* and ER stress-associated transcripts in LTL331R-CL CRISPRi cells. *EIF2AK1* expression was measured in sg*NC* and *sgEIF2AK1* LTL331R-CL CRISPRi cells to confirm knockdown of *EIF2AK1*. *ATF4*, *ATF3*, and *DDIT3* expression was measured in *sgNC* and *sgEIF2AK1* cells treated with DMSO, apilimod (1 µM), TVB-2640 (1 µM), or the combination for 6 hours. Data presented as mean ± SD. *P* values calculated using two-tailed unpaired t-test for *EIF2AK1* knockdown and ordinary two-way ANOVA for treatment-response comparisons.

F. qPCR of IRE1á branch target genes (*EDEM1* and *XBP-1s*) and ATF6 branch target genes (*GRP94* and *ERP72*) mRNA expression after DMSO, apilimod (1 µM), and PIK5-33d (0.3 µM) treatment, with or without TVB-2640 (1 µM) or ND-646 (1 µM), under low serum conditions in LTL331R-CL cells for 6 hours. Data presented as mean ± SD*. P* values calculated using two-way ANOVA.

G. Relative cell viability of NCI-H660 cells treated with DMSO, apilimod (3 µM), and PIK5-33d (0.3 µM), with or without TVB-2640 (500 nM) and ND-646 (50 nM) under full serum and low serum conditions for 7 days. Data presented as mean ± SD*. P* values calculated using two-way ANOVA.

H. Relative cell viability of LTL331R-CL cells after DMSO, TVB-2640 (50 nM), apilimod (1 µM), or the combination of TVB-2640 and apilimod treatment with or without thapsigargin (2 nM) for 7 days. Data presented as mean ± SD*. P* values calculated using two-way ANOVA.

**Figure S15. Combined PIKfyve and lipogenesis inhibition promotes ER stress-associated apoptotic vulnerability in NEPC cells. Related to Figure 6**.

A. Western blot analysis of c-PARP, ATF3, CHOP, and LC3A/B after DMSO, apilimod (1 µM), and PIK5-33d (0.3 µM) treatment, with or without atorvastatin (10 µM), under low serum conditions in LTL331R-CL cells for 24 hours.

B. Western blot analysis of ATF4, CHOP, and c-PARP after DMSO, chloroquine (100 µM), and bafilomycin A1 (0.1 µM) treatment, with or without TVB-2640 (1 µM) or ND-646 (1 µM), under low serum conditions in LTL331R-CL cells for 24 hours.

C. qPCR of *ATF3* and *DDIT3* mRNA expression after DMSO, chloroquine (100 µM), and bafilomycin A1 (0.1 µM) treatment, with or without TVB-2640 (1 µM) or ND-646 (1 µM), under low serum conditions in LTL331R-CL cells for 6 hours. Data presented as mean ± SD*. P* values calculated using two-way ANOVA.

D. Western blot of indicated targets after DMSO, PIK5-33d (0.1 µM), TVB-2640 (1 µM), or ND-646 (1 µM) treatment, alone or in combination, with or without the pan-caspase inhibitor Q-VD-OPh (20 µM, pre-treated for 2 hours), in LTL331R-CL cells under low serum conditions for 24 hours.

E. Relative cell viability of LTL331R-CL (left) and NCI-H660 (right) cells treated with DMSO, apilimod (1 µM), TVB-2640 (1 µM), or the combination, in the presence of vehicle (DMSO), Q-VD-OPh (20 µM), ferrostatin-1 (10 µM), or necrostatin 1S (10 µM). Data normalized to the DMSO group within each arm and presented as mean ± SD. *P* values calculated using two-way ANOVA.

F. Western blot analysis of ATF4 and c-PARP post DMSO, apilimod (1 µM), TVB-2640 (1 µM), or the combination for 24 hours in LTL331R-CL cells expressing CRISPRi *sgNC*, *sgATF4-1*, and *sgATF4-2*.

G. Relative cell viability of LTL331R-CL (left) and NCI-H660 (right) cells expressing *sgNC*, *sgATF4-1*, or *sgATF4-2* following treatment with DMSO, apilimod (1 µM), TVB-2640 (1 µM), or the combination for 7 days. All data normalized to the *sgNC* DMSO group and presented as mean ± SD. *P* values calculated using two-way ANOVA.

**Figure S16. Synergy matrices for combined inhibition of PIKfyve and lipogenesis. Related to Figure 6**.

A. Synergy matrix and percent inhibition of viability between ND-646 and apilimod or PIK5-33d under low serum conditions for 7 days in LTL331R-CL cells.

B. Synergy matrix and percent inhibition of viability of TVB-2640 or ND-646 and apilimod under low serum conditions for 7 days in NCI-H660 cells.

C. Synergy matrix and percent inhibition of viability of TVB-2640 or ND-646 and apilimod under low serum conditions for 7 days in LTL610-CL cells.

D. Synergy matrix and percent inhibition of viability of HMGCR inhibitors (atorvastatin, simvastatin) or FDFT1 inhibitor (YM-53601) and apilimod under low serum conditions for 7 days in LTL331R-CL cells.

E. Synergy matrix and percent inhibition of viability of SQLE inhibitor (NB-598) or SCD1 inhibitor (MK-8245) and apilimod under low serum conditions for 7 days in LTL331R-CL cells.

F. Western blot analysis of HIF-1α, ATF4, and LC3A/B in LTL331R-CL cells post exposure to 21% O_2_, CoCl_2_ (12.5 µM), or 0.1% O_2_ for 48 hours.

G. Synergy matrix and percent inhibition of viability between TVB-2640 and apilimod under 21% O_2_, CoCl_2_ (12.5 µM), or 0.1% O_2_ conditions in low serum conditions for 7 days in LTL331R-CL cells.

H. Percent inhibition of viability of TVB-2640 and apilimod, supplemented with BSA or OA-BSA (200 µM), under low serum conditions for 7 days in NCI-H660 cells.

Synergy scores were calculated using the Bliss independence model, with scores above 10 indicating synergy.

**Figure S17. Therapeutic activity of combined PIKfyve and lipogenesis inhibition in NEPC models and additional cancer cell lines. Related to Figure 7**.

A. Heatmap showing expression of leading-edge genes corresponding to the GSEA plots of the hallmark unfolded protein response pathway and the ATF4 target gene set in LTL331R tumors treated with vehicle, TVB-2640, ESK981, and the combo treatment of ESK981 and TVB-2640 for five days. Gene expression values were row-scaled and displayed as normalized Z scores.

B. qPCR of *ATF4*, *ATF3*, *DDIT3*, and *SCD1* mRNA expression in LTL331R tumors treated with vehicle, TVB-2640, ESK981, and the combo treatment of ESK981 and TVB-2640 for three days. Data presented as mean ± SEM. *P* values calculated using Brown-Forsythe and Welch ANOVA with Dunnett’s T3 multiple-comparison test.

C. Representative images of LC3A/B IHC in LTL331R tumors treated with vehicle, TVB-2640, ESK981, and the combo treatment of ESK981 and TVB-2640 for three days.

D. Individual tumor growth trajectories (left), relative body weight monitoring (middle), and tumor weights at indicated time points (right) in the NEPC LTL331R PDX model treated with vehicle, TVB-2640, ESK981, or the combination of ESK981 and TVB-2640. Data presented as mean ± SEM. *P* values calculated using unpaired t test.

E. Individual tumor growth trajectories (left) and relative body weight monitoring (right) in the NEPC LTL610 PDX model treated with vehicle, TVB-2640, ESK981, or the combination of ESK981 + TVB-2640. Data presented as mean ± SEM.

F. Synergy matrix and percent inhibition of viability between TVB-2640 and apilimod or ESK981 for 5 days in 7940B cells under full serum conditions.

G. Synergy matrix and percent inhibition of viability between TVB-2640 and apilimod or ESK981 for 5 days in PDAC cell lines (HPAC, PANC-1, and SW1990) under full serum conditions.

H-I. Synergy matrix and percent inhibition of viability between TVB-2640 and apilimod for 5 days in PDAC (HPAF-II and MIA PaCa-2) (H) and gastric cancer (AGS) (I) cell lines under full serum conditions.

**Figure S18. *In vivo* investigation of PIKfyve inhibition-based combination therapies in additional cancer models. Related to Figure 7**.

A-B. Individual tumor growth trajectories, Kaplan-Meier analyses showing freedom from ≥4x (A) and ≥10x (B) tumor growth (%), tumor weights at indicated time points, and relative body weight monitoring in the pNET BON-1 CDX model (A) and PDAC 7940B syngeneic allograft model (B), respectively, treated with vehicle, TVB-2640, ESK981, or the combination of ESK981 and TVB-2640. Data presented as mean ± SEM. *P* values calculated using log-rank (Mantel-Cox) tests for Kaplan-Meier analyses; unpaired t test for tumor weight comparisons.

C-D. Tumor growth curves (left) and relative body weight monitoring (right) in the NEPC LTL545 (C) and LTL610 (D) PDX models treated with vehicle, cisplatin, ESK981, or the combination of ESK981 and cisplatin. Data presented as mean ± SEM. *P* values calculated using two-way ANOVA for tumor growth curves.

**Figure S19. Relative body weight, hematologic, and histopathologic assessment of major organs in CD-1 mice following PIKfyve inhibition-based combination treatments. Related to Figure 7**.

A. Mouse body weight monitoring in CD-1 mice treated with vehicle, ESK981, TVB-2640, ESK981 + TVB-2640, cisplatin, or ESK981 + cisplatin, shown as relative body weight over time during chronic (post-dose day 30, PD30) treatments. Sample sizes: *n* = 5 for each treatment group.

B. Complete blood count (CBC) analysis in CD-1 mice at PD5 (top) or PD30 (bottom) following treatment with vehicle, ESK981, TVB-2640, ESK981 + TVB-2640, cisplatin, or ESK981 + cisplatin. Parameters include total white blood cells (WBC), neutrophils (NEU), lymphocytes (LYM), monocytes (MONO), eosinophils (EOS), basophils (BAS), NEU%, LYM%, MONO%, EOS%, BAS%, red blood cells (RBC), hemoglobin (HGB), hematocrit (HCT), mean corpuscular volume (MCV), red cell distribution width (RDW%), mean corpuscular hemoglobin (MCH), and mean corpuscular hemoglobin concentration (MCHC), platelets (PLT), and mean platelet volume (MPV). Data presented as mean ± SEM. One mouse from the cisplatin group at PD30 was excluded from CBC analysis due to insufficient blood volume for testing.

C-D. Serum chemistry analysis assessing liver and kidney function in CD-1 mice at PD5 (C) and PD30 (D) following the indicated treatments, including alanine aminotransferase (ALT), aspartate aminotransferase (AST), creatinine (CREA), and blood urea nitrogen (BUN). Data presented as mean ± SEM. *P* values calculated using one-way ANOVA; *ns*, non-significant (all *p*-values > 0.05 comparing treatment groups with the vehicle group).

E. Representative H&E-stained sections of major organs, including heart, lung, liver, spleen, kidney, small intestine, prostate (anterior lobe), testis, and bone marrow, were collected from CD-1 mice at PD30 following treatment with vehicle, ESK981, TVB-2640, ESK981 + TVB-2640, cisplatin, or ESK981 + cisplatin.

## Supplementary Tables

**Table S1.** Short Tandem Repeat (STR) profiling of LTL610-CL and LTL331R-CL cells across passages. Related to Figure 1.

**Table S2**. Gene-level decomposition of the Hallmark unfolded protein response signature following apilimod treatment in LTL331R-CL cells under low-serum conditions. Related to Figure 4.

**Table S3**. Gene-level decomposition of the Hallmark unfolded protein response signature following ESK981 treatment in LTL331R-CL cells under low-serum conditions. Related to Figure S6.

**Table S4**. Gene-level decomposition of the Hallmark unfolded protein response signature following PIK5-33d treatment in LTL331R-CL cells under low-serum conditions. Related to Figure S6.

**Table S5**. Summary of branch-associated Hallmark unfolded protein response gene induction across PIKfyve-targeting agents. Related to Figure 4 and Figure S6.

**Table S6**. List of primers used in the study. Related to the STAR Methods.

**Table S7**. Gene list for NEPC signature, AR signature, ER stress, and ATF4 targets. Related to the STAR Methods.

**Table S8**. Whole-cell lipidomics dataset from NCI-H660 cells following indicated treatments for 24 hours. Related to Figure 5, Figure 6, Figure S10, Figure S12, and the STAR Methods.

**Table S9**. Lyso-IP based lipidomics dataset from NCI-H660 cells following apilimod treatment for indicated time points. Related to Figure 5, Figure S11, and the STAR Methods.

**Table S10**. Quantified proteins identified by whole-cell proteomics following PIK5-33d treatment in LTL331R-CL cells compared to DMSO. Related to Figure 5, Figure S9, and the STAR Methods.

**Table S11**. Quantified proteins identified by lyso-IP based proteomics following PIK5-33d treatment in LTL331R-CL cells compared to DMSO. Related to Figure 5, Figure S11, and the STAR Methods.

**Table S12**. Quantified proteins identified by lyso-IP based proteomics following apilimod treatment in LTL331R-CL cells compared to DMSO. Related to Figure S11 and the STAR Methods.

**Table S13**. Top metabolites correlated with PRISM drug response to PIK5-33d and ESK981. Related to Figure 6 and the STAR Methods.

