## Supplemental Figures S1 - S19 for "A Stress-Adaptive Lipid Kinase Axis Defines Metabolic Vulnerabilities in Neuroendocrine Prostate Cancer"

### Figure S1. Related to Figure 1.

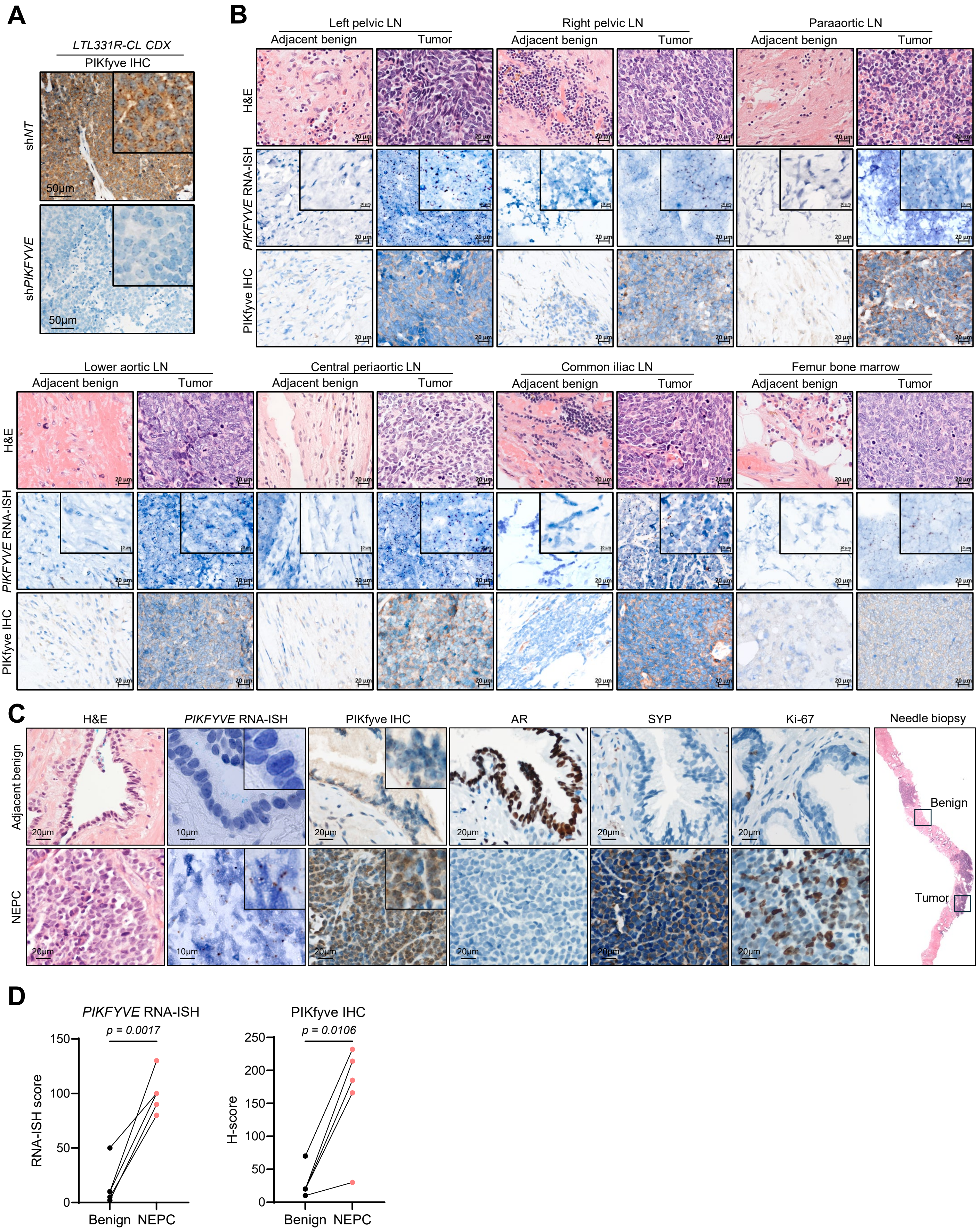

### Figure S2. Related to Figure 1.

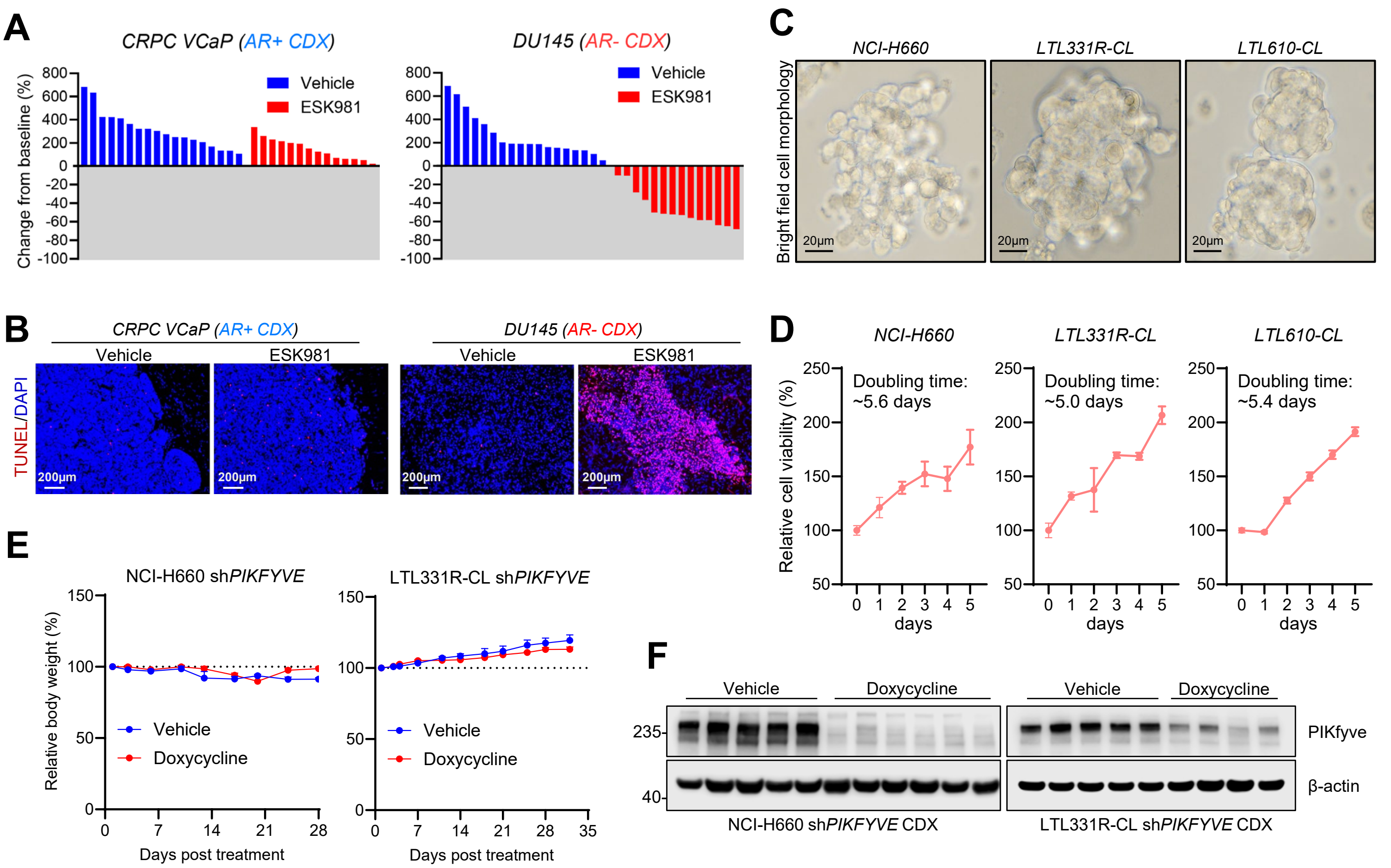

### Figure S3. Related to Figure 2.

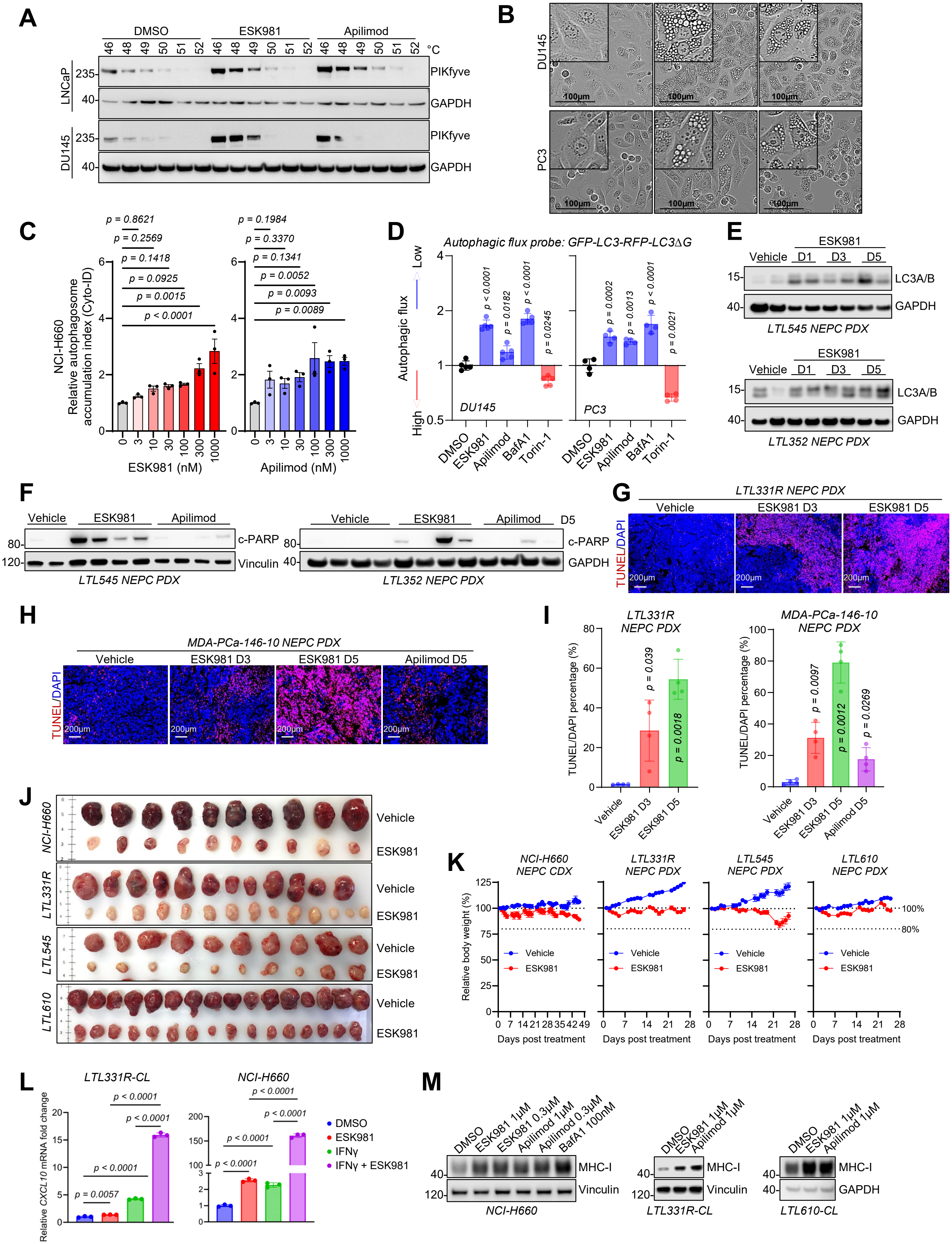

### Figure S4. Related to Figure 3.

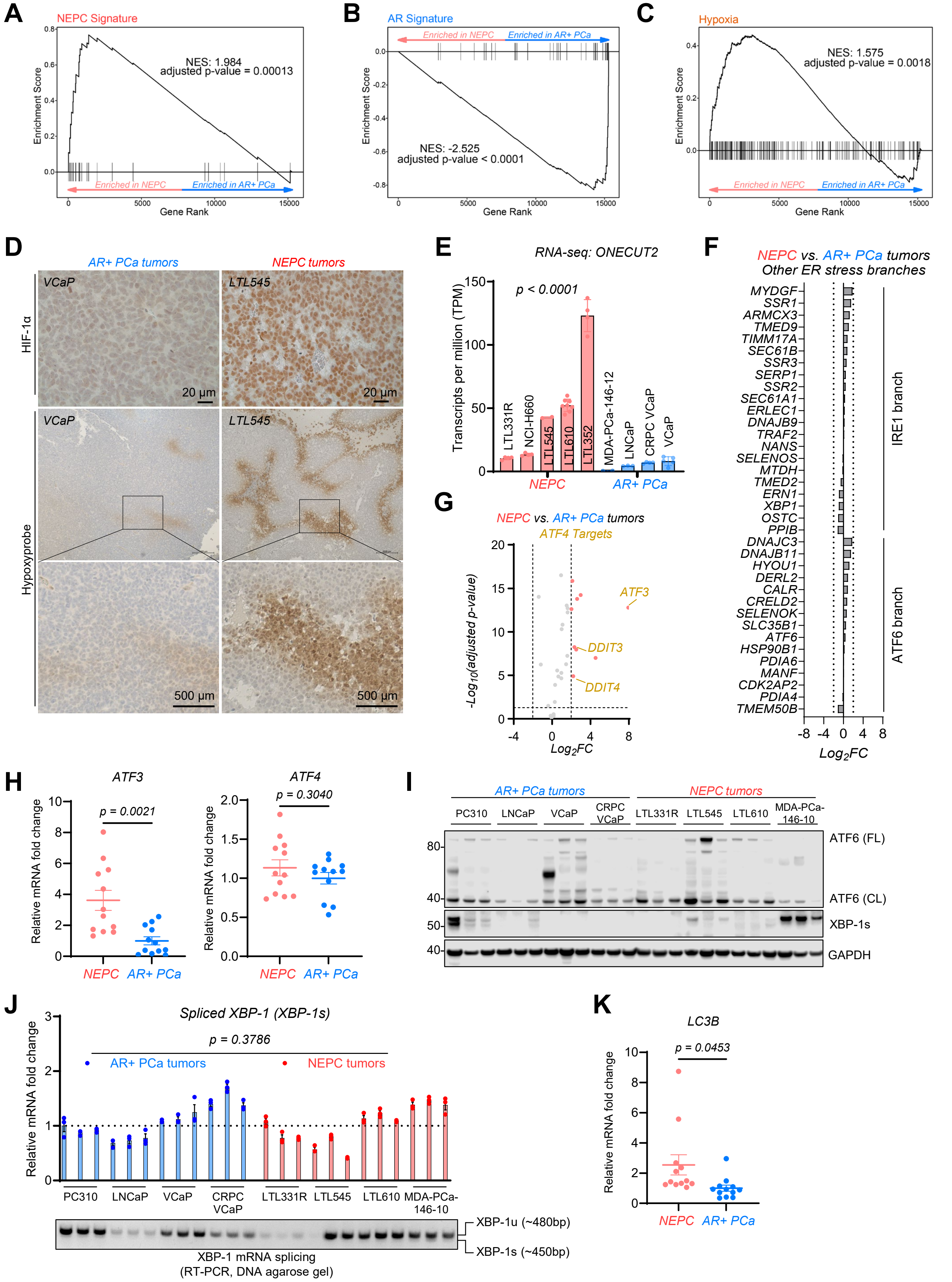

### Figure S5. Related to Figure 4.

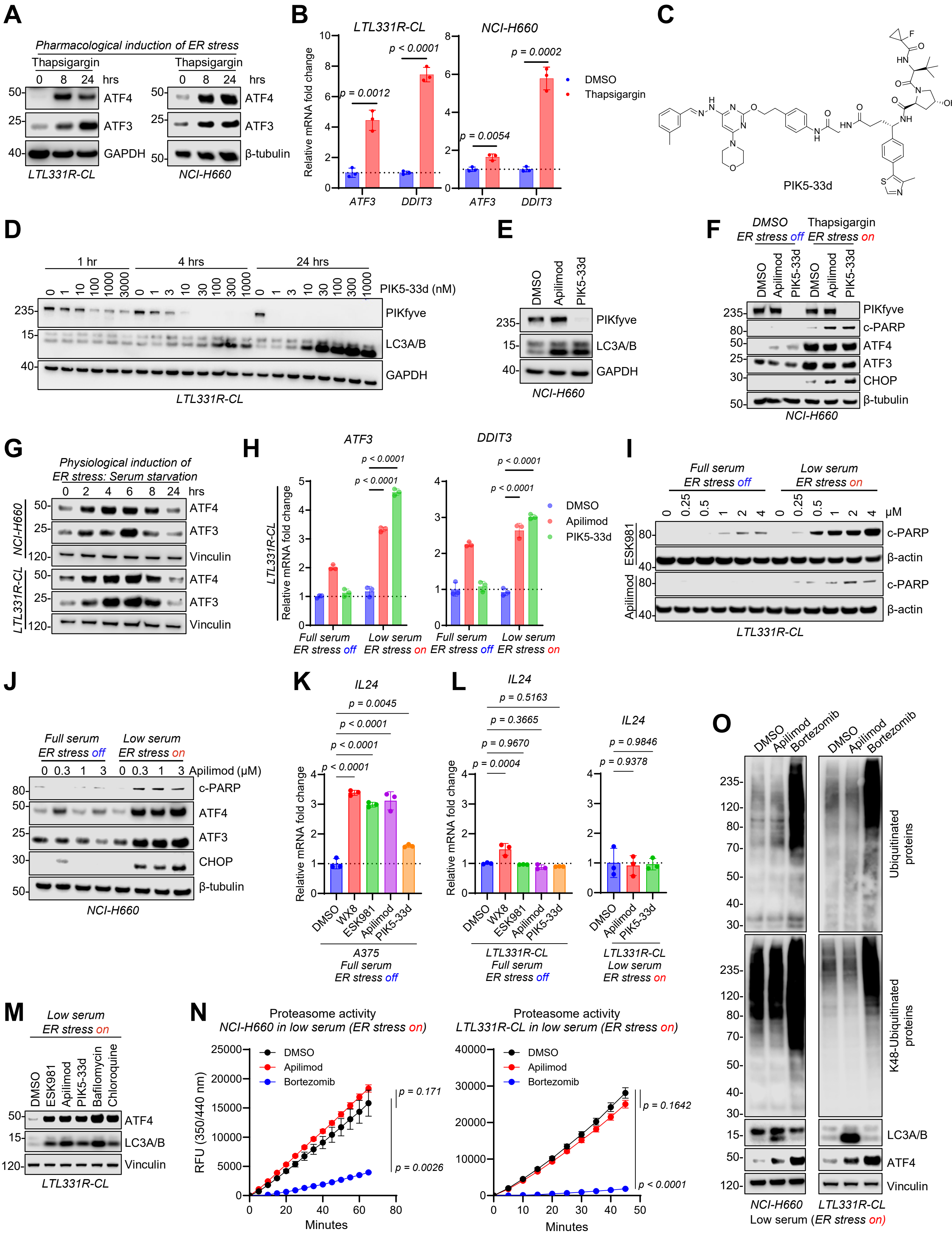

### Figure S6. Related to Figure 4.

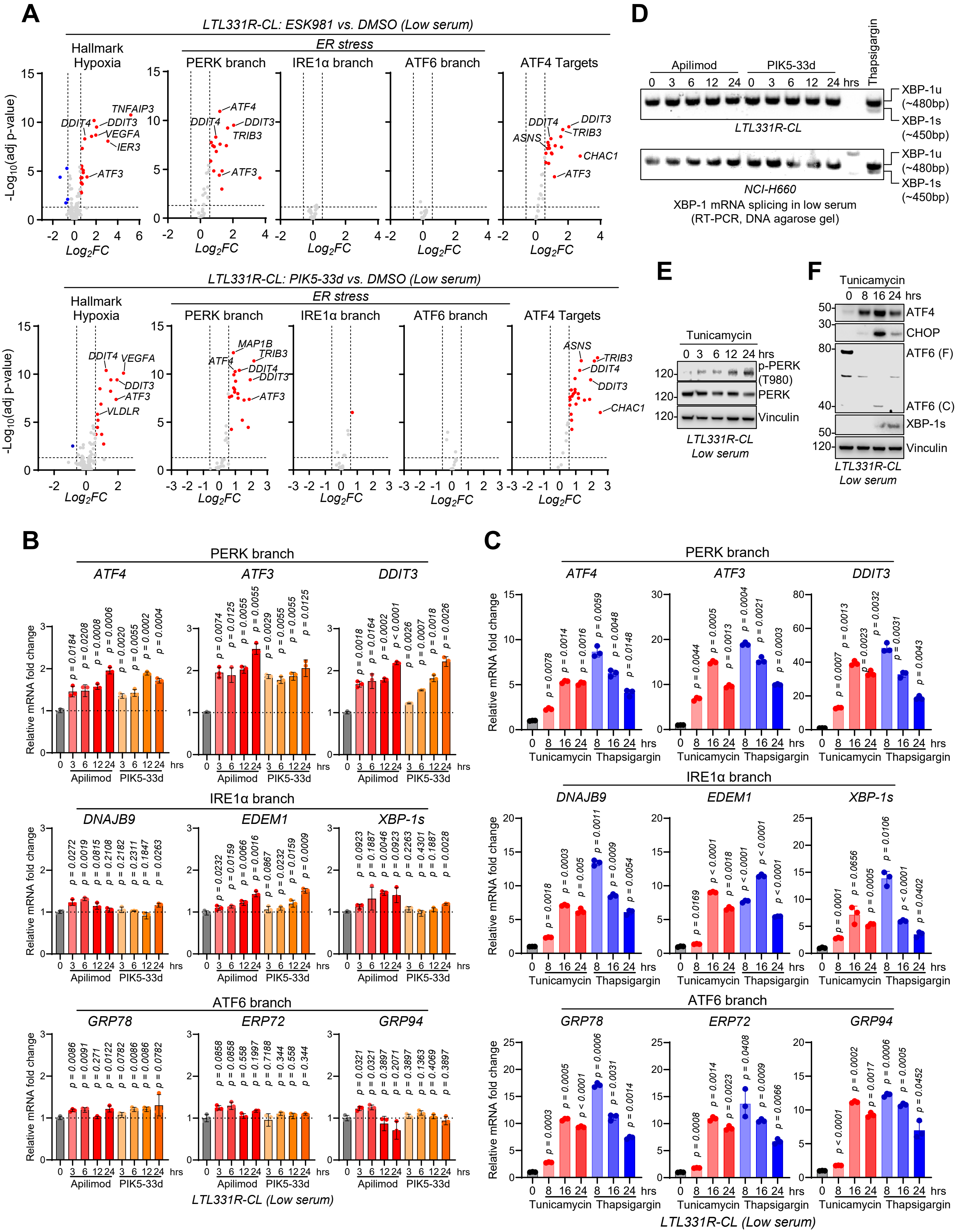

### Figure S7. Related to Figure 4.

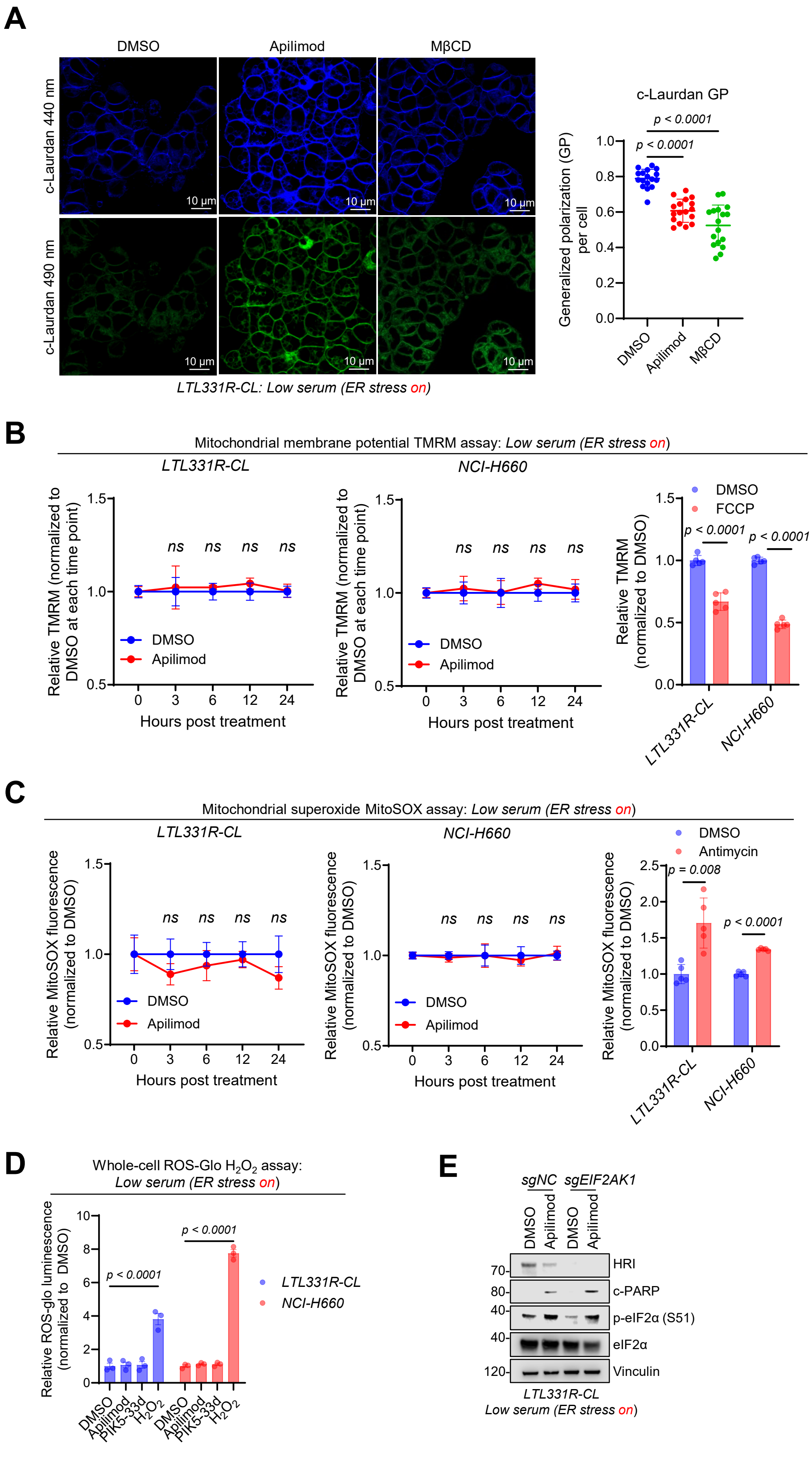

### Figure S8. Related to Figure 4.

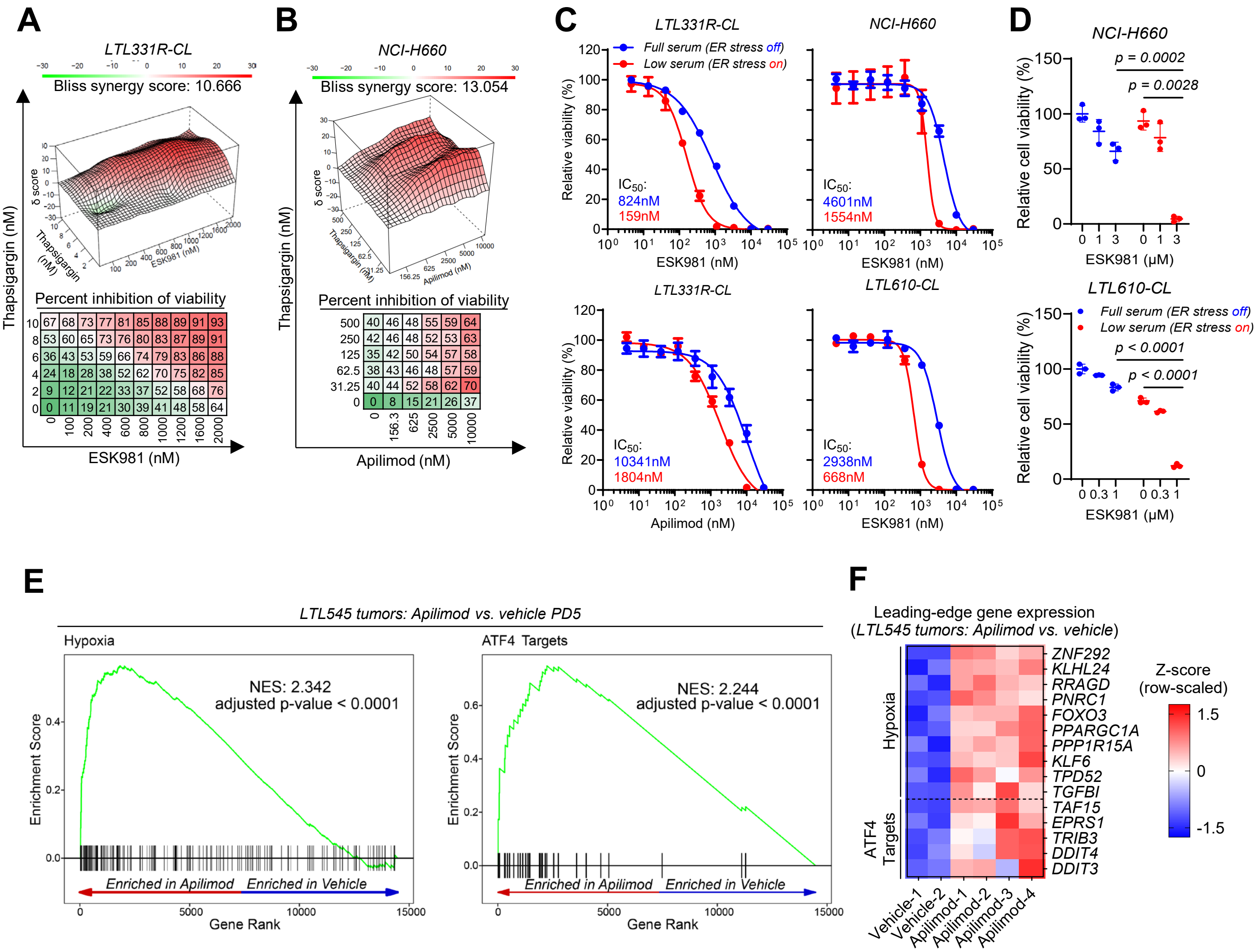

### Figure S9. Related to Figure 5.

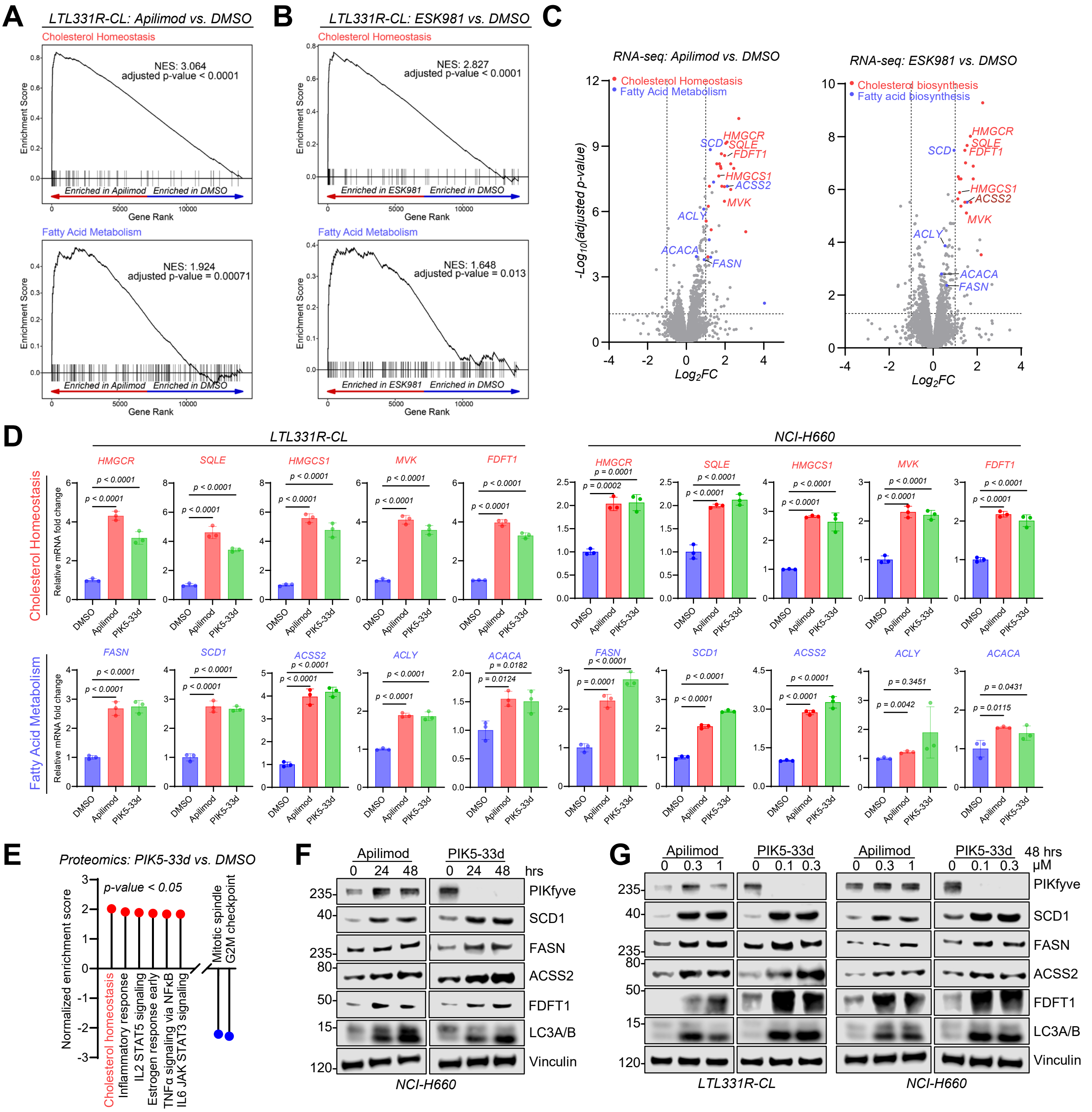

### Figure S10. Related to Figure 5.

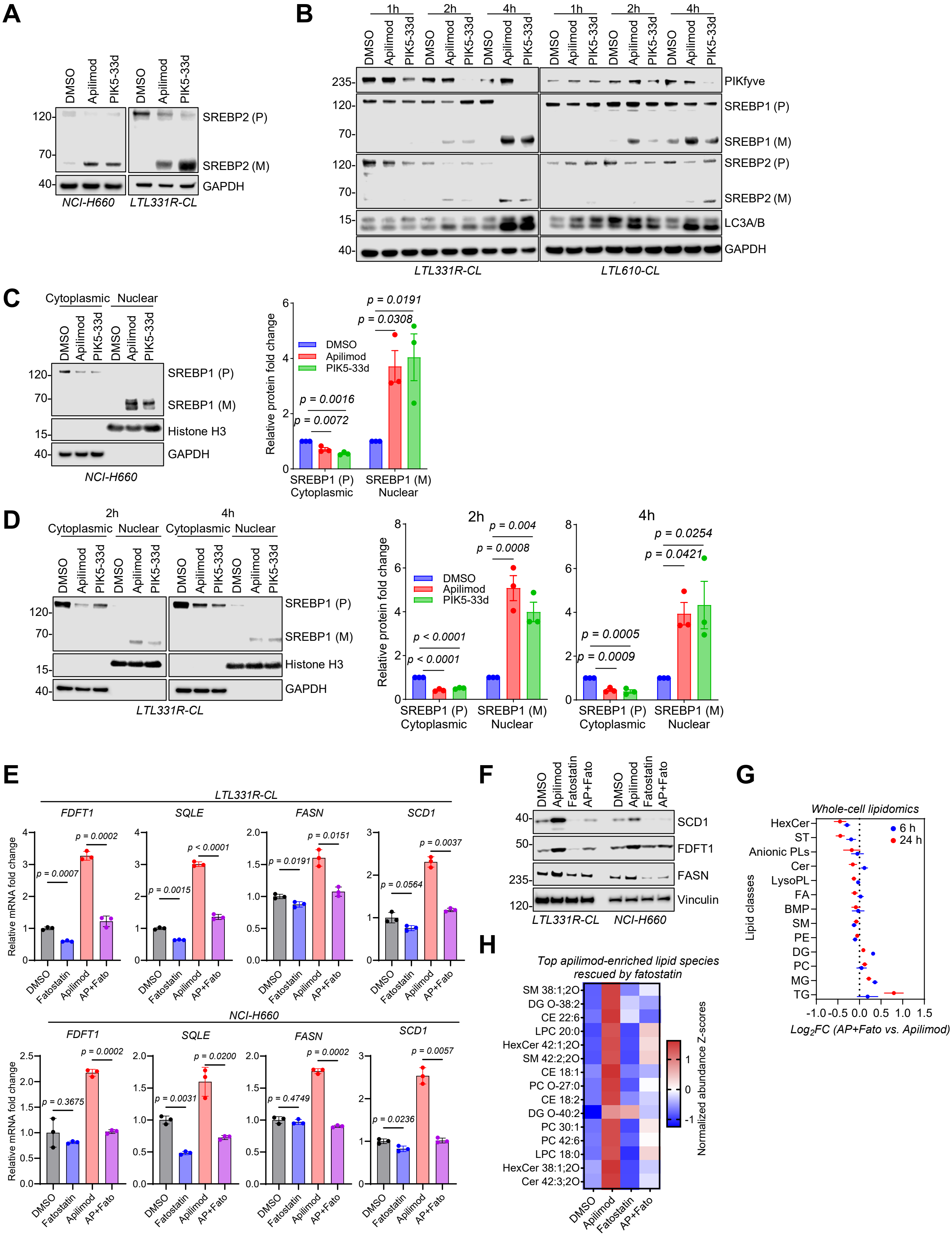

### Figure S11. Related to Figure 5.

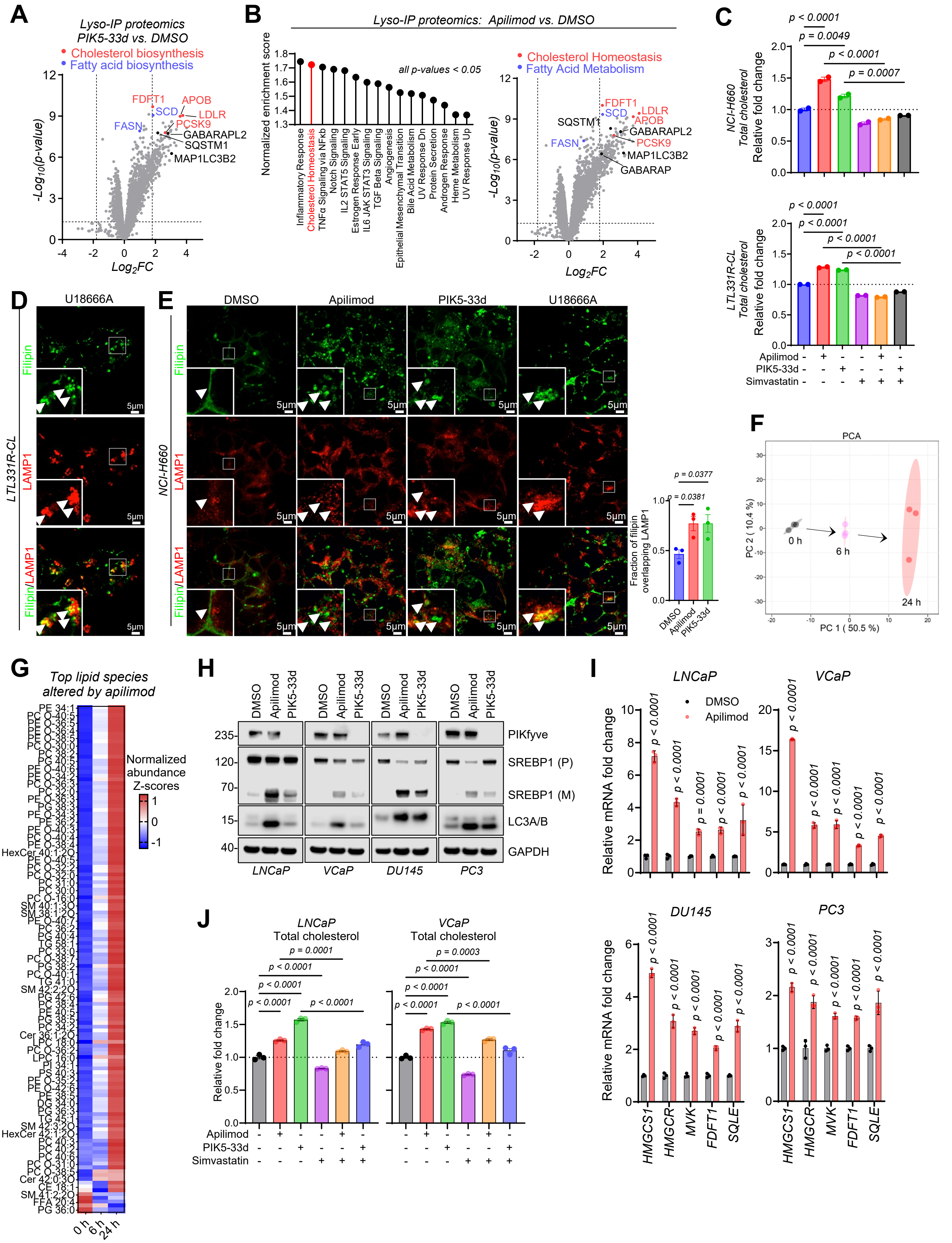

### Figure S12. Related to Figure 6.

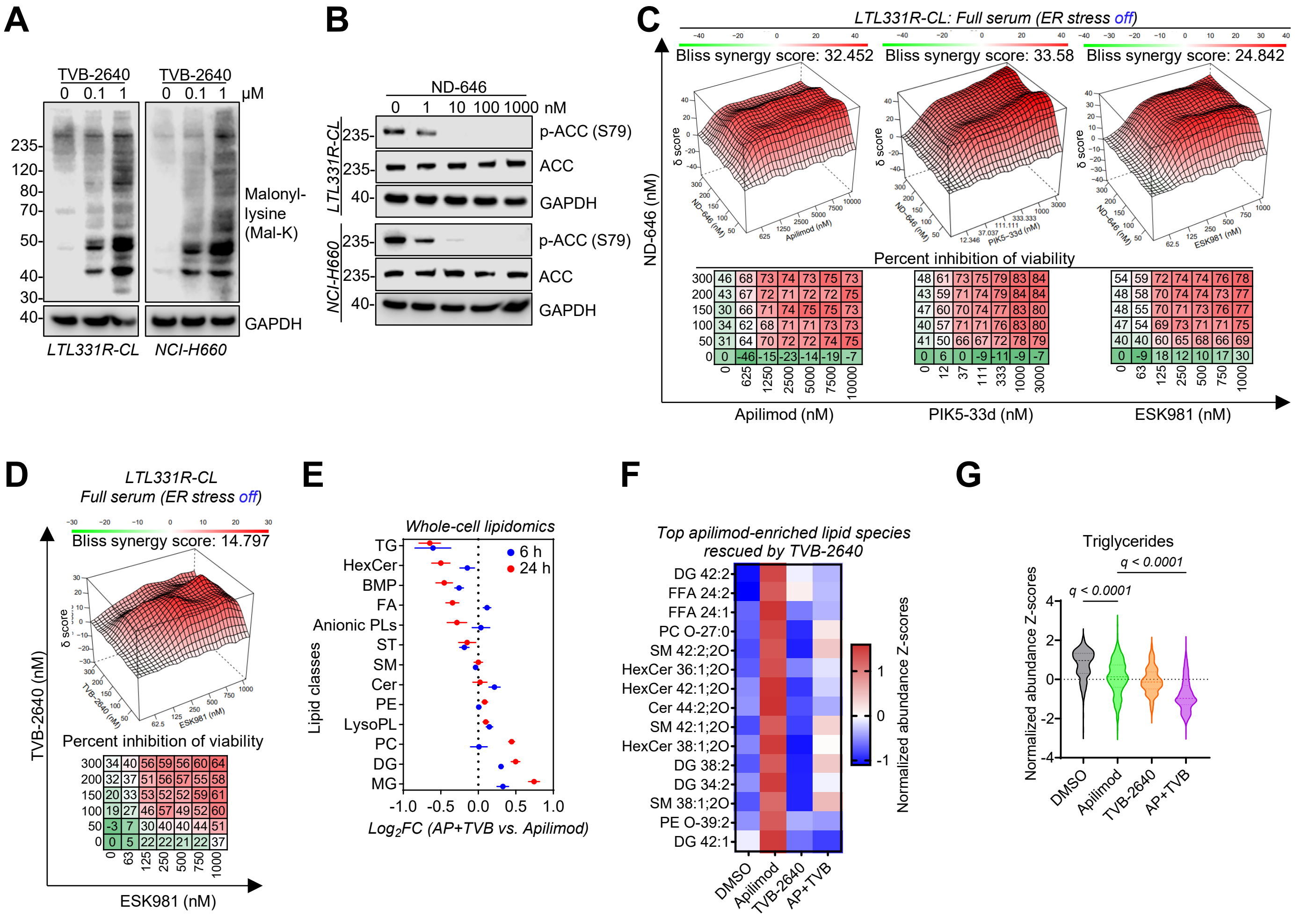

### Figure S13. Related to Figure 6.

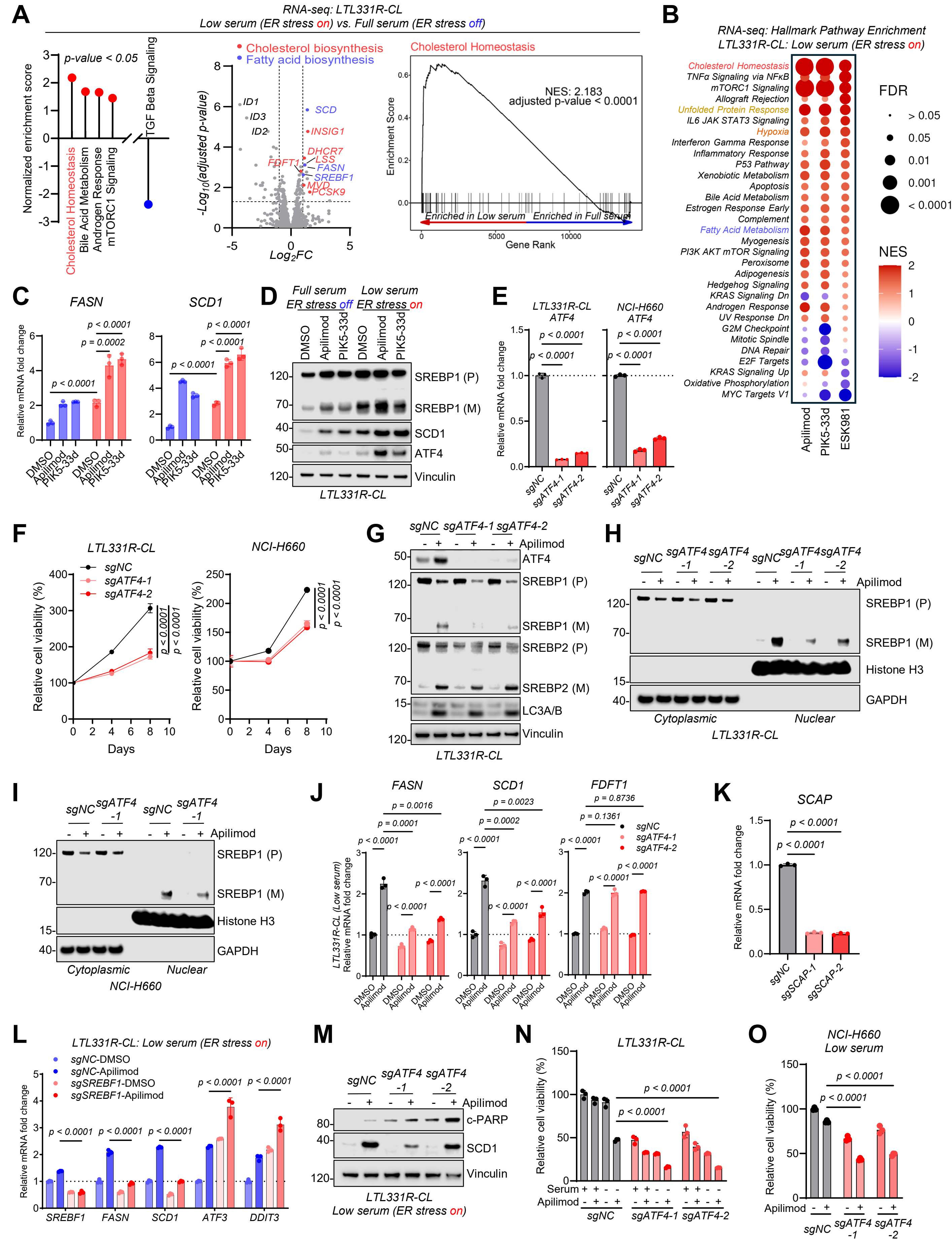

### Figure S14. Related to Figure 6.

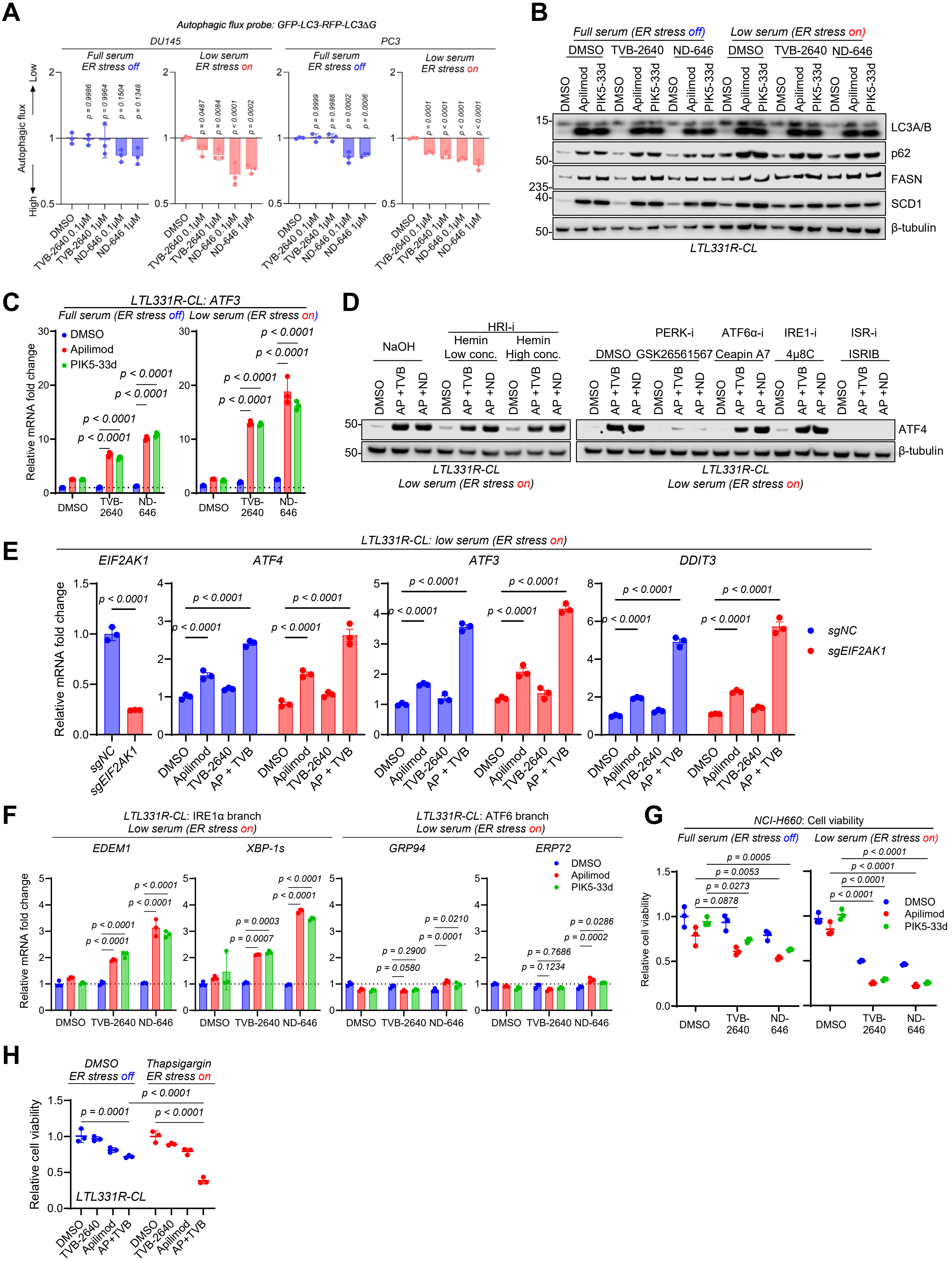

### Figure S15. Related to Figure 6.

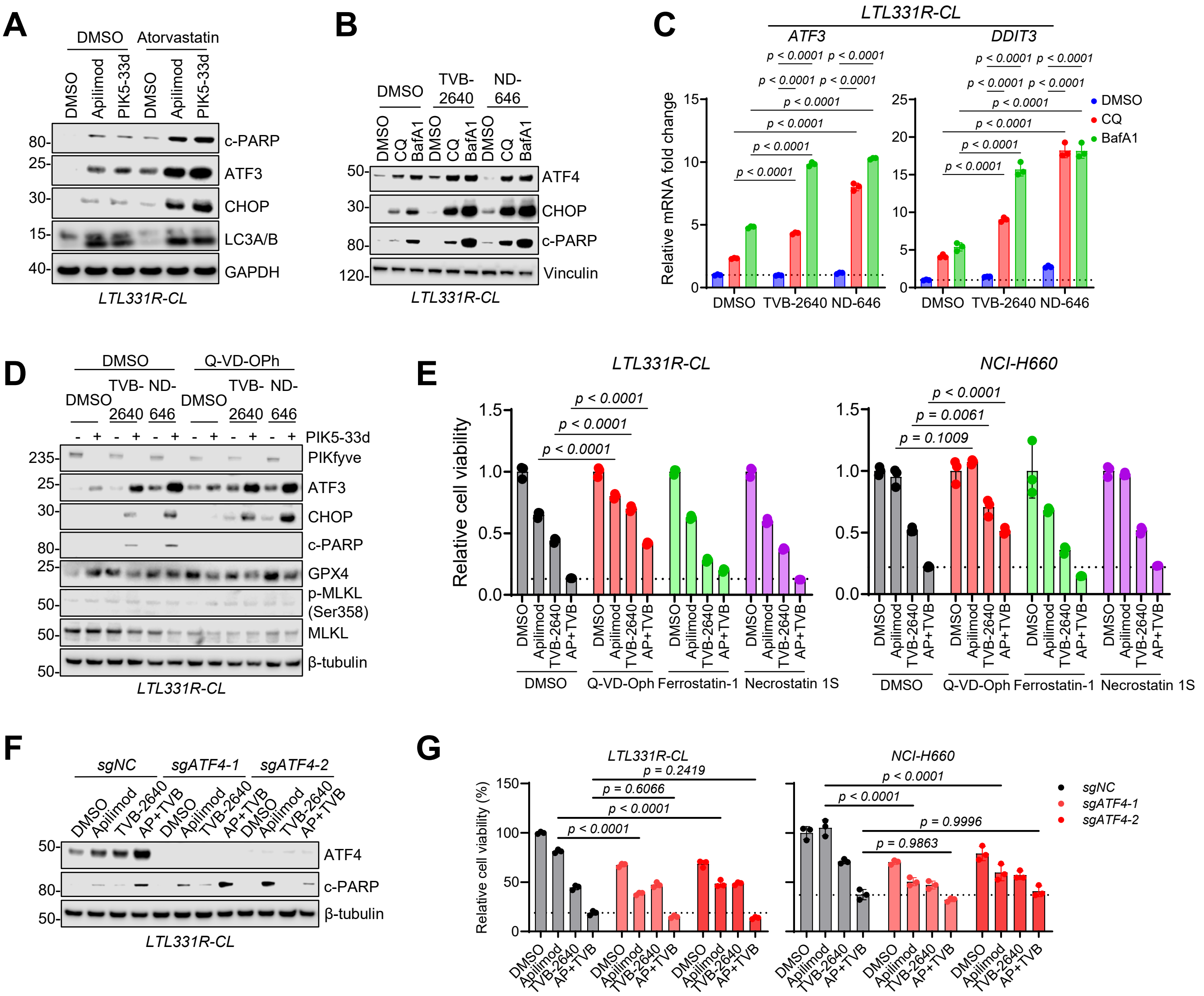

### Figure S16. Related to Figure 6.

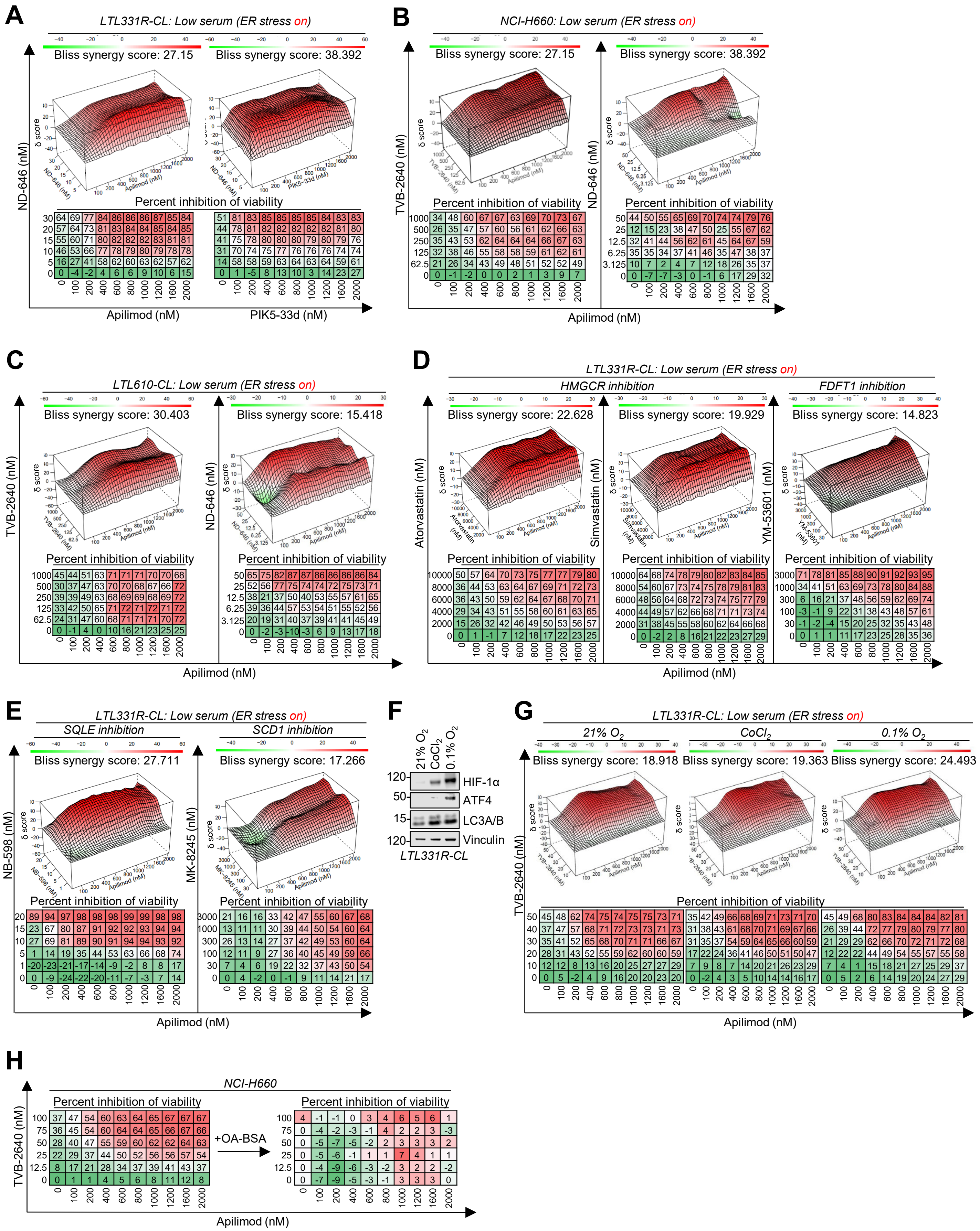

### Figure S17. Related to Figure 7.

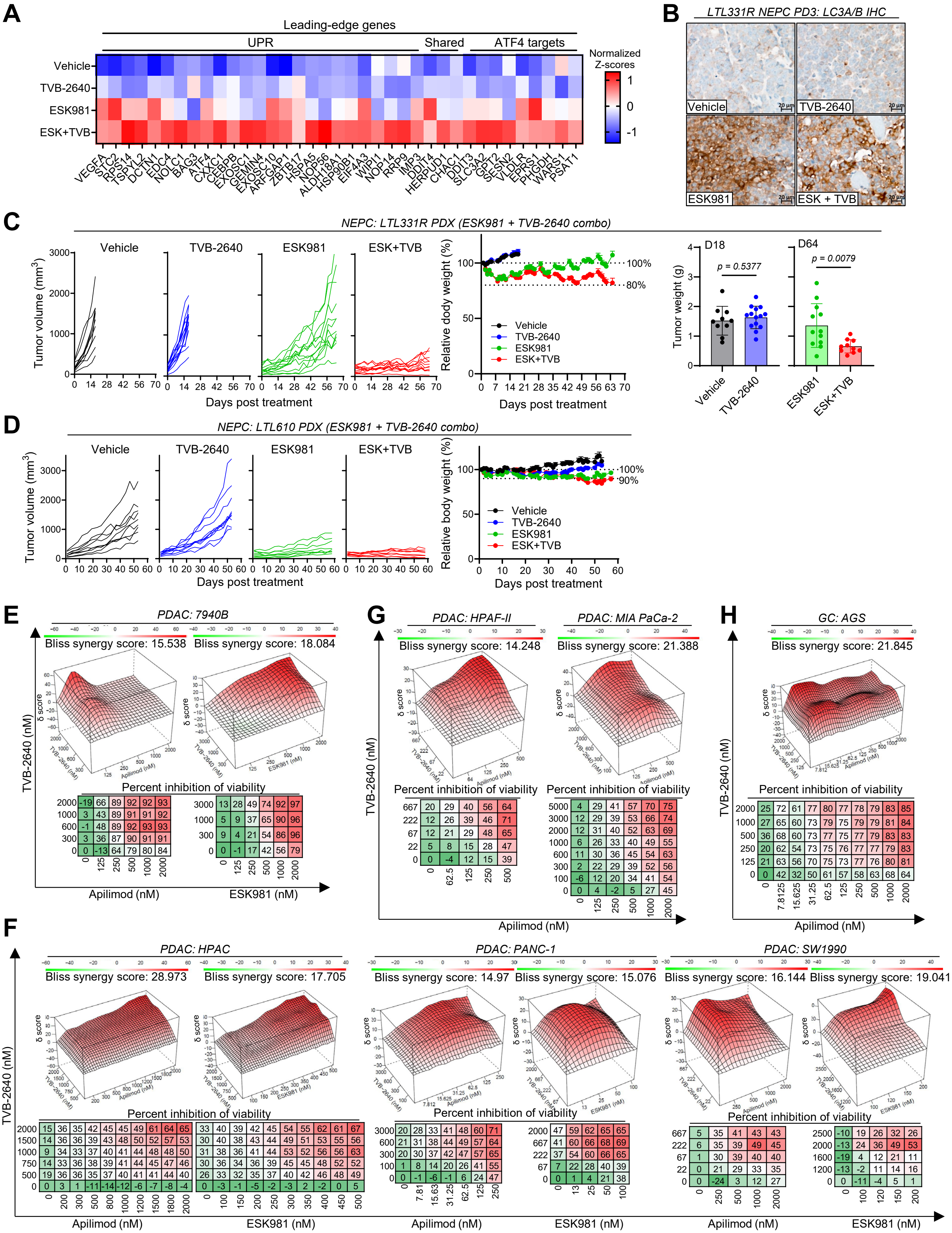

### Figure S18. Related to Figure 7.

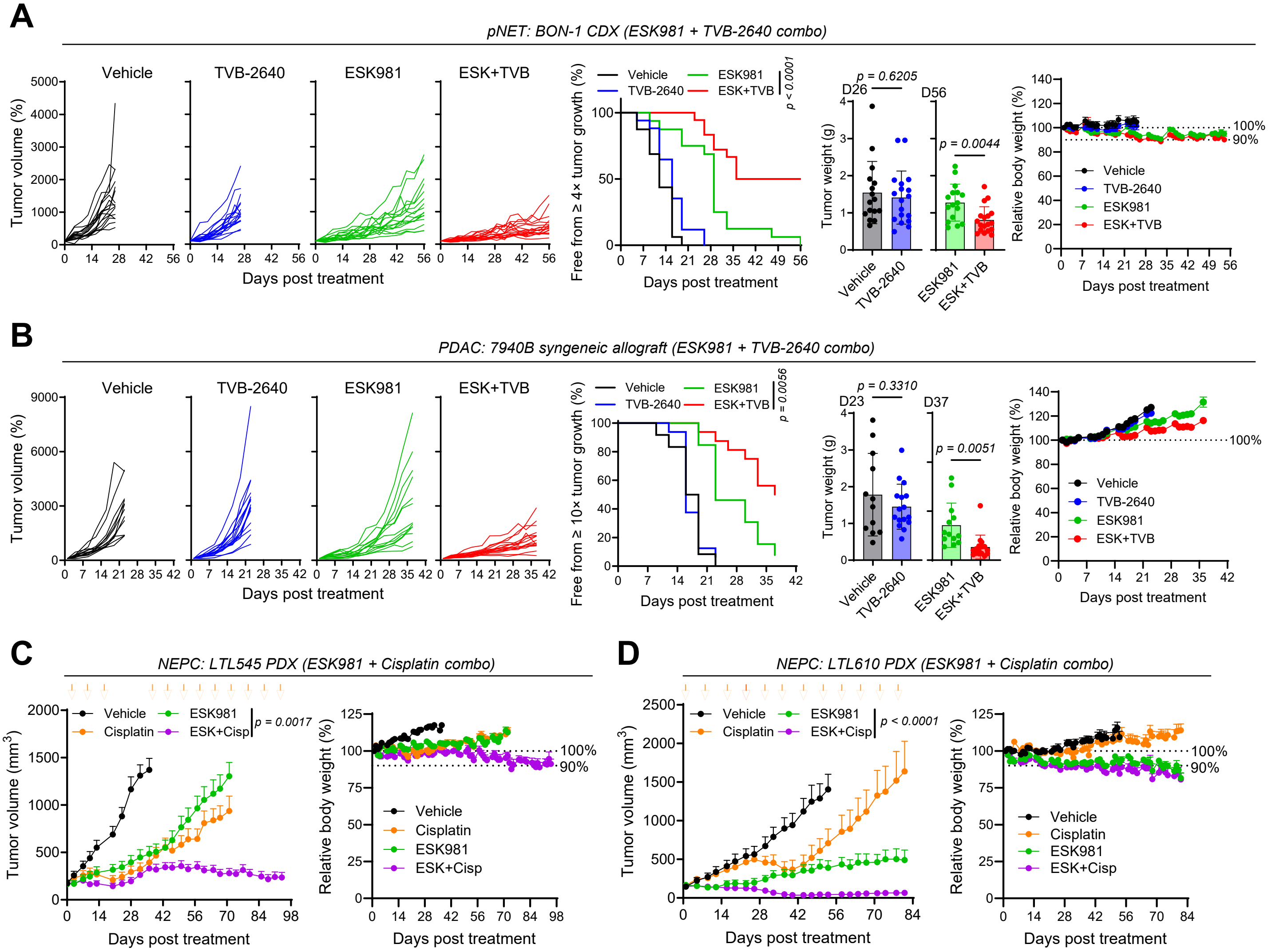

### Figure S19. Related to Figure 7.

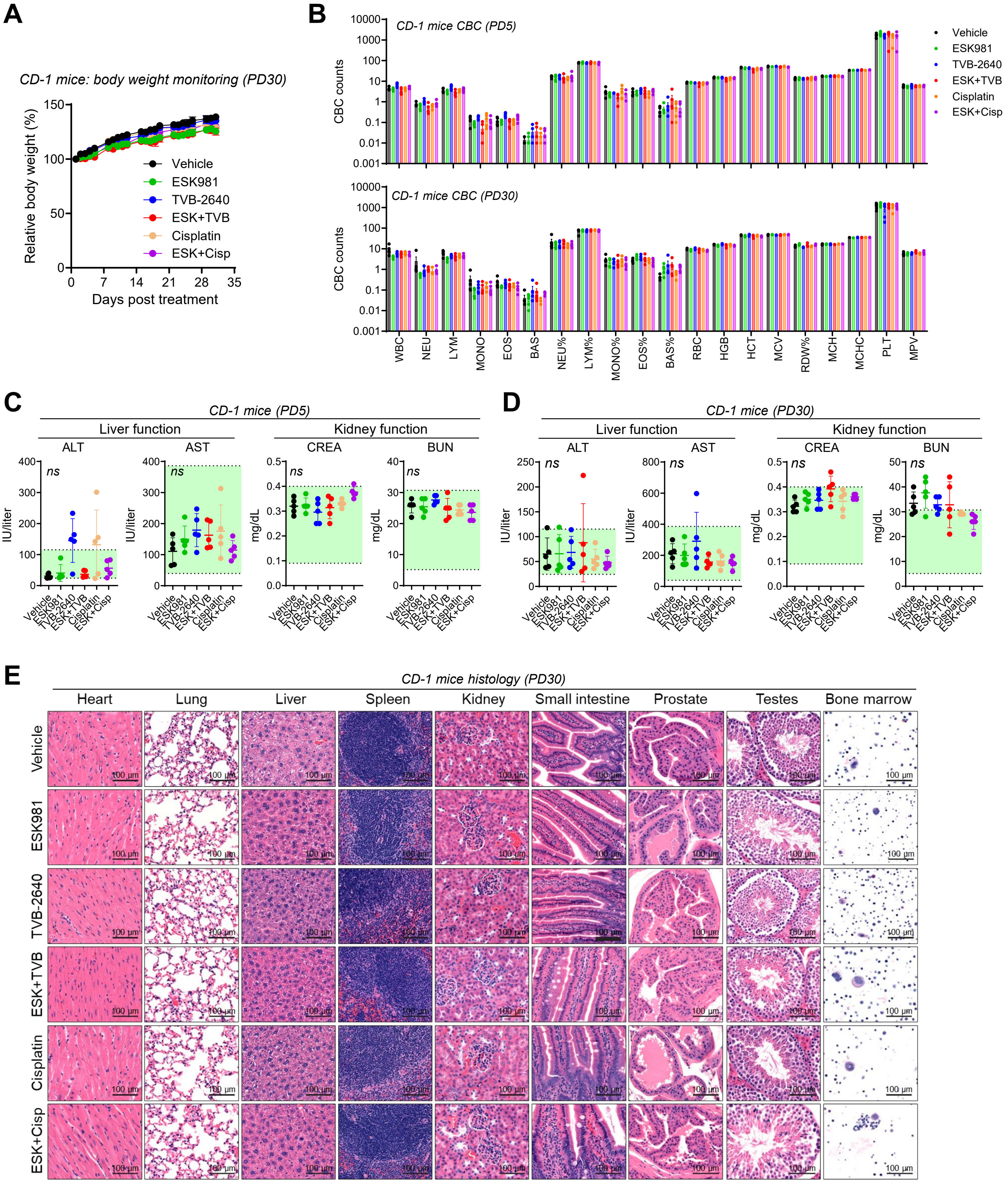
